# Divergent evolutionary trajectories of bryophytes and tracheophytes from a complex common ancestor of land plants

**DOI:** 10.1101/2021.10.28.466308

**Authors:** Brogan J. Harris, James W. Clark, Dominik Schrempf, Gergely J. Szöllősi, Philip C.J. Donoghue, Alistair M. Hetherington, Tom A. Williams

## Abstract

The origin of plants and their colonization of land resulted in the transformation of the terrestrial environment. Here we investigate the evolution of the land plants (embryophytes) and their two main lineages, the tracheophytes (vascular plants) and bryophytes (non-vascular plants). We used new fossil calibrations, relative lineage dating implied by horizontal gene transfer, and new phylogenomic methods for mapping gene family origins. Distinct rooting strategies resolve tracheophytes and bryophytes as monophyletic sister groups that diverged during the Cambrian, 515-494 Ma. The embryophyte stem is characterised by a burst of gene innovation, while bryophytes subsequently experienced a no less dramatic episode of reductive genome evolution in which they lost genes associated with the elaboration of vasculature and the stomatal complex. Overall, our analyses confirm that extant tracheophytes and bryophytes are both highly derived; as a result, understanding the origin of land plants requires tracing character evolution across the diversity of modern lineages.

## Introduction

The evolution of land plants (embryophytes) was a formative episode in Earth history, transforming the terrestrial landscape, atmosphere, and carbon cycle (Berry, Beerling and Franks, 2010; Pires and Dolan, 2012). Along with bacteria, algae, lichens, and fungi (Wellman and Strother, 2015), land plants were fundamental to the creation of the earliest terrestrial ecosystems and their subsequent diversification has resulted in more than 350,000 extant species (*The Plant List*). Embryophytes form a monophyletic group nested within fresh water streptophyte algae (de Vries and Archibald, 2018) and their move to land, while providing a new ecological niche, presented new challenges that required adaptation to water loss and growth against gravity (Raven, 2002). Early innovations that evolved in response to these challenges include a thick waxy cuticle, stomata, and means of transporting water from the roots up vertically growing stems (Pires and Dolan, 2012; de Vries and Archibald, 2018; Harrison and Morris, 2018; Donoghue *et al*., 2021). Modern land plants comprise two main lineages, vascular plants (tracheophytes) and non-vascular plants (bryophytes), that have responded to these evolutionary challenges in different ways.

The evolutionary origin of many gene families, including key regulators and transcription factors, has been shown to predate the colonisation of land (Wilhelmsson *et al*., 2017; Bowles, Bechtold and Paps, 2020). However, studies of gene family evolution within land plants have typically been restricted to individual gene families or sets of genes that encode single traits (Floyd and Bowman, 2007; Wang *et al*., 2010; Bowman *et al*., 2017; Gao *et al*., 2020; Harris *et al*., 2020; Radhakrishnan *et al*., 2020). A lack of genome-scale data from non-flowering plants has also hindered efforts to reconstruct patterns of genome and gene content evolution more broadly across land plants (Szövényi, Gunadi and Li, 2021), although large transcriptomic datasets have recently improved the situation (Leebens-Mack *et al*., 2019). Progress has also been made recently towards resolving the ambiguous phylogenetic relationships at the root of land plants (Cox *et al*., 2014; Puttick *et al*., 2018; Rensing, 2018; de Sousa *et al*., 2019; Leebens-Mack *et al*., 2019; Harris *et al*., 2020; Su *et al*., 2021). Further, the bryophyte fossil record has recently undergone a radical reinterpretation such that there are now many more potential calibrations to constrain the timing of early land plant evolution (Tomescu *et al*., 2018; Feldberg *et al*., 2021; Flores *et al*., 2021). Even in the absence of new fossils, novel methods have been developed to provide calibrations by using horizontal gene transfers to provide relative time constraints (Szöllősi *et al*., 2020).

Here we investigate the phylogeny, timescale, and gene content evolution of tracheophytes and bryophytes. We first infer a rooted phylogeny of land plants, using outgroup-free rooting methods and both concatenation and coalescent approaches. We then estimate an updated timescale of land plant evolution incorporating densely sampled fossil calibrations reflecting an up-to-date view of the fossil record. We extend this analysis to exploit the dating signal in gene transfer events and better calibrate the timescale of hornwort evolution, a poorly-constrained region of the land plant tree. The resulting dated phylogeny allows the reconstruction of gene content evolution in bryophytes and tracheophytes, and reveals how key genes, pathways and genomes have diverged since the common ancestor of land plants.

## Methods

### Sequence Data

The amino acid sequence dataset is comprised of 177 species, consisting of 23 algae and 154 plants **(Supplementary Table 1).** Sequence data was from published transcriptomes (Matasci *et al*., 2014; Leebens-Mack *et al*., 2019), or whole genome sequences from the NCBI repository (Federhen, 2012). A second dataset containing 30 whole genomes, consisting of 25 plants and 5 algae was also constructed (**Supplementary Table 2)**. Completeness of each genome/transcriptome was assessed using the BUSCO algorithm (Simão *et al*., 2015) with completeness measured as the percentage of present BUSCO genes (**Supplementary Tables 1; Supplementary Fig. 1)**.

### Software

All custom python scripts used in this analysis are available at – https://github.com/ak-andromeda/ALE_methods/. Software usage is described in the README document, along with a demonstration dataset.

### Orthologue inference

Orthologous gene families were inferred with OrthoFinder (Emms and Kelly, 2019); no universally present single-copy orthologous gene families were recovered **(Supplementary Fig. 1).** We therefore used a custom Python program (*prem3.py)* was used to systematically compute low-copy number orthologous gene families and from these identify single-copy gene families suitable for phylogenomics (**Supplementary Methods; Supplementary Fig. 2**). This approach yielded 160 single-copy gene families from 114016 orthogroups.

### Phylogenetics

#### Supermatrices

160 single copy gene families were aligned using MAFFT (Katoh and Toh, 2008), and poorly aligning sites were identified and removed with BMGE using the BLOSUM30 matrix (Criscuolo and Gribaldo, 2010). For ML analyses, we used the best-fitting substitution model as selected by BIC (LG+C60+G4+F) in IQ-TREE (V.1.6.12) (Nguyen *et al*., 2015); Bayesian analyses were performed under the CAT+GTR+G4 model in PhyloBayes version 2.3 (Blanquart and Lartillot, 2008; Lartillot *et al*., 2013). These models accommodate site-specific amino acid compositions via a fixed number of empirical profiles (C60) or an infinite mixture of profiles (CAT) (Lartillot and Philippe, 2004; Wang, Minh, Susko, 2014).

#### Supertrees

Individual maximum likelihood gene trees were inferred for each of the 160 single copy gene family in IQ-TREE (Nguyen *et al*., 2015), using the best-fitting model, selected individually for each gene using BIC. A supertree was then inferred using ASTRAL version 5.7.6 (Zhang *et al*., 2018).

### Divergence time estimation

Estimates of the origin of major lineages of land plants have proven robust to different phylogenetic hypotheses (Hedges *et al*., 2018; Morris *et al*., 2018), but not different interpretations of the fossil record (Hedges *et al*., 2018; Morris *et al*., 2018; Su *et al*., 2021). Recent studies of the timing of land plant evolution have argued that fossil calibrations should not exert undue influence over divergence time estimates (Su *et al*., 2021). However, in the absence of fossil calibrations, relaxed molecular clocks fail to distinguish rate and time and, as such, fossil calibrations are important across the tree to inform rate variation and in turn increase the accuracy of age estimates. Our approach thus sought to maximise the information in the fossil record and increase the sampling of fossil calibrations over previous studies (Morris *et al*., 2018; Su *et al*., 2021). Minimum age calibrations were defined based on the oldest unequivocal evidence of a lineage. Specifying a maximum age calibration can be controversial (Hedges *et al*., 2018; Su et al. 2021), yet maximum ages are always present, either as justified user-specified priors or incidentally as part of the joint time prior (Warnock, Yang and Donoghue, 2012; Warnock *et al*., 2015). On this basis, we defined our maxima following the principles defined in Parham *et al*., (2012) and fossil calibrations were defined as minimum and maximum age constraints, which in each case were modelled as uniform distributions between minima and maxima, with a 1% probability of either bound being exceeded (**Supplementary Methods**). We fixed the tree topology to that recovered by the Bayesian inference analysis and used the normal approximation method in MCMCtree (Yang, 2007), with branch lengths first estimated under the LG+G4 model in codeml (Yang, 2007). We divided the gene families into 4 partitions according to their rate, determined based on the maximum likelihood distance between *Arabidopsis thaliana* and *Ginkgo biloba.* We implemented a relaxed clock model (Uncorrelated; Independent Gamma Rates), where rates for each branch are treated as independent samples drawn from a lognormal distribution. The shape of the distribution is assigned a prior for the mean rate (μ) and for the variation among branches (σ), each modelled as a gamma distributed hyperprior. The gamma distribution for the mean rate was assigned a diffuse shape parameter of 2 and a scale parameter of 10, based on the pairwise distance between *Arabidopsis thaliana* and *Ginkgo biloba*, assuming a divergence time of 350 Ma (Morris *et al*., 2018). The rate variation parameter was assigned a shape parameter of 1 and a scale parameter of 10. The birth and death parameters were each set to 1, specifying a uniform kernel (Dos Reis *et al*., 2015). Four independent MCMC runs were performed, each running for 4 million generations to achieve convergence. Convergence was assessed in Tracer (Rambaut *et al*., 2018) by comparing posterior parameter estimates across all four runs and by ensuring effective sample sizes exceeded 200.

### Temporal constraint from a hornwort-to-fern horizontal gene transfer

Horizontal gene transfers (HGT) provide information about the order of nodes on a species phylogeny in time. Consequently, inferred HGT events can be used as relative node order constraints (Szöllősi *et al*., 2020). We used the horizontal transfer of the chimaeric neochrome photoreceptor (NEO) from hornworts to a derived fern lineage (Polypodiales; (Li *et al*., 2015)), as an additional source of data about divergence times in hornworts, a lineage that diverged early in plant evolution but is poorly represented in the fossil record. We inferred a new gene tree for NEO using the expanded sampling of lineages now available which confirmed the donor and recipient lineages originally reported (Li *et al*. 2015; **Supplementary Fig. 3**). Thus, we propose a relative calibration where the MRCA of the *Megaceros*-*Phaeoceros* clade is constrained to be older than Polypodiales.

This relative node order constraint was used together with the 66 fossil calibrations in a novel Bayesian inference software (mcmc-date, https://github.com/dschrempf/mcmc-date) to infer a species tree with branch lengths measured in absolute time. In contrast to MCMCTree, mcmc-date uses the posterior distribution of branch lengths estimated by Phylobayes, as described above, together with a multivariate normal distribution accounting for correlations between branches, to approximate the phylogenetic likelihood. Further, an exponential hyper-prior with mean 1.0 was used for the birth and death rates, as well as for the mean and variance of the gamma prior of the branch rates. A tailored set of random-walk proposals executed in random order per iteration, and the Metropolis-coupled Markov chain Monte Carlo (MC3) algorithm (Geyer, 1991) with four parallel chains, resulted in near-independence of consecutive samples. After a burn-in of approximately 5000 iterations, 15000 iterations were performed. All inferred parameters and node ages have effective samples sizes above 8000 as calculated by Tracer. Subsequently, the relative node dating analysis and the partitioned molecular clock analysis were combined by using the posterior distributions for the divergence times within hornworts from the relative node dating as a prior for the partitioned analysis in MCMCtree.

### Gene tree-species tree reconciliation

Modelling of gene duplication, transfer, and loss (DTL) with Amalgamated Likelihood Estimation (ALE) was used to assess the most likely root of embryophytes. We constructed a dataset comprising 24 genomes with the highest BUSCO completion for each lineage sampled (**Supplementary Fig. 4, 5; Supplementary Table 2)**. An unrooted species tree was constructed using IQ-TREE under the LG+C60+G4+F model, as described in the phylogenetics section. The unrooted species tree was then manually rooted on 12 candidate branches, with each alternatively rooted tree scaled to geological time using the mean node ages from the dating analysis. Gene family clusters were inferred by an ‘all-versus-all’ DIAMOND BLAST (Buchfink, Xie and Huson, 2014) with an E-value threshold of 10^−5^, in combination with Markov clustering (MCL) with an inflation parameter of 2.0 (Enright, Van Dongen and Ouzounis, 2002). All gene family clusters were aligned (MAFFT) and trimmed (BMGE), and bootstrap tree distributions were inferred using IQ-TREE as described above. Gene family clusters were reconciled under the 12 candidate root positions tree using the ALEml algorithm (Szöllosi *et al*., 2013). The likelihood of each gene family under each root was calculated; the credible roots were determined using an approximately unbiased (AU) test (Shimodaira and Hasegawa, 2001; Shimodaira, 2002). A detailed description of the ALE implementation can be found at https://github.com/ak-andromeda/ALE_methods/.

### Ancestral gene content reconstruction

Gene family clusters for the genomic dataset were inferred using the same methods as previously described, but the dataset was expanded to contain the genomes of five algal outgroups to allow inference of gene content evolution prior to the embryophyte root **(Supplementary Fig. 6)**. Ancestral gene content was determined by reconciling the gene family clusters with the rooted species tree under the ALEml model. A custom python script called *Ancestral_reconstruction_copy_number.py* was used to identify the presence and absence of gene families on each branch of the tree from the ALE output **(Supplementary Methods).** To functionally annotate the gene families, the consensus sequence of each gene family alignment was inferred using hidden Markov modelling (Eddy, 2011). Consensus sequences were functionally annotated using eggNOG-mapper (Huerta-Cepas *et al*., 2017), and Gene Ontology (GO) terms were summarised using the custom python script – *make_go_term_dictionary.py.* For deeper nodes of the tree where GO terms were infrequent, genes were annotated with the KEGG database using BlastKOALA (Kanehisa, Sato and Morishima, 2016). KEGG annotations were summarised using the python script *kegg_analysis.py.* Additionally, the number of DTL events per branch were calculated using the custom python script *branchwise_number_of_events.py*.

## Results

### Phylogenomics resolves the relationships among land plants

Recent genome and transcriptome sequencing of under-sampled plant lineages - such as hornworts (Li *et al*., 2020), liverworts (Leebens-Mack *et al*., 2019), and early diverging polypod ferns (Huang *et al*., 2020) - in conjunction with improvements to phylogenetic models can address poorly resolved nodes in the embryophyte phylogeny. We compiled a comprehensive dataset of 177 plant and algae species, with improved sampling of previously underrepresented lineages - most notably the addition of two high-coverage hornwort genomes, a lineage difficult to place in previous analyses (Puttick *et al*., 2018; de Sousa *et al*., 2019; Bell *et al*., 2020). In agreement with recent work (Puttick *et al*., 2018; Leebens-Mack *et al*., 2019; Harris *et al*., 2020; Su *et al*., 2021), our analyses recover the monophyly of bryophytes with high support **(Fig. 1**). The best fitting model CAT+GTR+G4 (Blanquart and Lartillot, 2008), accounting for site-rate heterogeneity, supports the monophyly of bryophytes and the sister relationship between mosses and liverworts (Setaphyta) with maximum support (**Fig. 1a**). Consistent with the CAT+GTR+G4 analysis, both the maximum likelihood (ML) and gene tree coalescent analysis (ASTRAL) recovered the monophyly of bryophytes, with ML generating maximum bootstrap support and ASTRAL with 95% support (**Fig. 1b, Fig. 1c**). All other nodes defining major lineages received maximum support across all three analyses.

**Fig. 1.**
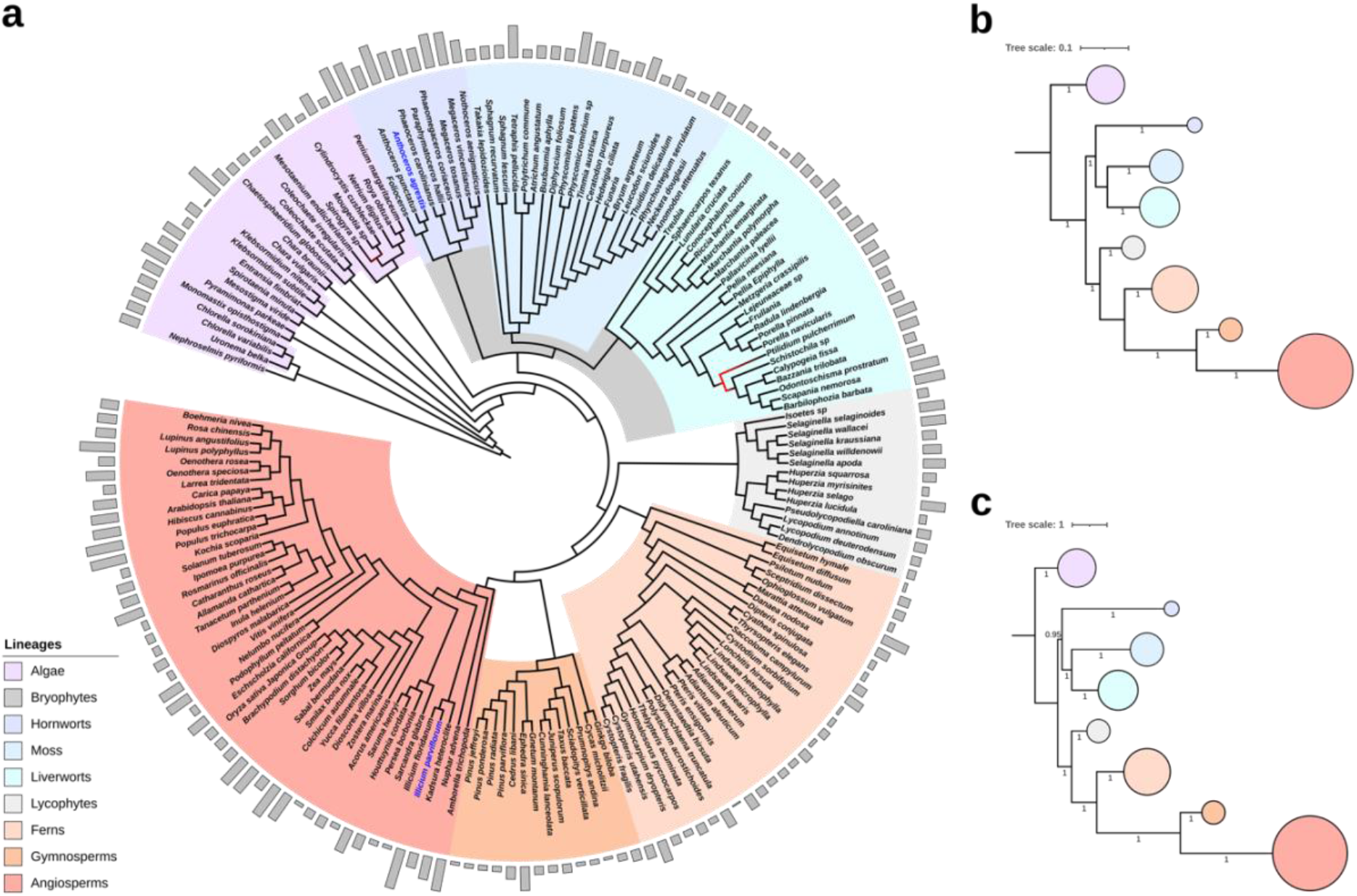
Phylogenetic analysis of land plant provides robust support for the monophyly of bryophytes. a, phylogenetic tree inferred from a concatenated alignment of 30919 sites consisting of 160 single copy orthogroups using the CAT-GTR model (Blanquart and Lartillot, 2008). Branch colour is proportional to the posterior probability; black branches received maximum support, and red received less than maximum and greater values than 0.9. The grey bars assigned to each species are proportional to the percentage of gaps in the alignment. Species with more than 50% gaps in the alignment have their labels coloured blue. The branches of the tree are not drawn to scale. b, Summarised maximum likelihood tree inferred from the same alignment as above using the LG+C60+G4+F model, which accounts for site heterogeneity in the substitution process. All major nodes received maximum boot strap support. c, Phylogenetic tree inferred using the ASTRAL; gene trees were inferred from the 160 single copy orthogroups used to construct the concatenate. All branches except the one defining bryophytes received maximum coalescent support, albeit the branch still received strong support (0.95). The size of the circles in both a and b are proportional to sample size of the lineage they represent.

### Outgroup-free inference of the land plant phylogeny

Rooting phylogenies on an outgroup can influence the ingroup topology due to an artefact known as long branch attraction (LBA) (Bergsten, 2005) and the large evolutionary distance between land plants and their algal relatives has been suggested to be a potential source of bias when inferring the land plant phylogeny (Bell *et al*., 2020). To address the impact of LBA, we used ALE to investigate the support for 12 root positions (**Fig. 2**) on an unrooted embryophyte tree derived from only high-quality genomes (**Supplementary Table 2**). For each topology, we used the divergence time estimates from the molecular clock analysis to convert branch lengths into units of geological time, allowing us to perform time-consistent reconciliations (i.e., to prevent reconciliations in which gene transfers occur into the past). We reconciled 18560 gene families under the 12 rooted and dated embryophyte trees (**Fig. 2a**) and used an approximately-unbiased (AU) test (**Fig. 2b**) to evaluate support for the tested root positions. The AU test rejected all root positions except for a root on the moss stem, hornwort stem, and a root between bryophytes and tracheophytes (*P <* 0.05; **Fig. 2b; Supplementary Table 3**). These three credible roots are in close proximity on the tree and root positions further from this region are rejected with higher confidence (**Fig. 2b; Supplementary Table 3**). To evaluate the nature of the root signal for these three branches, we performed a family-filtering analysis in which families with high DTL rates were sequentially removed and the likelihood re-evaluated. The rationale for this analysis is that the evolution of these families may be poorly described by the model, and so they may contribute misleading signal (Coleman *et al*., 2021). In this case, the root order did not change following removal of high-DTL rate families (**Supplementary Fig. 7**), suggesting broad support for these root positions from the data and analysis. For comparison, we also used an alternative outgroup-free rooting method, STRIDE (Emms and Kelly, 2017), which placed the root between tracheophytes and monophyletic bryophytes (**Supplementary Fig. 8**). Finally, we constrained the topology of the tree inferred from the concatenated alignment (**Fig. 1a**) to be in accordance with the three credible roots (**Fig. 2c**) and computed the likelihood of sequence data along those trees. Trees with embryophyte roots constrained to hornworts and moss were significantly rejected (*P* < 0.05, AU test; **Supplementary Table 4**). Taking our analyses together with other recent work (Puttick *et al*., 2018; Leebens-Mack *et al*., 2019; Harris *et al*., 2020; Su *et al*., 2021) suggests that a root between monophyletic tracheophytes and bryophytes is the best-supported hypothesis of land plant phylogeny. Bryophyte monophyly is therefore the default hypothesis with which to interpret land plant evolution.

**Fig. 2.**
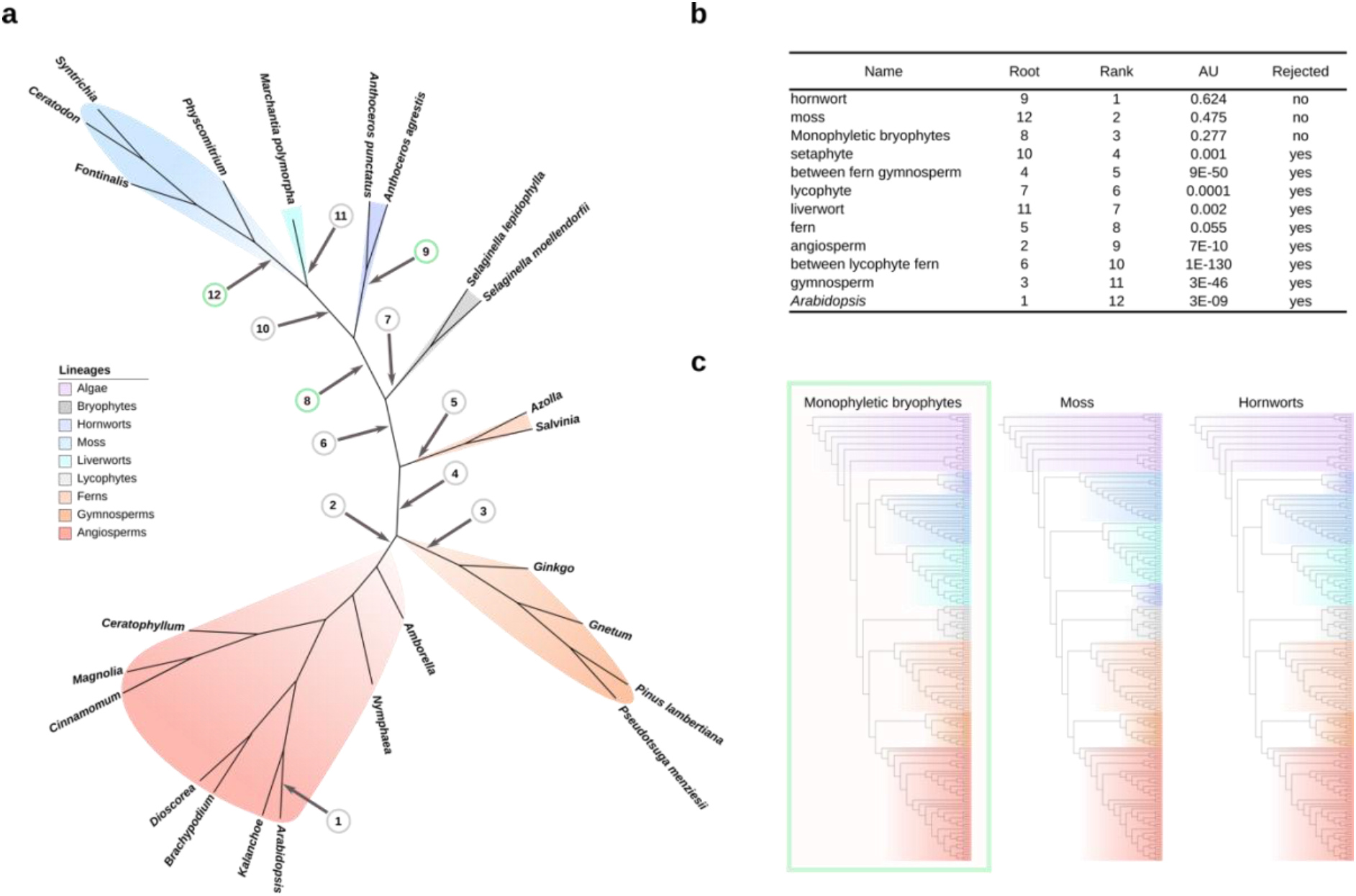
Investigating the root of embryophytes using outgroup-free rooting. **a,** Unrooted maximum likelihood tree inferred from an alignment of 24 species and 249 single copy orthogroups under the LG+C60+G4+F model. Twelve candidate root positions for embryophytes were investigated – including the *Arabidopsis* branch to test the efficacy of the ALE algorithm. The unrooted tree was rooted in each of the twelve candidate positions and scaled to geological time based on the results of the divergence time analysis. 18560 gene clusters were reconciled with each of the twelve rooted and dated species tree using the ALEml algorithm. **b,** The likelihood of the twelve embryophyte roots was assessed with an approximate unbiased (AU) test. The AU test significantly rejected 9 out of the 12 roots, with roots on hornworts, moss, and monophyletic bryophytes (root positions 9, 12 and 8 respectively) comprising the credible set. **c,** Phylogenetic trees constrained to the credible roots were inferred in IQ-TREE (Nguyen *et al.,* 2015) under the LG+C60+G+F model. An AU test was used to assess the likelihood of each of the constrained trees – with the root resulting in monophyletic bryophytes being the only one not to be significantly rejected.

### The evolutionary timeline of land plants

We estimated divergence times on the resolved land plant phylogeny (Fig. 3). We assembled a set of 68 fossil calibrations, representing every major lineage of land plant and notably sampling more bryophyte fossils than previous studies (**Supplementary Methods**). Our results are congruent with those of previous studies (Morris *et al*., 2018) but offer greater precision on many nodes and in some cases greater accuracy (**Supplementary Fig. 9**). This has been leveraged by denser sampling of fossil calibrations, improved taxonomic sampling, especially among bryophytes, and the ability to condition divergence times on a single topology.

**Fig. 3.**
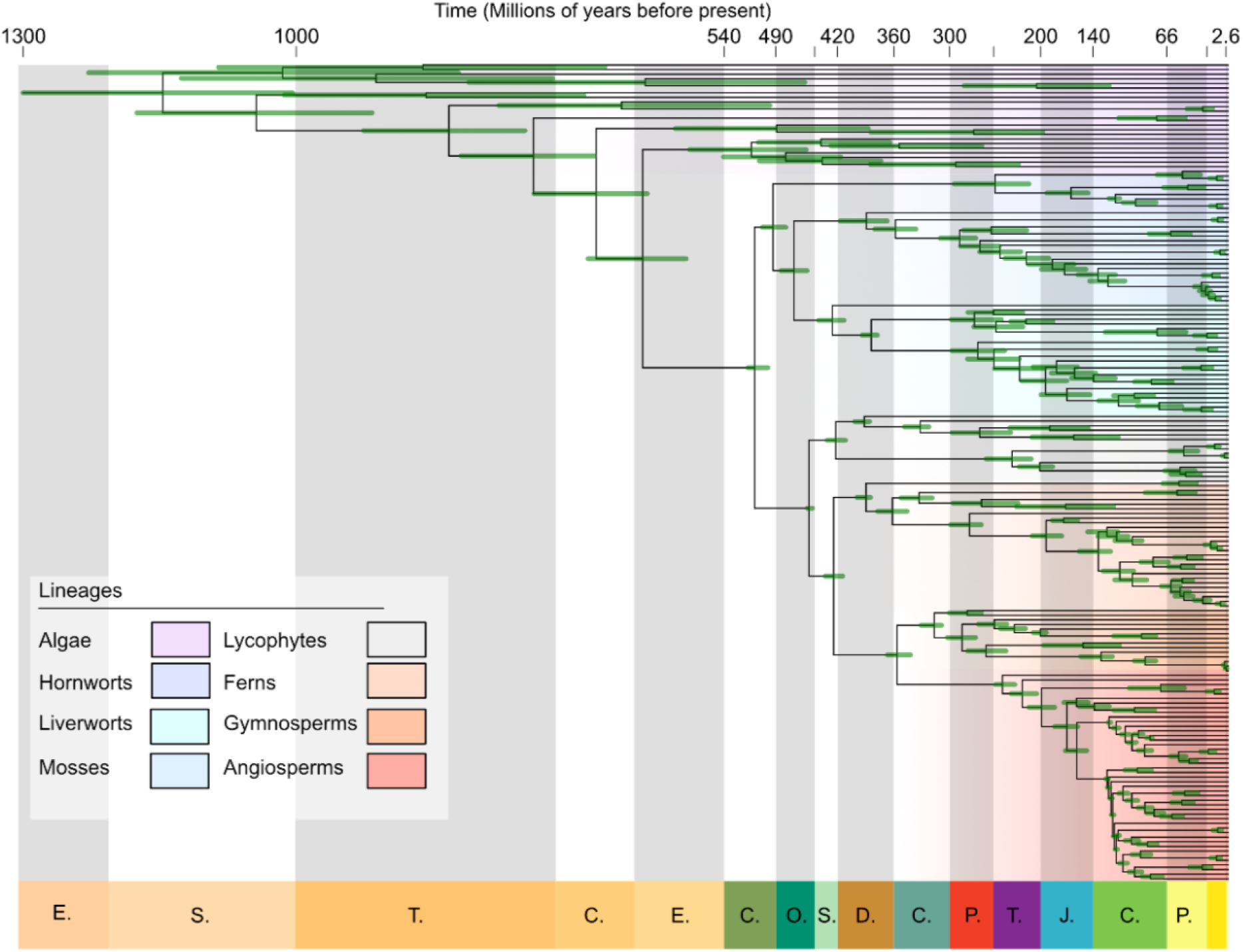
The timescale of land plant evolution. Divergence times in millions of years as inferred using a molecular clock model and 68 fossil calibrations. The inference that the common ancestor of embryophytes lived during the Cambrian is robust to the choice of maximum ages (see Supplementary Methods). The divergence times of hornworts are constrained by a horizontal gene transfer from hornworts into ferns, with the result that the hornwort crown is inferred to be older than estimated previously. The posterior age distribution from the relative node dating analysis was used as a prior for the present analysis (see Methods). Nodes are positioned on the mean age, with bars represent the 95% highest posterior density (HPD).

The role and influence of fossil calibrations in molecular clock studies, especially maximum age calibrations, remains controversial (Hedges *et al*., 2018; Su *et al*., 2021). While the fossil record is an incomplete representation of past diversity, our analyses account for this uncertainty in the form of soft minima and maxima. Morris et al. (2018) inferred a relatively young age for the embryophyte crown-ancestor (515-470 Ma), making use of a maximum age constraint based on the absence of embryophyte spores in strata for which fossilisation conditions were such that spores of non-embryophyte algae have been preserved. Hedges et al. (2018) and Su et al. (2021) argued against the suitability of this maximum age constraint on the basis that calibrations derived from fossil absences are unreliable and that the middle Cambrian maximum age exerts too great an influence on the posterior estimate. To assess the sensitivity of our approach to the effect of maximum age calibrations, we repeated clock analyses with less informative maximum age calibrations (**Supplementary Methods)**. Removing the maximum age constraint on the embryophyte node produced highly similar estimates to when the maximum in employed (**Supplementary Fig. 10**). Relaxing all maxima did result in more ancient estimates for the origin of embryophytes, although still considerably younger than recent studies (Su *et al*., 2021) extending the possible origin for land plants back to the Ediacaran (540-597 Ma; **Supplementary Fig. 10**). Older ages estimated in Su et al. (2021) appear to reflect, in part, differences in the phylogenetic assignment of certain fossils (**Supplementary Methods**), such as the putative algae *Proterocladus antiquus* and the liverwort *Ricardiothallus devonicus,* rather than a dependence on the maximum age calibration. Our results reject the possibility that land plants originated during the Neoproterozoic, instead supporting an origin of the land plant crown group during the Cambrian, 515-493 Ma, with crown tracheophytes and crown bryophytes originating 452-447 Ma and 500-473 Ma, respectively. Within bryophytes, Setaphyta (mosses + liverworts) had diverged by 479-450 Ma, with the divergence of crown mosses by 420-364 Ma and liverworts 440-412 Ma. Among tracheophytes, the origin of lycophytes is estimated during the late Silurian to Early Devonian, 431–411 Ma, coeval with the origin of euphyllophytes 432–414 Ma.

### Constraining the timing of hornwort evolution with horizontal gene transfer

Previous estimates for the age of hornworts have been limited by taxonomic sampling (Morris et al. 2018), the limited number of fossil calibrations (Villarreal and Renner, 2012), or have made use of a strict clock calibrated based on an external rate (Villarreal and Renner, 2014). In the absence of fossil calibrations on deep nodes, hornworts are characterised by an ancient stem lineage and the youngest crown lineage among land plants. While this result might be correct, it could also result from the scarcity of the fossil record for hornworts (Villarreal and Renner, 2014). To investigate, we implemented a relative node age constraint based on the horizontal transfer of the chimeric photoreceptor NEOCHROME from hornworts into ferns (Li *et al*., 2014). In the absence of direct fossil calibrations for hornworts, this gene transfer provides a relative constraint that ties the history of hornworts to that of ferns, for which more fossils are available. The effect of the relative age constraint is to make the crown group older (294-214 Ma; Fig. 3) and thus shorten the length of the stem group, with divergence times within the crown group all moving older. This is older than the earliest unequivocal fossils assigned to hornworts. However, given the scarcity of hornwort fossils, it seems likely that the fossil record underestimates the true age of this clade.

### Gene content of the embryophyte common ancestor

We used gene tree-species tree reconciliation to estimate the gene content of the embryophyte common ancestor **(Supplementary Table 6)**. We used the genome dataset from the ALE rooting analysis with the addition of 5 algal genomes, to better place the origin of families that pre-date the origin of embryophytes (**Supplementary Fig. 11**). The tree was dated following the same methodology as the larger dating analysis while using an applicable subset of calibrations, allowing the use of a dated reconciliation algorithm (ALEml) to improve estimation of duplication, transfer, and loss events (**Supplementary Fig. 12**).

Analysis of ancestral gene content highlighted considerable gene gain along the ancestral embryophyte branch (Fig. 4a; **Supplementary Table 7)** – a substantial number of duplications defined this transition, with fewer transfers and losses observed. Our analysis suggests that the common ancestor of embryophytes and Zygnematales had more of the building blocks of plant complexity than extant Zygnematales, which appear to have undergone a loss of 1442 gene families since their divergence, the largest loss observed on the tree (Fig. 4a). Functional characterisation of the genes lost in the Zygnematales using the KEGG database identified gene families involved in the production of cytoskeletons, exosomes and phenylpropanoid synthesis **(Supplemental Table 8)**. Exosomes and complex cytoskeletons are essential for multicellular organisms to function (Raposo and Stoorvogel, 2013; Chen and Wang, 2019), and the inferred loss of these gene families is consistent with the hypothesis that the body plan of the algal ancestor of embryophytes was multicellular (de Vries and Archibald, 2018), rather than possessing the single cell or filamentous architecture observed in extant Zygnematales. The more complex cytoskeleton could be associated with increased rigidity, helping overcome the gravitational and evaporative pressures associated with the transition to land (Raven, 2002). Interestingly, phenylpropanoids are associated with protection against UV irradiance (Popper *et al*., 2011) and homiohydry (de Vries and Archibald, 2018), suggesting the common ancestor may have been better adapted to a terrestrial environment than extant Zygnematales.

**Fig. 4.**
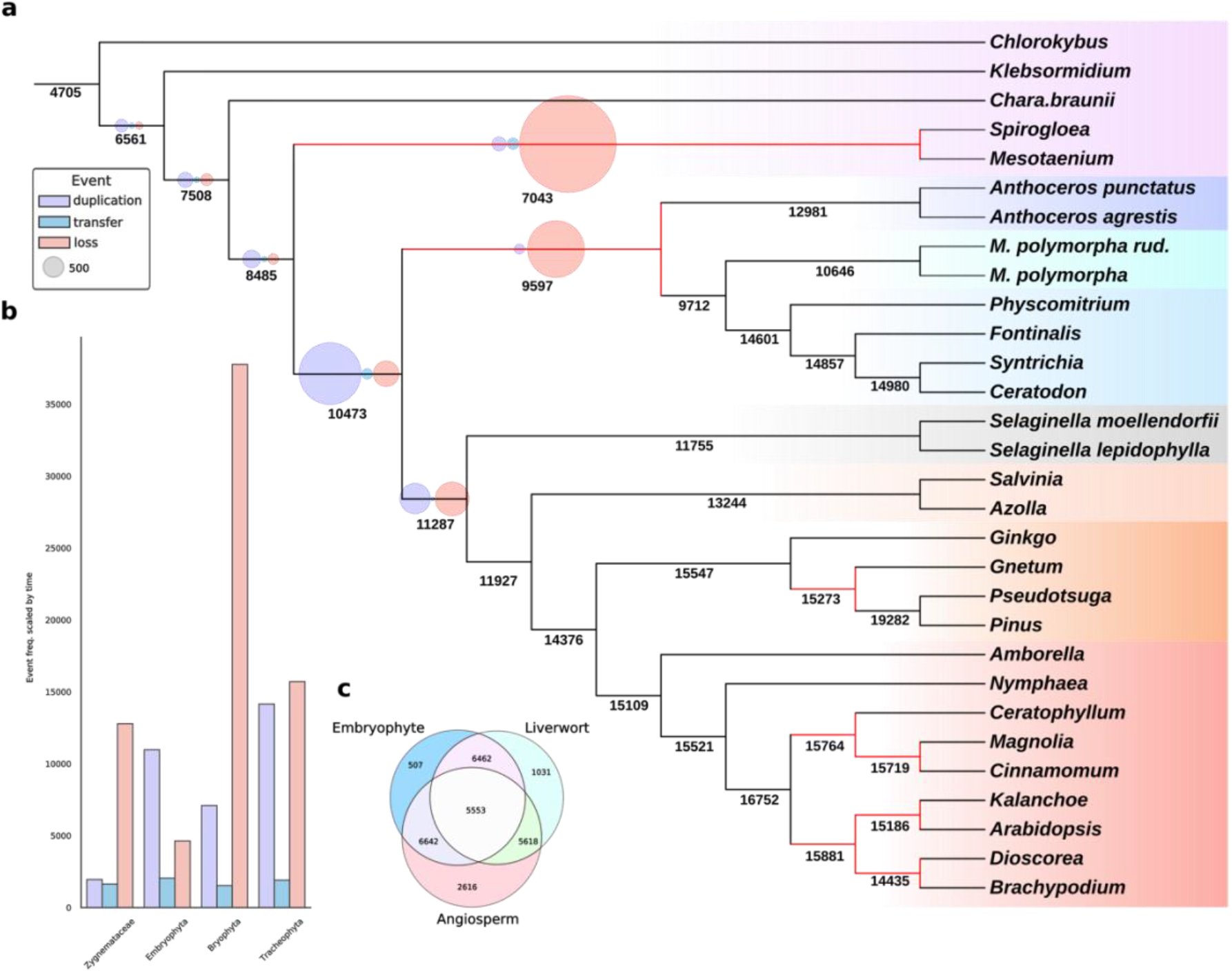
Gene content reconstruction of the ancestral embryophyte. **a,** Ancestral gene content was inferred for the internal branched of the embryophyte tree. A maximum likelihood tree was inferred from an alignment of 30 species of plants and algae, comprising of 185 single copy orthologs and 71855 sites, under the LG+C60+G4+F model (Le and Gascuel, 2008) in IQ-TREE (Nguyen *et al.,* 2015), and rooted in accordance with our previous phylogenetic analysis. A timescale for the tree was then calculated using a subset of 18 applicable fossil calibrations in MCMCtree. 20822 gene family clusters, inferred using MCL (Enright et al., 2002), were reconciled against the rooted dated species tree using the ALEml algorithm (Szöllosi *et al.,* 2013). The summed copy number of each gene family (under each branch) was determined using the custom python code – branchwise_number_of_events.py. Branches with reduced copies from the ancestral node are coloured in red. The number of duplications, transfer, and loss (DTL) events are represented by purple, blue and red circles, respectively. The size of circles is proportional to the summed number of events – scale indicated by grey circle. **b,** The number of DTL events scaled by time for four clade-defining branches in the embryophyte tree. **c,** The number of shared gene families between the ancestral embryophyte, liverwort and angiosperm – the ancestral embryophyte shares more gene families with the ancestral angiosperm than the ancestral liverwort.

We also observed greater gene loss along the bryophyte stem lineage (Fig. 4a). While the total number of genes lost on the bryophyte stem was not the largest loss event observed, the rate of gene loss (in terms of gene families/year) was substantially greater than all other major clades (Fig. 4b). GO term functional annotation of the gene families lost in bryophytes reveal reductions in shoot and root development from the ancestral embryophyte (**Supplementary Table 9; Supplementary Fig. 13)**. To investigate the evolution of genes underlying morphological differences between tracheophytes and bryophytes, we investigated the evolutionary history of gene families containing key *Arabidopsis* genes for vasculature and stomata (**Supplementary Table 11**); gene families associated with both vasculature and stomatal function exhibited lineage-specific loss in bryophytes (**Supplementary Fig. 14**). Specifically, three orthologous gene families that are critical to vasculature development in *Arabidopsis*, which contain WOX4, WER and STM respectively, were inferred to be lost on the bryophyte stem (Fig. 4, **Supplementary Table 11**. To investigate these inferred losses in more detail, we manually curated sequence sets and inferred phylogenetic trees for these families (**Supplementary Fig. 15**). These analyses of individual gene families corroborated the pattern of loss along the branch leading to bryophytes. The loss of these orthologous gene families strengthens the hypothesis that ancestral embryophyte had a more complex vasculature system than that of extant bryophytes (Donoghue *et al*., 2021). Overall, the loss of gene families (Fig. 4) and change in GO term frequency (**Supplementary Fig. 13**) suggests widespread reduction in complexity in bryophytes, and the ancestral embryophyte being more complex than previously envisaged. Indeed, gene loss defines the bryophytes early in their evolutionary history, but large numbers of duplication and transfers events are observed following the divergence of the setaphytes and hornworts (**Supplementary Table 6**), with (for example) extant mosses boasting a similar gene copy number to tracheophytes (Fig. 4).

## Discussion

We have presented a time-scaled phylogeny for embryophytes, which confirms the growing body of evidence that bryophytes form a monophyletic group (Fig. 1, Fig. 2) and our precise estimates of absolute divergence times provide a robust framework to reconstruct genome evolution across early land plant lineages (Fig. 4). Our results confirm that many well-characterised gene families predate the origin of land plants (Bauer *et al*., 2013; Wilhelmsson *et al*., 2017; Bowles, Bechtold and Paps, 2020; Cannell *et al*., 2020; Harris *et al*., 2020). However, our analyses also show that extensive gene loss has also characterised the evolution of major embryophyte groups. Reductive evolution in bryophytes has been demonstrated previously, where the loss of several genes has resulted in the lack of stomata (Harris *et al*., 2020). Our results suggest that these patterns of gene loss are not confined to stomata but are instead pervasive across bryophyte genomes, and that much of the genome reduction occurred during a relatively brief period of c.20 million years following their divergence from tracheophytes during the Cambrian. The evolutionary pressures that underlay this “Cambrian Reduction”, and the ways in which gene loss contributed to the evolution of the bryophyte body plan (such as the loss of genes associated with vasculature), remain unclear. It has been proposed that the radiation of vascular plants, heralded by the increased diversity of trilete spores in the palynological record, relegated bryophytes to a more marginal niche (Wellman, Steemans and Vecoli, 2013). However, it seems possible that bryophytes independently evolved to exploit this niche, shedding the molecular and phenotypic innovations of embryophytes where they were no longer necessary. A large body of research has focussed on the importance of gene and whole-genome duplication in generating evolutionary novelty in land plant evolution (Chanderbali *et al*., 2017; Clark and Donoghue, 2018; Walden *et al*., 2020; Stull *et al*., 2021). However, gene loss is an important driver of phenotypic evolution in other systems (Albalat and Cañestro, 2016; O’Malley, Wideman and Ruiz-Trillo, 2016; Guijarro-Clarke, Holland and Paps, 2020), notably in flying and aquatic mammals (Sharma *et al*., 2018) and yeast (Helsen *et al*., 2020). It has also been shown that rates of genome evolution, rather than absolute genome size, correlate with diversification across plants (Puttick, Clark and Donoghue, 2015). Extant bryophytes remain highly diverse, and it is possible that bryophytes represent another example of specialisation and evolutionary success via gene loss.

Bryophytes have sometimes been used as models in physiological and genetic experiments to infer the nature of the ancestral land plant. Our analysis suggests that modern bryophytes are highly derived: in terms of gene content, our analysis suggests that the ancestral angiosperm may have shared more genes with the ancestral land plant than did the ancestral liverwort (Fig. 4c). Such differences in gene content between species can be visualised as an ordination, where the two-dimensional distances between species represent dissimilarity in gene content. Reconstructed gene content at ancestral nodes can be projected into this space, showing the evolution of gene content along the phylogeny (Fig. 5). These genome disparity analyses reveal that the genomes of bryophytes and tracheophytes are both highly derived. Neither lineage occupies an ancestral position, with lineage-specific gene gain and loss events driving high disparity in both bryophytes and tracheophytes, reinforcing the view that there are no extant embryophytes that uniquely preserve the ancestral state (Puttick *et al*., 2018; Rensing, 2018; Rich and Delaux, 2020). Despite the paucity of data for some groups, these analyses reveal that the diversity among bryophyte genomes is comparable to those of tracheophytes. These results are perhaps unsurprising given that bryophytes have been evolving independently of tracheophytes since the Cambrian and the similarly ancient divergence of each of the major bryophyte lineages, but emphasise the point that, in general terms, bryophytes serve as no better a proxy for the ancestral land plant than do tracheophytes. Our results therefore agree that a view of bryophytes as primitive plants may mislead inferences of ancestral gene content or character evolution (McDaniel, 2021). Instead, the best model organism(s) for investigating the nature of early plants will depend on the trait being investigated, alongside a careful appraisal of the phylogenetic diversity, including algal outgroups. Likewise, interpretations of the early land plant fossil record have been contingent on the first land plants appearing more like extant bryophytes than tracheophytes. That the ancestral embryophyte may have been more complex than living bryophytes is in keeping with many early macrofossils being both more complex than bryophytes and possessing a mosaic of tracheophyte and bryophyte traits (Edwards *et al*., 2014; Donoghue *et al*., 2021).

**Fig. 5.**
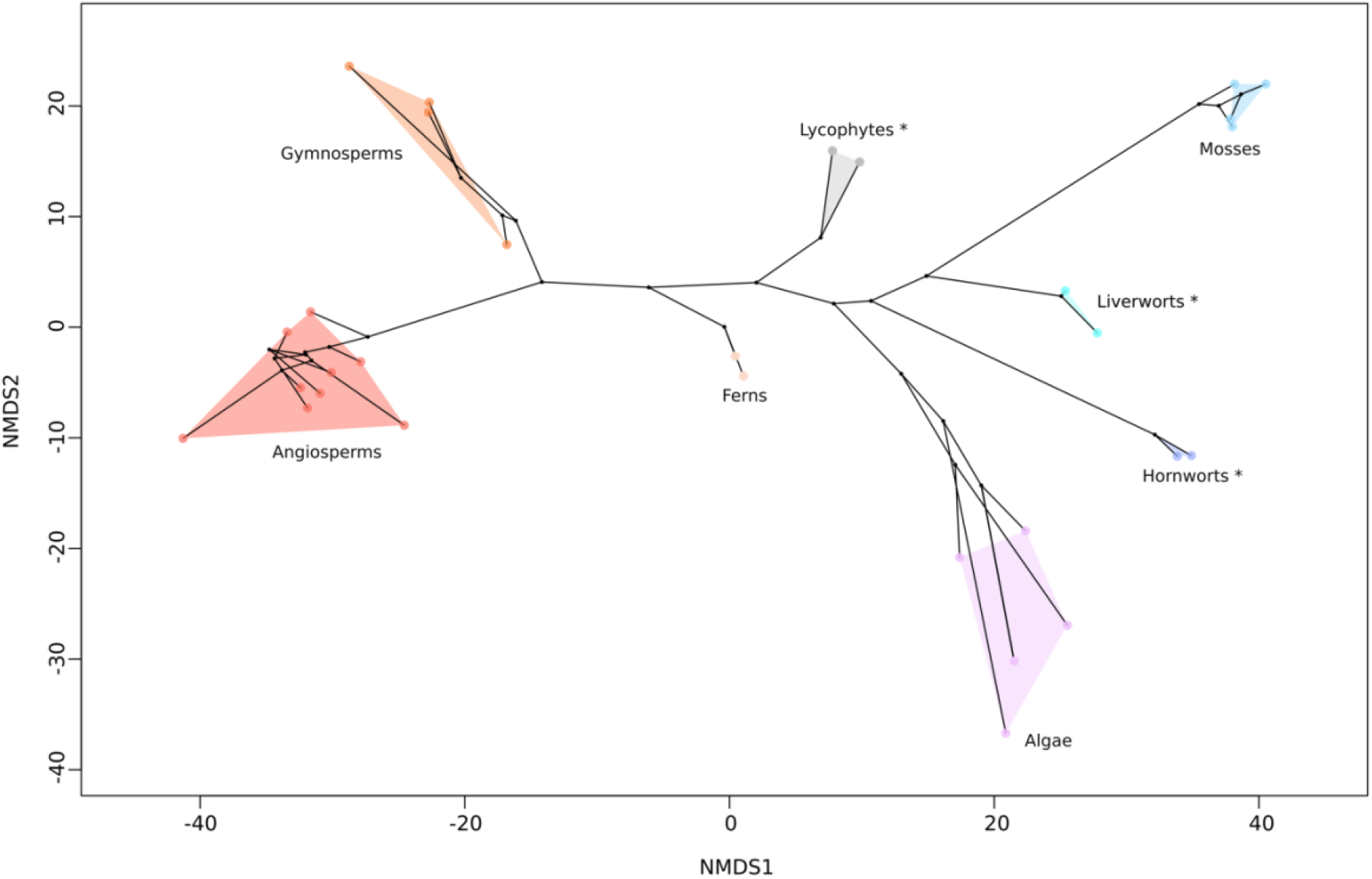
Genome disparity analysis demonstrates that the gene content of both tracheophytes and bryophytes is highly derived. Non-metric multidimensional scaling (NMDS) analysis of the presence/absence of gene families. The presence or absence of each gene family was determined from the ALE analysis for each tip and internal node in the phylogeny. Presence/absence data was used to calculate the Euclidean distance between species and nodes, which was then ordinated using NMDS. Branches were drawn between the nodes of the tree, with convex hulls fit around members of each major lineage of land plants.

## Author Contributions

All authors conceived the study and designed experiments; All experiments were performed by BJH, JWC and DS; All authors contributed to the interpretation of results; All authors contributed to the drafting of the manuscript.

## Acknowledgements

TAW, JC and AMH are supported by a Leverhulme Trust Research Project Grant (RPG-2019-004). TAW is also supported by a Royal Society University Research Feollowship (URF\R\201024). BJH is supported by a PhD studentship from the New Phytologist Trust. PCJD was funded by Natural Environment Research Council (NERC) grant (NEP013678/1), part of the Biopshere, Evolution, Transitions and Resilience (BETR) programme, which is co-funded by the Natural Science Foundation for China (NSFC); as well as Biotechnology and Biological Sciences Research Council (BBSRC) grant (BB/T012773/1). GJSz and DS are supported by the European Research Council under the European Union’s Horizon 2020 research and innovation programme under grant agreement no. 714774.

## Data availability

All data are available on FigShare at https://doi.org/10.6084/m9.figshare.c.5682706. Scripts and code available at: https://github.com/ak-andromeda/ALE_methods/ and https://github.com/dschrempf/mcmc-date.

---

**Supplementary Figure 1.**
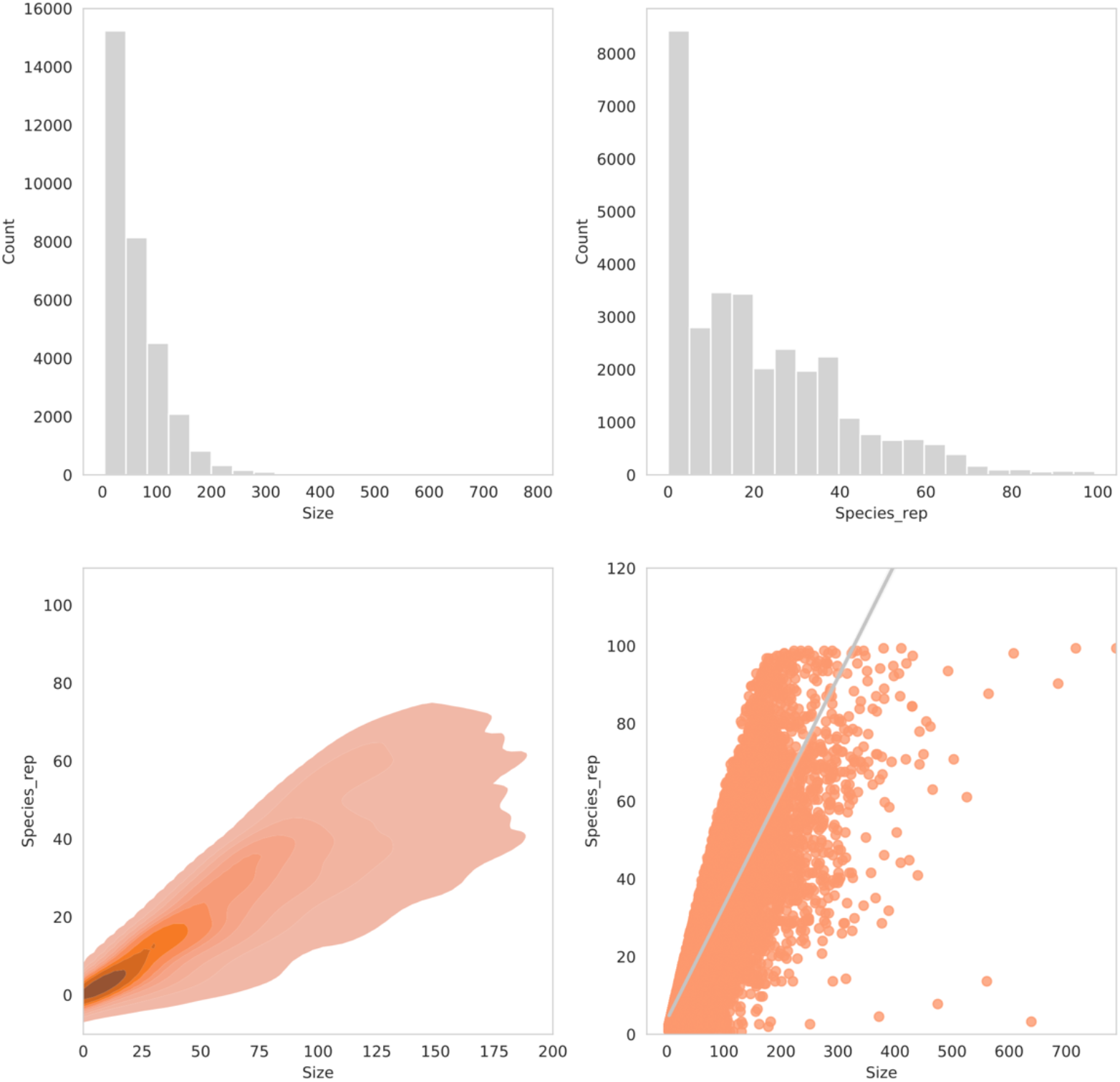
Gene family clusters for genome and transcriptome dataset – 154 species. Top left, distribution of gene family size. Top right, distribution of gene family species representation. Bottom left, density of gene family species repesentation and size. Bottom right, regression of gene family species representation and size.

**Supplementary Figure 2.**
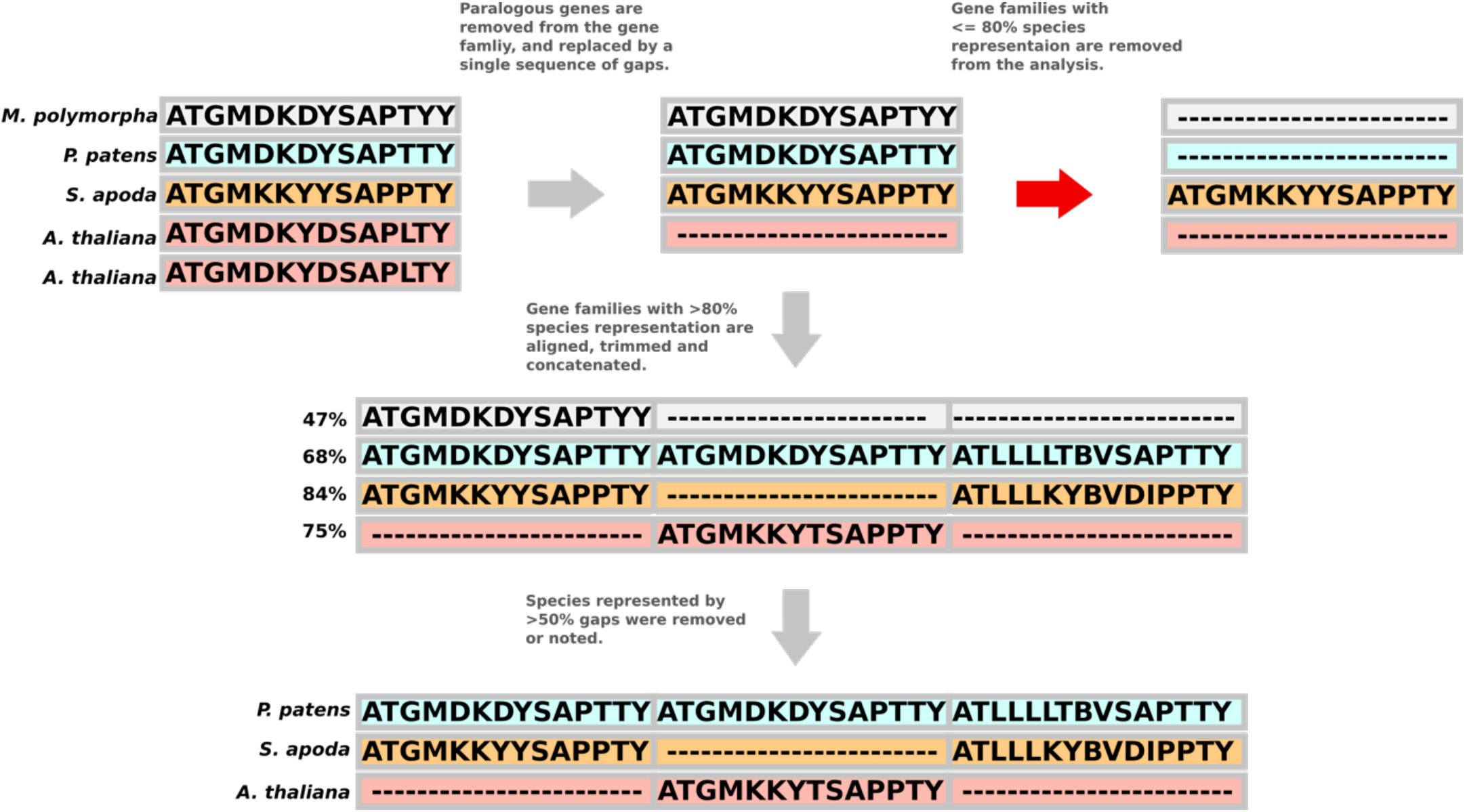
**Graphical representation of orthology inference algorithm**. Species represented by more than one gene copy are removed from the gene family alignment and replaced by a single sequence of gaps. Gene families with more than 80% of the original species present are retained. Gene families are then aligned, trimmed and concatenated together to form a super matrix. Species with less than 50% gaps in the alignment are removed or noted if within 10% error – this step prevents single species that are continually removed from gene families to be in the final alignment.

**Supplementary Figure 3.**
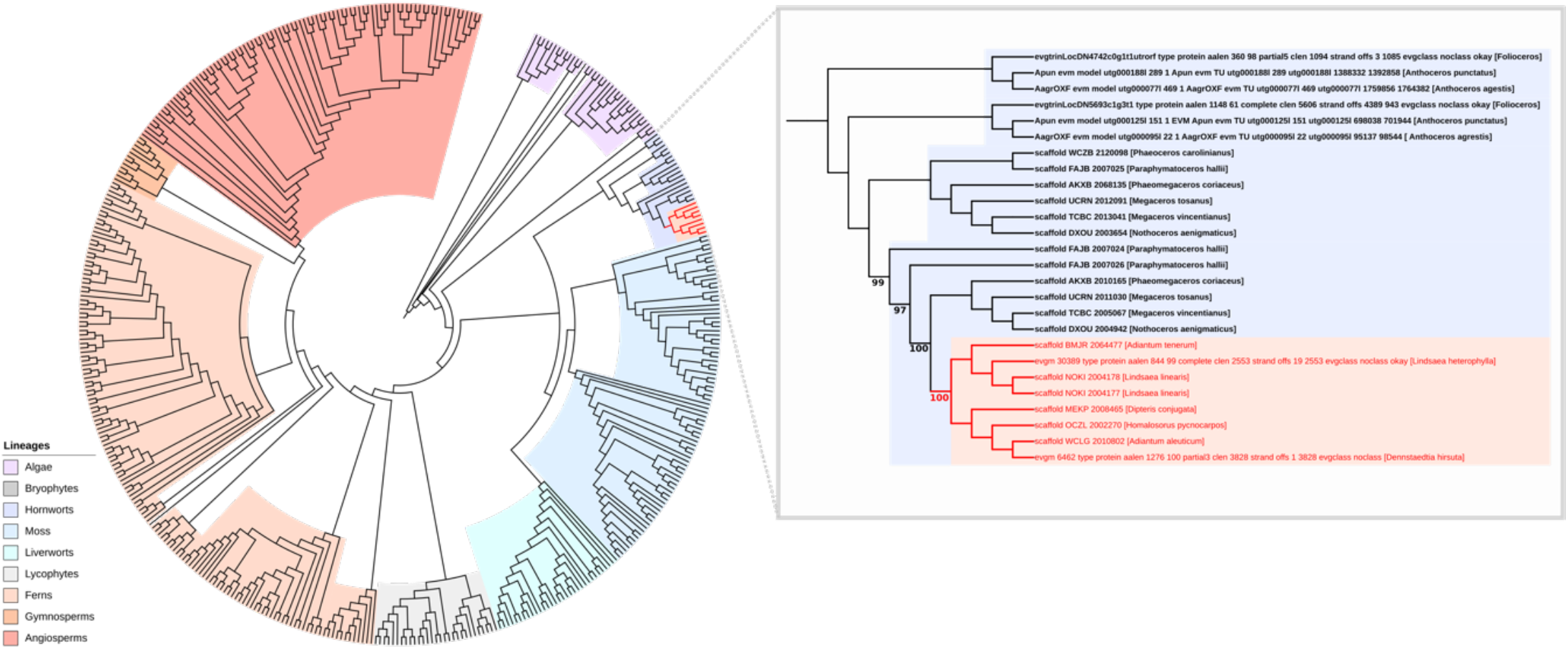
Phylogenetic tree highlighting the horizontal transfer of the chimeric neochrome photoreceptor (NEO). The *Arabidopsis thaliana* protein sequence for PHOT1 was used to BLAST a database of 177 species of plant and transcriptomes. The homologous sequences were aligned with MAFFT and trimmed with BMGE. A maximum likelihood tree was inferred in IQ-TREE under the best fitting substitution model inferred with Bayesian Inference Criterion. 8 fern genes were resolved within the hornworts and were inferred to have undergone horizontal gene transfer (coloured red). This transfer was previously characterised (Li *et al.,* 2014), and we corroborate this finding with maximum bootstrap support.

**Supplementary Figure 4.**
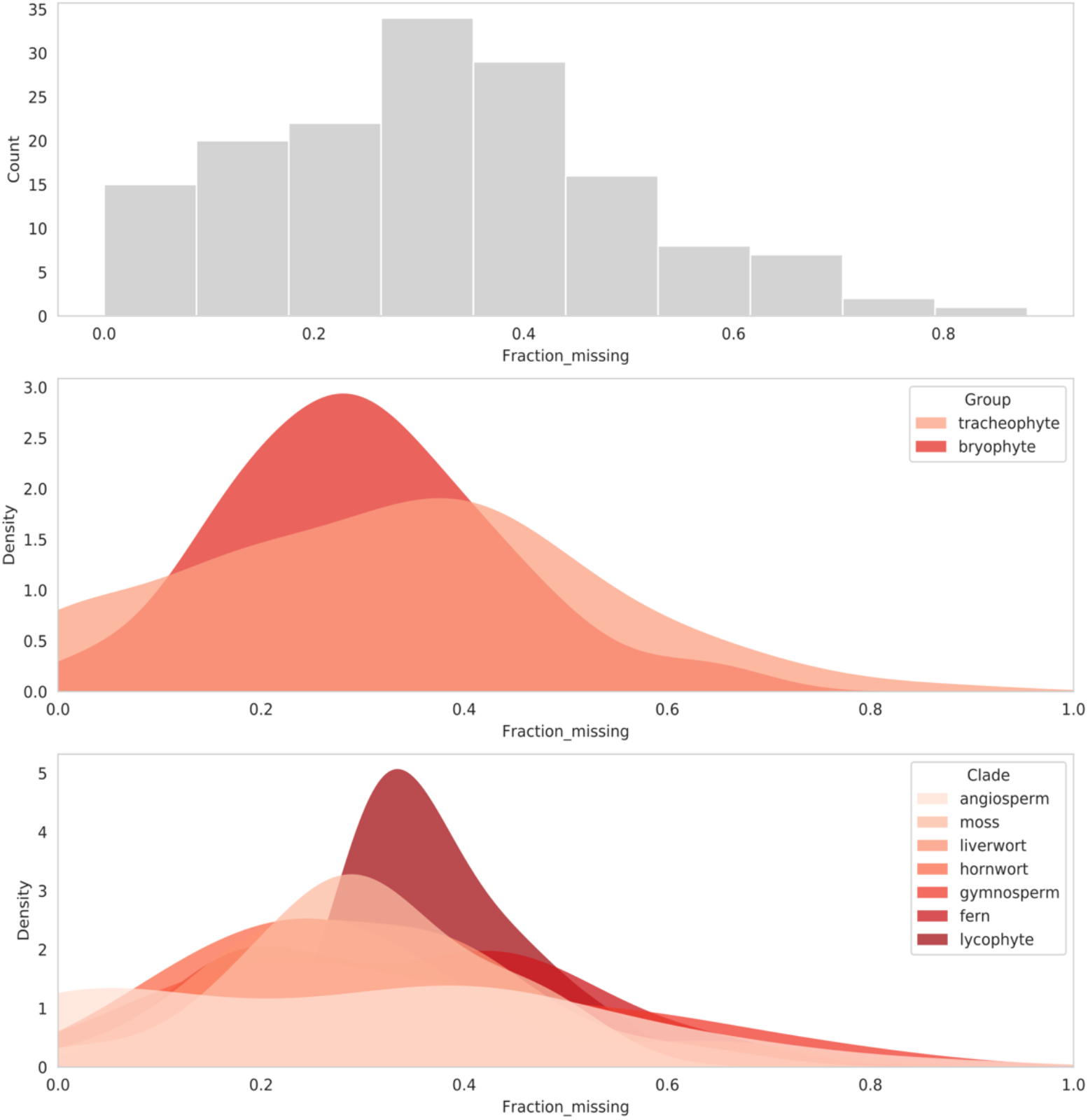
BUSCO completion of genome and transcriptome dataset – 154 species. Top, histogram of BUSCO completeness. Middle, distribution of BUSCO completeness separated into the two main plant groups. Bryophytes, on average having more complete BUSCOs than tracheophytes. Bottom, distribution of BUSCO completeness separated by lineage. Angiosperms had the most varied completeness values. As the dataset comprised of both genomes and transcriptomes, the underlying variation per lineage may be ascribed to the more incomplete transcriptomes.

**Supplementary Figure 5.**
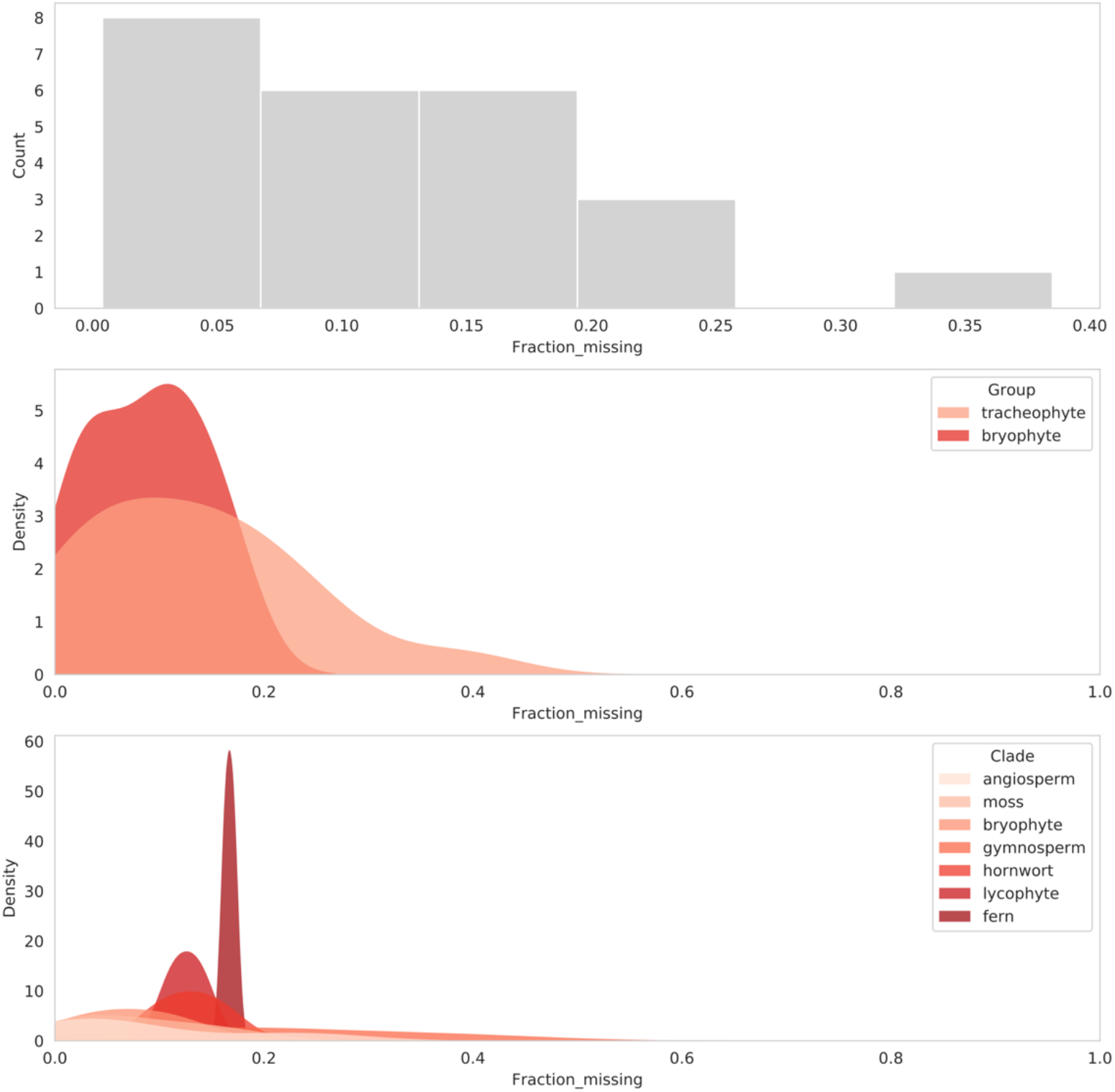
BUSCO completion of genome only dataset – 24 species. Top, histogram of BUSCO completeness. Middle, distribution of BUSCO completeness separated into the two main plant groups. Bryophytes, on average having more complete BUSCOs than tracheophytes. Bottom, distribution of genome completeness separated by lineage. BUSCO completeness is significantly higher for the genome only dataset.

**Supplementary Figure 6.**
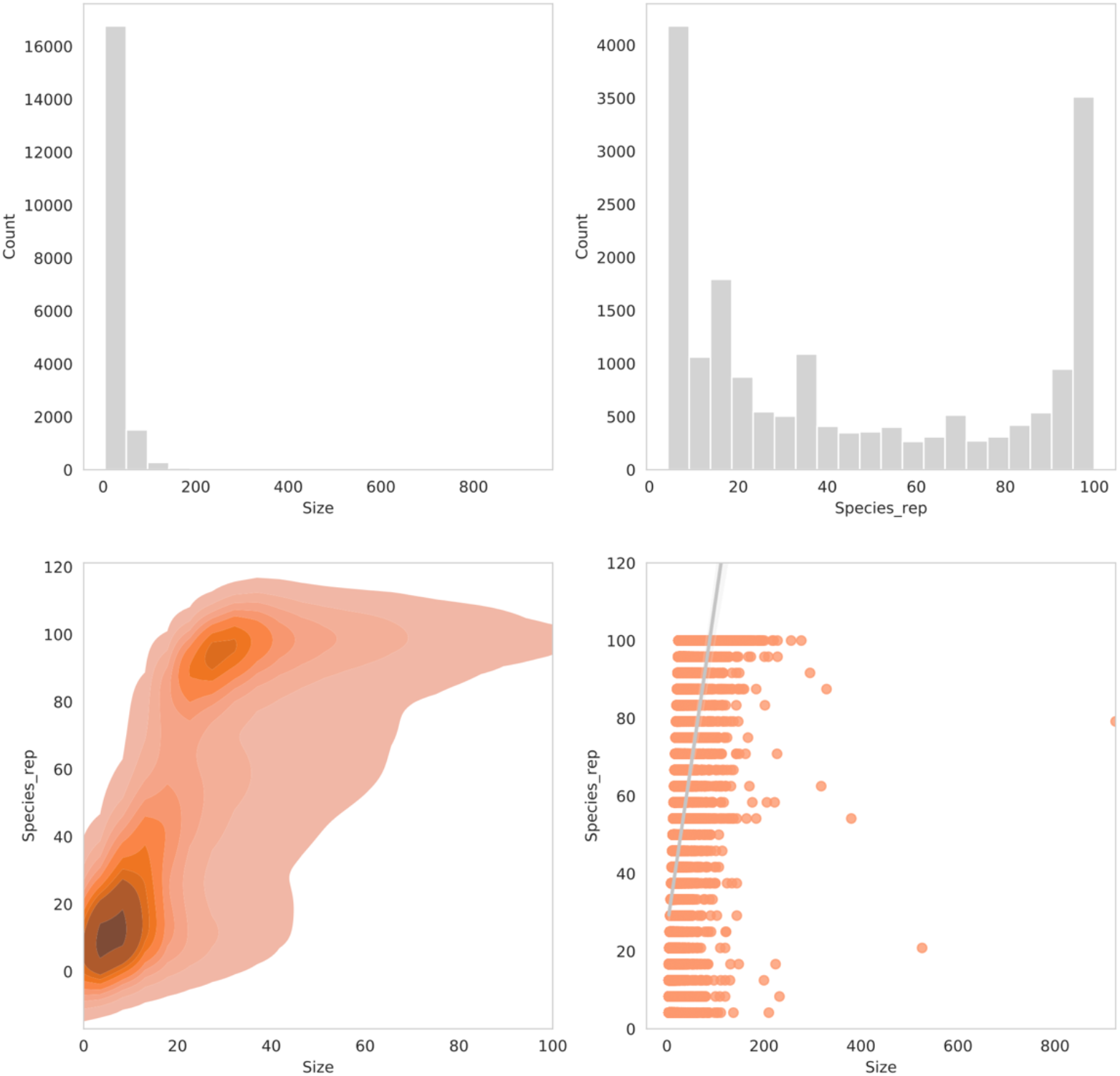
Gene family clusters for genome dataset - 24 species. Top left, distribution of gene family size. Top right, distribution of gene family species representation. Bottom left, density of gene family species repesentation and size. Bottom right, regression of gene family species representation and size.

**Supplementary Figure 7.**
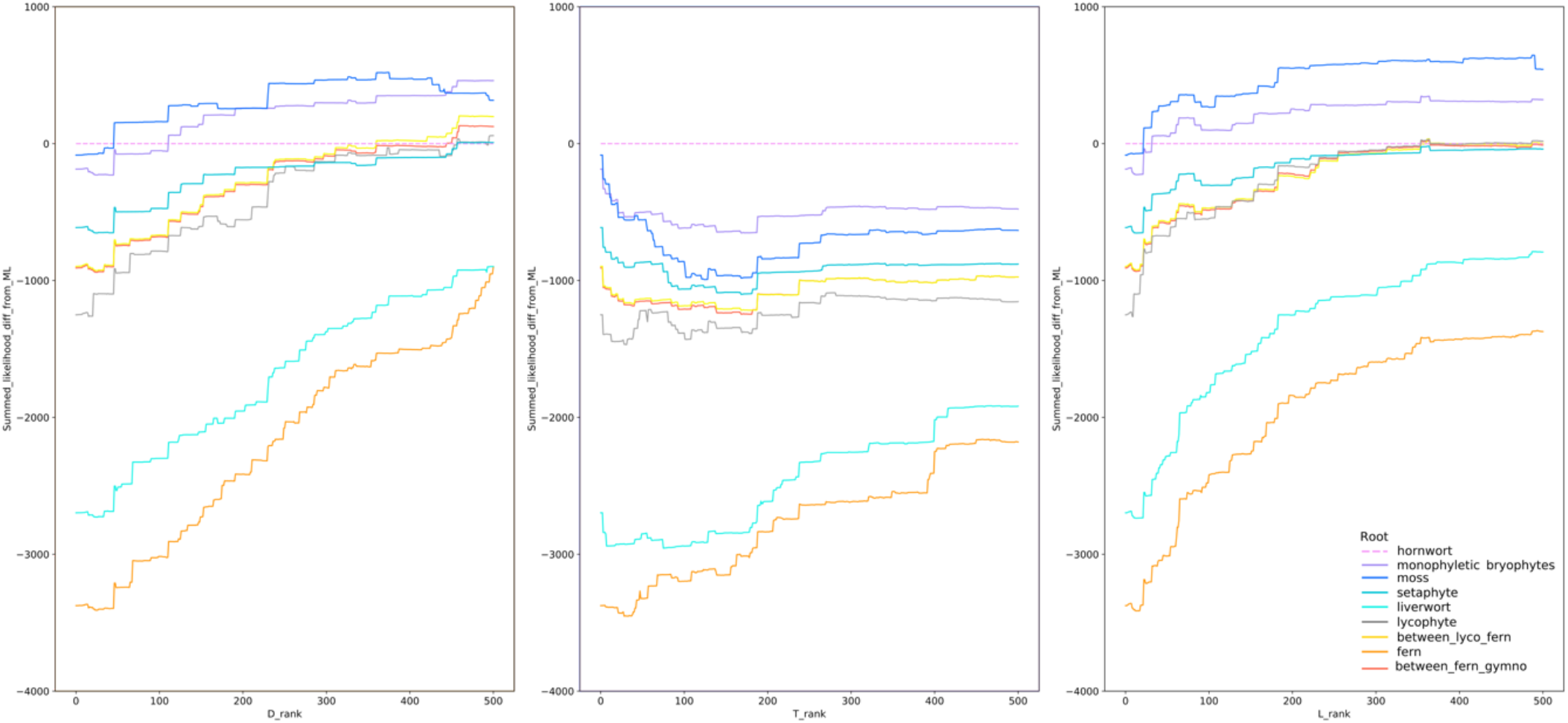
Evaluating the root signal inferred by ALE. The impact of high duplication, transfer or loss rate families on inferred root order was investigated by iteratively removing top-ranked families and recalculating the summed likelihood on remaining families. While the roots of the credible set remained the top three by summed likelihood, their order varied as high D, T or L-rate families were removed. Removal of high-D and high-L families reduced support for the hornwort root (the ML root on all data), while removal of high-T families increased support for the hornwort root relative to the second- and third-ranked roots. As D, T and L rates were calculated based on the ML root, the preference of high-T families for alternative roots might reflect the difficulty of placing the sparsely-sampled hornworts accurately in individual gene trees.

**Supplementary Figure 8.**
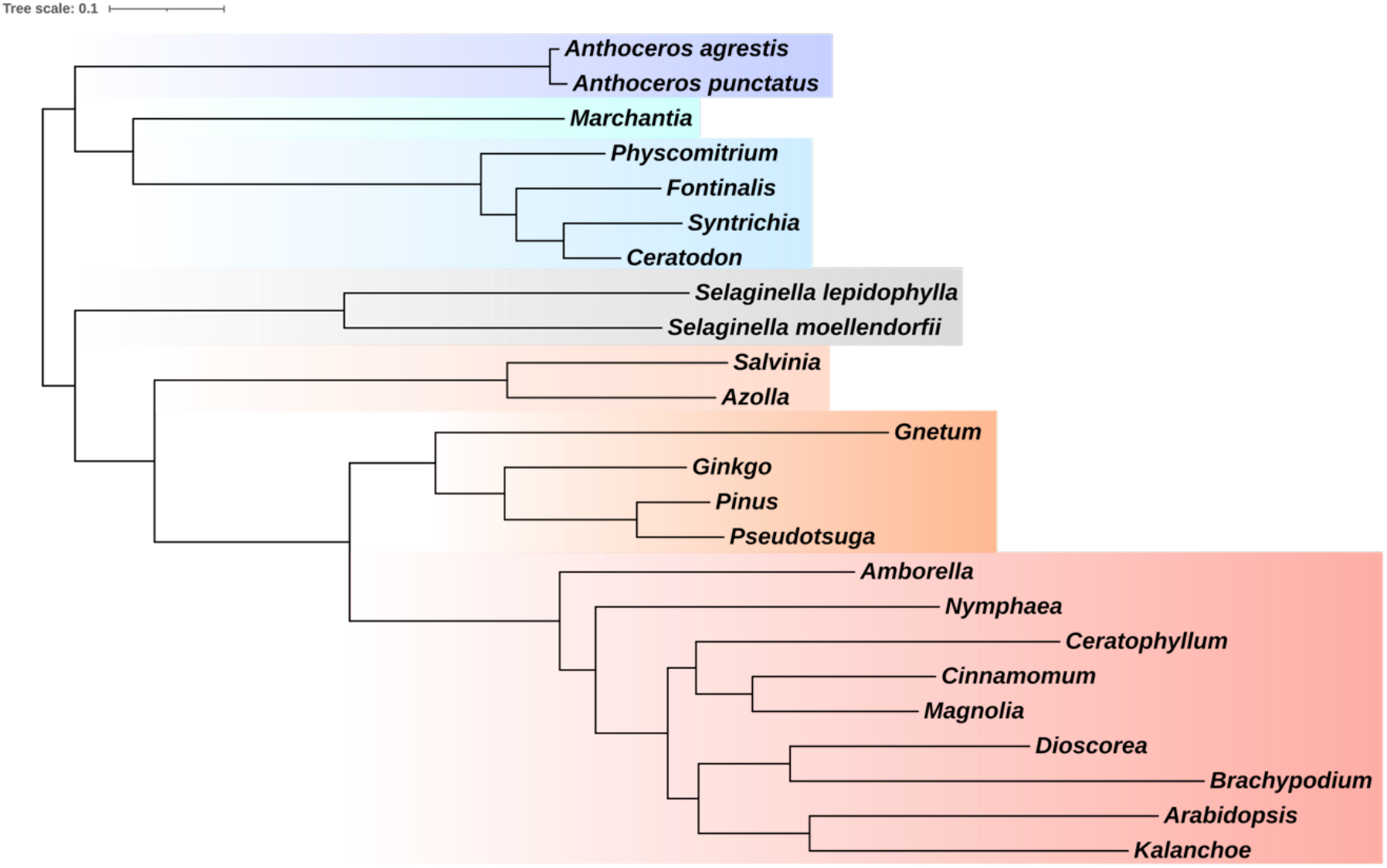
Embryophyte tree rooted with STRIDE. OrthoFinder 2.0 (Emms & Kelly, 2020) was used to infer orthology groups from the 24 high quality genomes. 23145 orthogroups were inferred. Gene duplication events in each of the orthogroups were assessed using STRIDE (Emms & Kelly, 2017) to root the species tree.

**Supplementary Figure 9.**
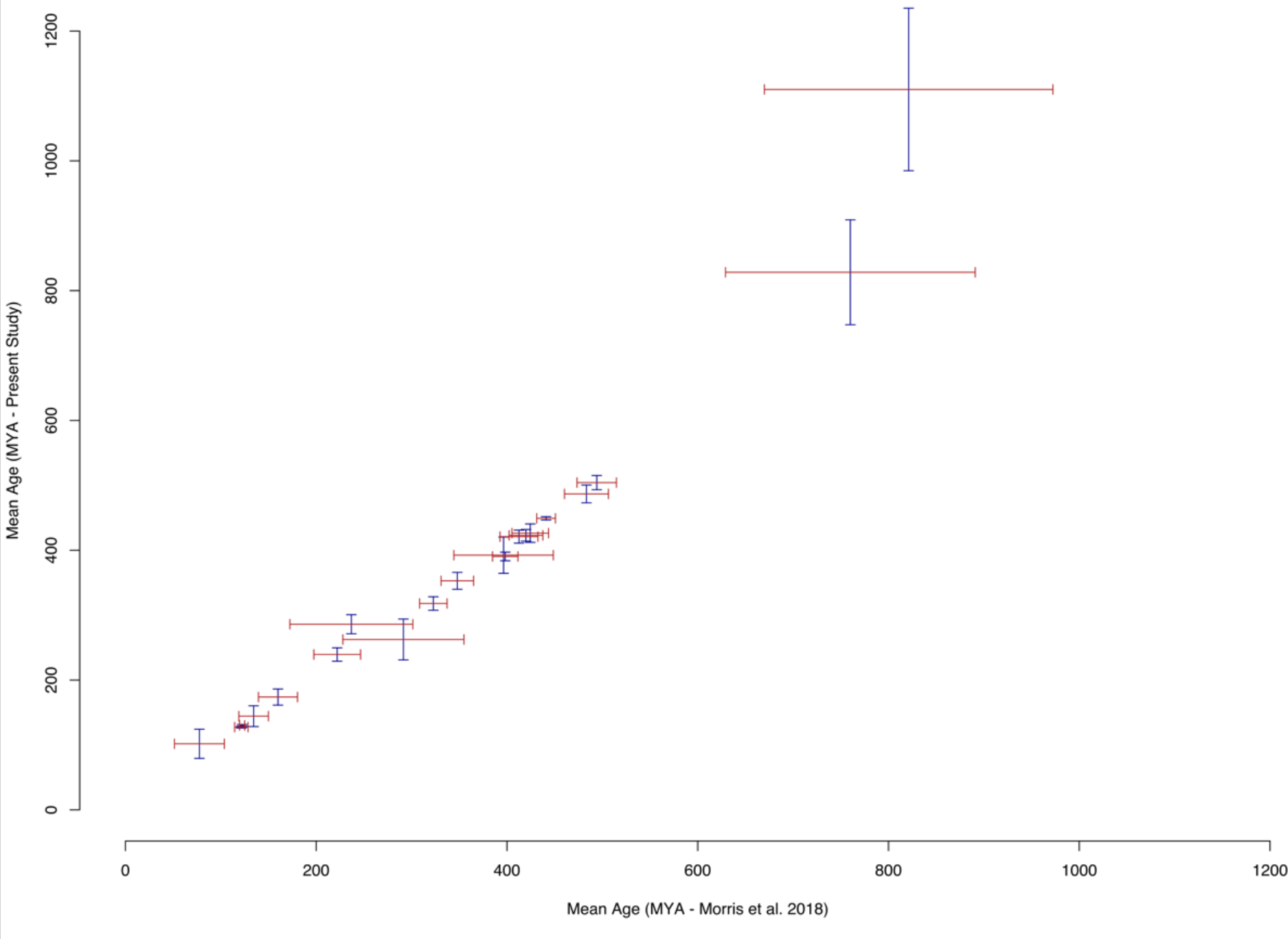
A comparison of divergence time estimates between the current study and the closest benchmark, that of Morris et al 2018. Points on each axis are centred on the mean age, with the 95% highest posterior density represented by the width of the bars.

**Supplementary Figure 10.**
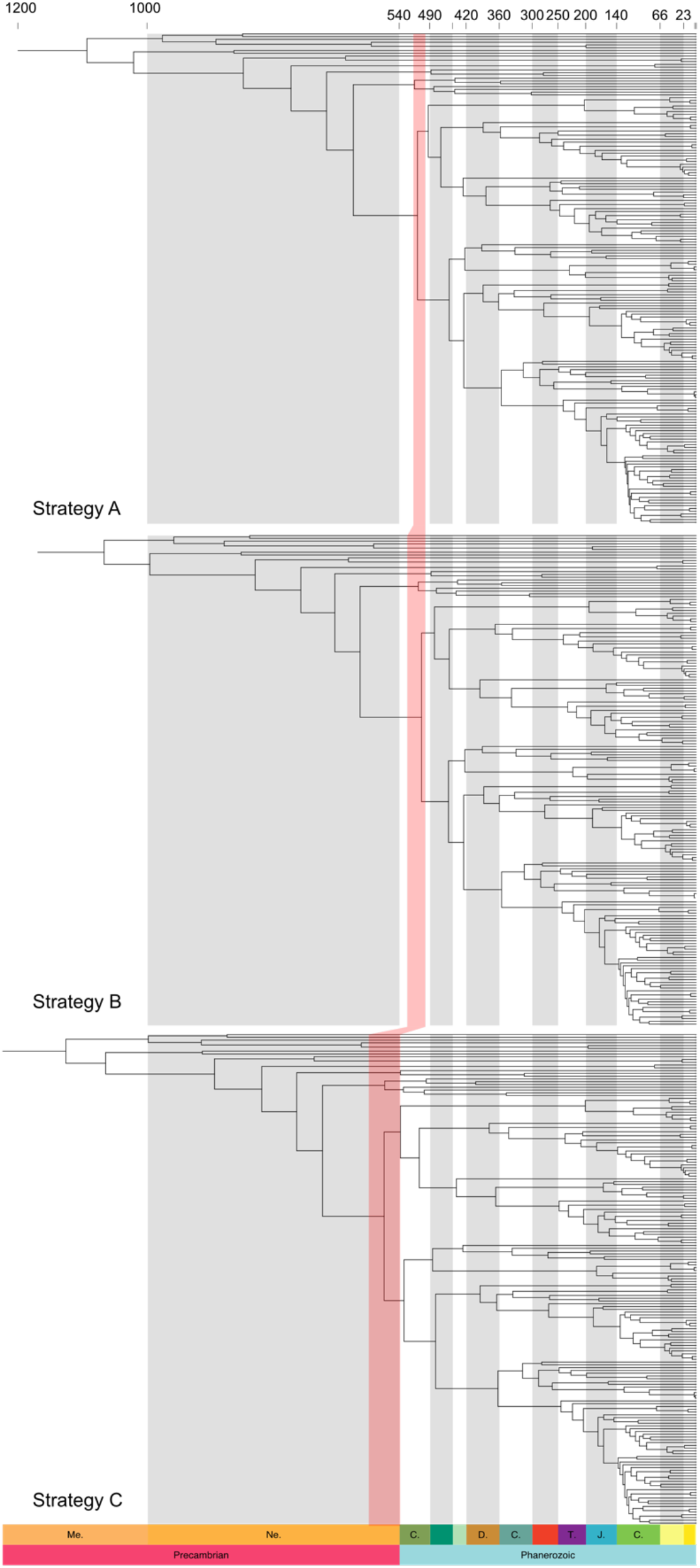
The effect of alternative calibration strategies on the age of crown group embryophytes. Calibrations were altered by variously relaxing maximum age calibrations on the age of embryophytes (Strategy B) and embryophytes and tracheophytes (Strategy C). The width of the red band across the phylogenies represents the 95% HPD interval.

**Supplementary Figure 11.**
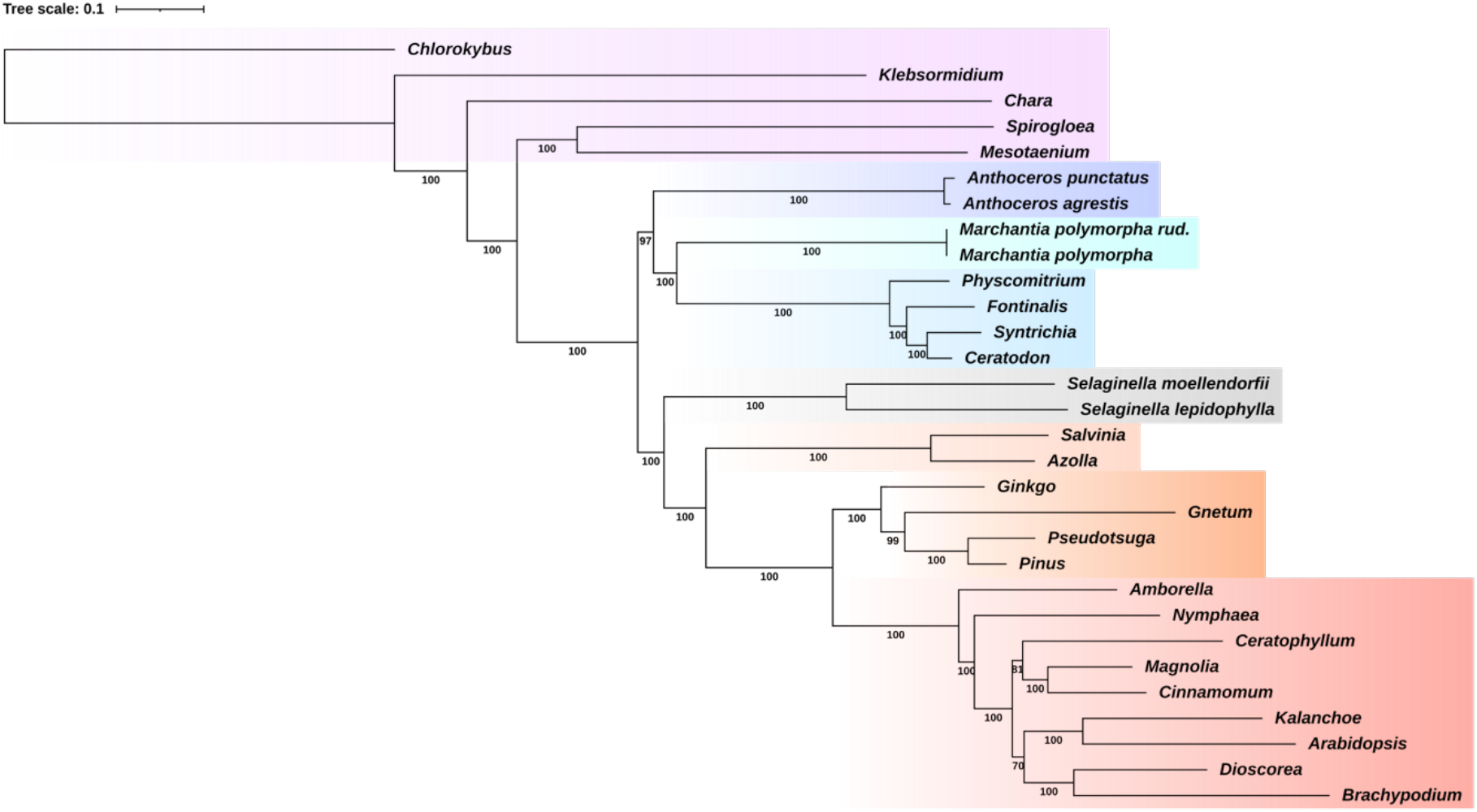
Phylogenetic tree of embryophytes used for ancestral reconstruction. The maximum likelihood tree was inferred from an alignment comprised of 30 species, 185 single copy orthologs and 71855 sites under the LG+C60+G4+F model in IQ-Tree (Nguyen *et al.,* 2015). Bootstrap support values are placed underneath the corresponding branches. The branch lengths are proportional to the number of average number of substitutions.

**Supplementary Figure 12.**
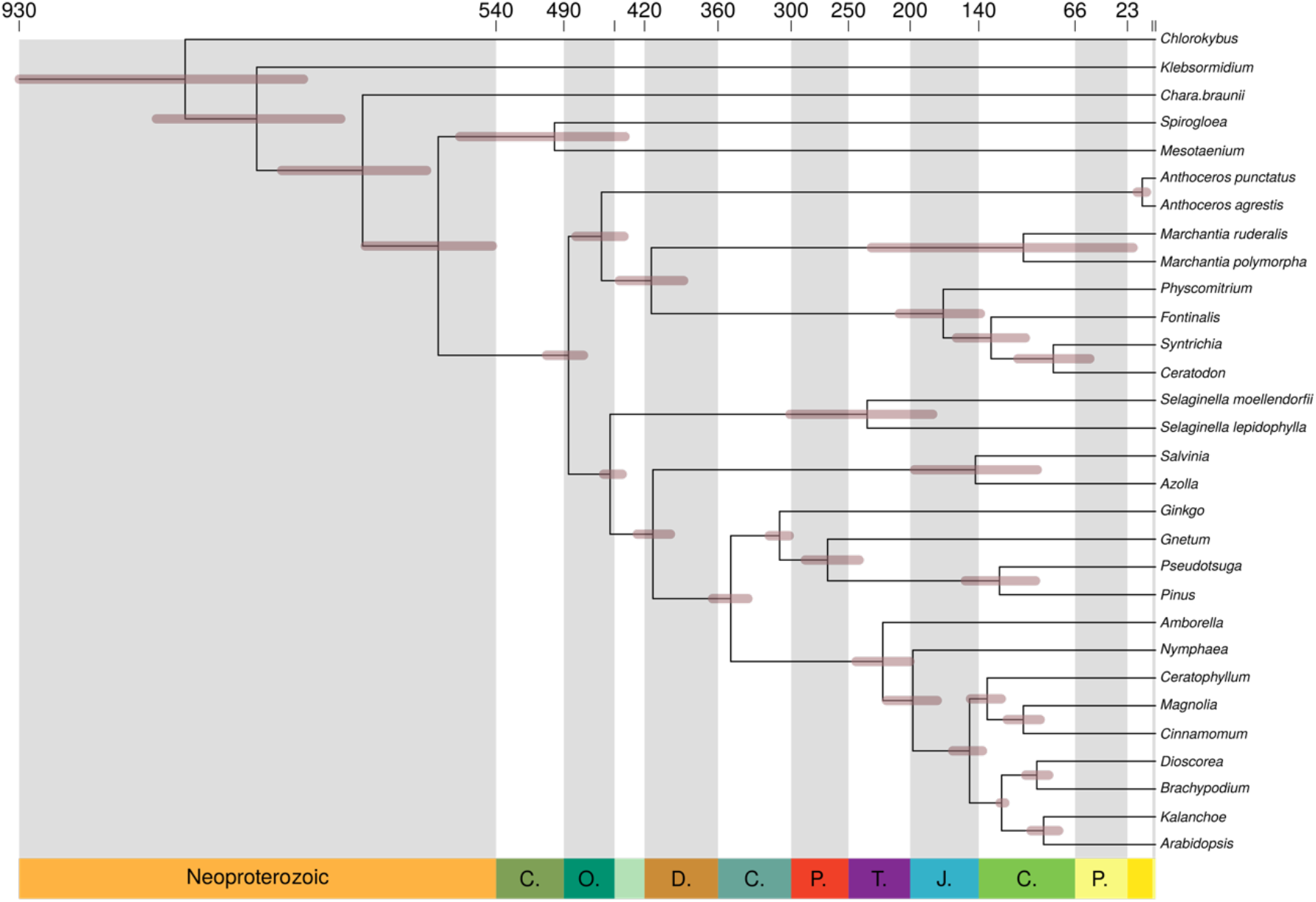
Time scale of embryophyte tree used for ancestral reconstruction. The amino acid sequence alignment was combined with 18 fossil calibrations subsampled from the list of calibrations from the transcriptomic analysis. The normal approximation method was used in MCMCTree, with branch lengths estimated under the LG+G4 model. Calibrations were modelled as a uniform distribution between a minimum and soft maximum. 95% highest posterior density (HPD) width is shown as bars on node.

**Supplementary Figure 13.**
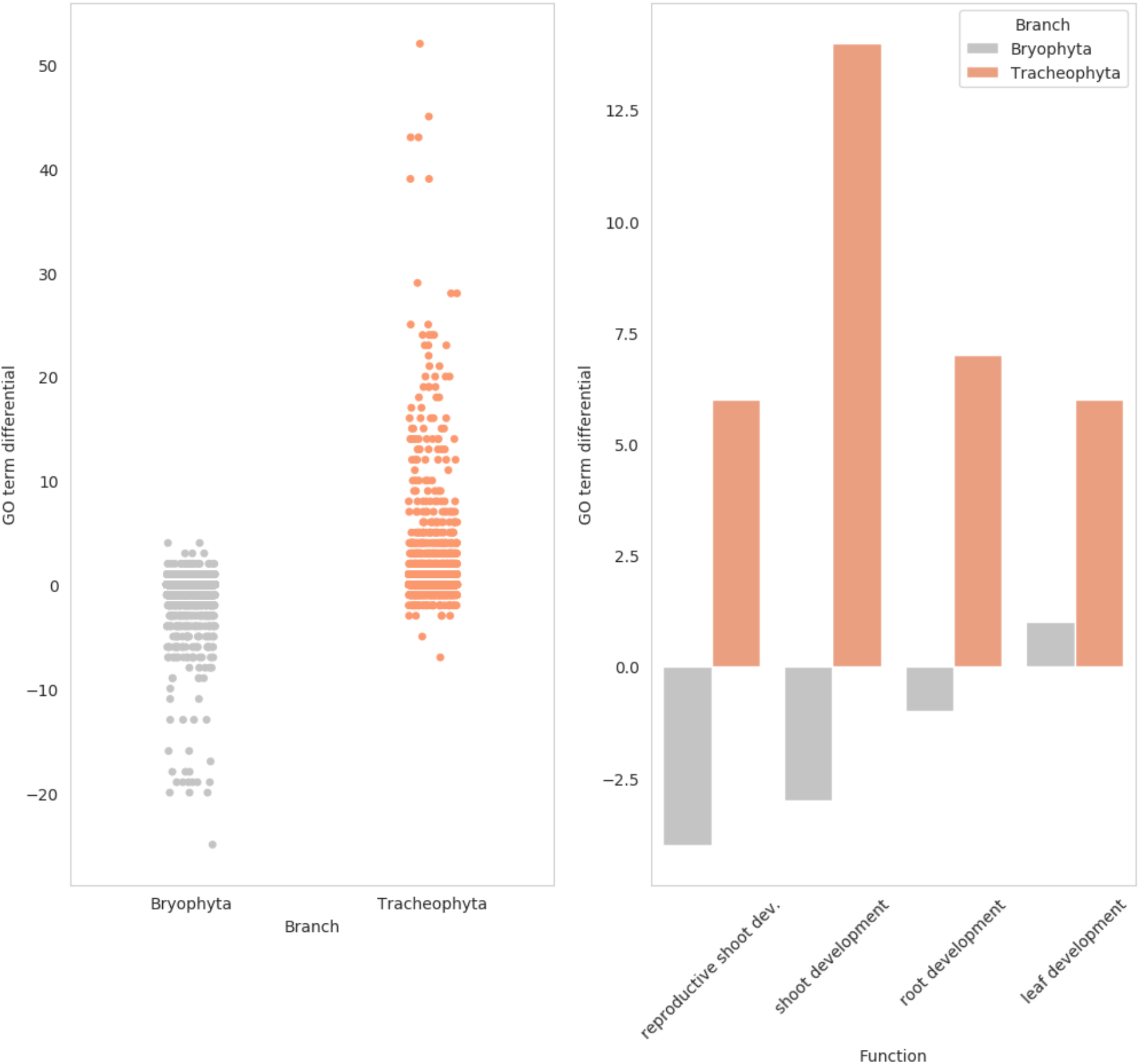
Functional annotation of gene family changes between the ancestral embryophyte, bryophytes and tracheophytes. Left, overall change in GO term frequency between the ancestral embryophyte and the ancestral bryophyte/tracheophyte. GO terms on average become less frequent in bryophytes. Right, change in the frequency of specific GO terms between the ancestral embryophyte and the ancestral bryophyte/tracheophyte. Bryophytes have a reduction in gene families associated with shoot and root development, whilst we see an increase in gene families associated with these GO terms in the tracheophyte ancestor.

**Supplementary Figure 14.**
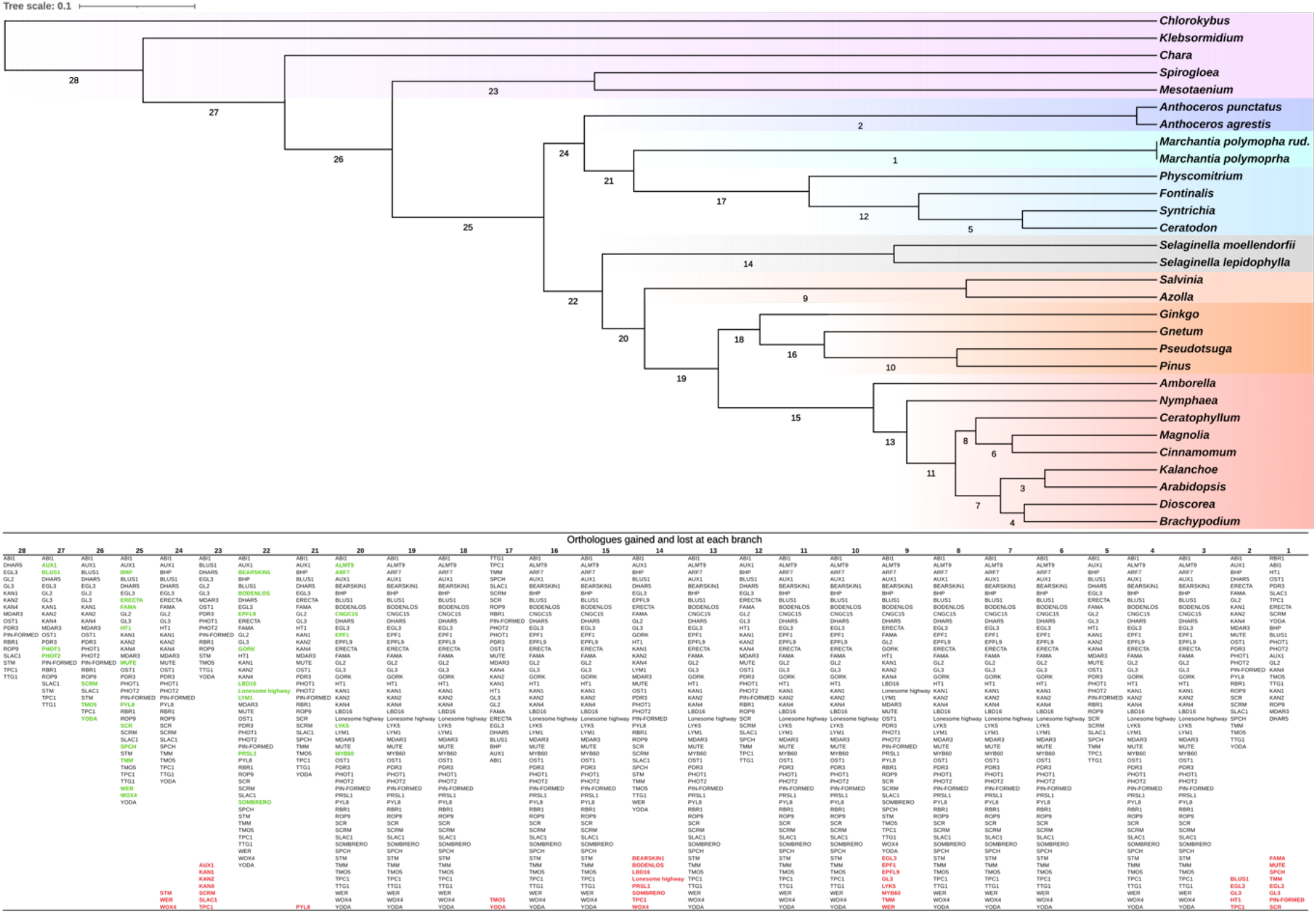
Gain and loss of orthologous gene family clusters across the embryophyte tree. Above, maximum likelihood time calibrated tree, with labels assigned to each branch. Below, the labels correspond to a list of the key orthologous gene family clusters present at each branch. Fifty critical genes for *Arabidopsis,* stomata, vasculature and symbiosis related function/development were used to illustrate the gain and loss of orthologous gene family clusters across the embryophyte tree. Gene families are labelled according to their characterised representatives; for example, the “SLAC1” gene family is the family that contains *Arabidopsis* SLAC1. Gene family clusters that originated on the branch are coloured in green, whilst ones that are lost are coloured red and placed at the bottom of the list. The order of the branches in the table is not consistent with the path of embryophyte evolution i.e. branch 1 is not a child of branch two, but is a child of branch 21, with gain and loss following the phylogenetic tree and not the numerical order of the branch labelling.

**Supplementary Figure 15.**
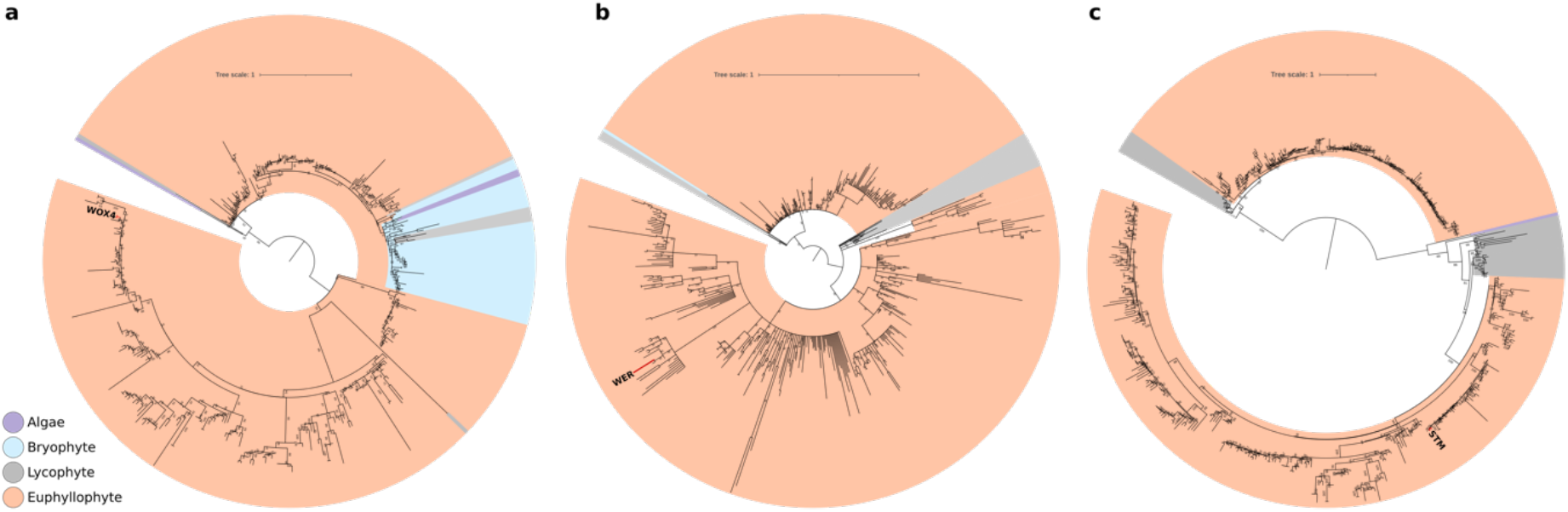
Phylogenetic trees of key losses on the bryophyte stem. Gene trees were constructed from homologous sequence families determined via a BLAST. Gene families were aligned with MAFFT and trimmed with BMGE. Gene trees were inferred in IQ-Tree, under the best fitting phylogenetic model determined by Bayesian Inference Criterion. The trees are rooted in accordance with the species tree displayed in Fig. 1. **a,** Gene tree of WOX4 (*Arabidopsis*) and homologous sequences. The tree suggests that an ancestral WOX gene duplicated prior to the divergence of bryophytes and tracheophytes. Bryophytes have retained one of the WOX genes, but have appeared to have lost the duplicate that went on to become WOX4 in *Arabidopsis,* consistent with the ALE analysis. **b,** Gene tree of WER/WEREWOLF (*Arabidopsis*) and homologous sequences. One bryophyte (*Atrichum)* has retained a WER homolog, thus dating the duplication of the ancestral WER gene to prior to the divergence of bryophytes and tracheophytes. Bryophytes have subsequently lost the duplicate that went on to become WER in *Arabidopsis,* consistent with the ALE analysis. **c,** Gene tree of STM (*Arabidopsis*) and homologous sequences. The ancestral STM gene underwent a duplication, with an extant alga (*Mougeotia*) retaining one of these STM paralogs, thus dating the duplication of the ancestral STM gene to prior the divergence of bryophytes and tracheophytes. Bryophytes have subsequently lost both these paralogs, including the gene that went on to become WER in *Arabidopsis,* consistent with the ALE analysis.

## Supplementary Methods

### 1. Selecting gene families for phylogenetic analyses

Whole genome duplications are prevalent in embryophytes (Clark & Donoghue, 2018), leading to multiple gene copies and paralog-dominated gene families. As a result, single copy orthologous gene families used to construct concatenations/supermatrices are difficult to identify – especially when sampling taxon-rich datasets. We employed OrthoFinder 2.0 (Emms & Kelly, 2020) to infer single copy orthologs from our dataset of 177 species of plants and algae. No single copy orthologous gene families were inferred. In previous studies single copy orthologous gene families have been manually curated (Puttick *et al.,* 2018). We chose to develop a systematic algorithm identifying the gene families closest to single copy (Supplementary Fig. 2). The filtering method employed ensured only high-quality gene families comprised the final alignment – with only 2/177 species being represented by more than 50% gaps. The algorithm is split into two scripts *prem3.py* and *super_matrix_2.py* - prior to calling *super_matrix_2.py* the gene families need to be aligned and trimmed. This can be done prior to use, however, aligning all orthogroups prior to filtering would require considerable time and computational resources.

### 2. Transcriptome Assembly

Raw sequencing reads were downloaded from the Sequence Reads Archive (SRA) and trimmed using trimomatic (Bolger et al. 2014) under default settings to remove adapter sequences and trim low-quality bases. Transcriptomes were de novo assembled in Trinity (Haas et al. 2013). Transcriptomes were filtered using the evidentialGene pipeline (Gilbert 2013) to remove isoforms and translate transcripts into amino acid sequences.

### 3. Root inference with Amalgamated Likelihood Estimation (ALE)

ALE models gene duplication, transfer and loss (DTL) to assess the likelihood of gene families under differing roots. Due to our phylogenetic dataset (Supplementary Table 1) containing both genomes and transcriptomes, the incomplete nature of the transcriptomes would be difficult to model accurately. In Supplementary Figure 4, the fraction missing of each lineage is displayed (detailed in Supplementary Table 1) – for many lineages more than half the BUSCOs are absent. As a result, we decided to curate a new dataset comprising of only high quality and more complete genomes. The dataset comprised of 24 species, with each of the seven major lineages of embryophytes being represented by at least one genome (Supplementary Fig. 5). 23 out of 24 species had more than 70% BUSCO completion, with most of the data being more than 80% complete. Moreover, the gene family clusters inferred from the high-quality dataset had greater species representation, and thus would be more informative of the root (Supplementary Fig. 6). There were also fewer large clusters with poor species representation in the high-quality dataset i.e., less gene families that have either undergone myriad duplications and/or had multiple isoforms of genes from sequencing anomalies. As a result, we only took forward the high-quality dataset into further analysis.

The high-quality gene families were analysed by ALEml – ALEml is time-consistent with the fossil-calibrated species tree and avoids time-inconsistent gene transfers (that is, gene transfers travelling back in time). We ran ALEml, on 12 rooted and dated embryophyte trees – with an AU test suggesting three credible roots: hornworts, moss, and monophyletic bryophytes (Supplementary Table 3). The credible roots were all in the same region of the tree, and the likelihood of the roots decreased with increasing distance from the root region, unlike the undated analysis, generating greater confidence in the analysis. Moreover, analysis of the DTL signal (Supplementary Fig. 7), revealed that the signal was stable for this set of credible roots.

We also employed two additional techniques to test the root of embryophytes to ensure confidence in our topology for the reconstruction of ancestral states. Firstly, we inferred constrained maximum likelihood trees from the concatenation to be in accordance with the three credible roots selected by ALE i.e., hornworts were constrained to group with algae at the base of the tree, and thus be the root of embryophytes. The constrained trees were inferred in IQ-Tree (Nguyen *et al.,* 2015), under the LG+C60+G4+F model. The likelihood of the three constrained trees was then assessed with an AU test (Shimodaira, 2002) – rejecting both moss and hornworts as earliest diverging lineages of embryophytes (Supplementary Table 4). Secondly, we ran STRIDE (Emms & Kelly, 2017), an alternative rooting software which employs duplication events to determine the root. STRIDE in combination with OrthoFinder 2.0 (Emms & Kelly, 2020), reconciled 23145 orthogroups, placing the root between bryophytes and tracheophytes (Supplementary Fig. 8), concurrent with the out-group rooting analysis displayed in Fig. 1.

### 4. Ancestral gene content reconstruction

In addition to reconstructing the gene content of the ancestral embryophyte, we included several algal species that had highly complete genomes; the inclusion of algae allowed us to investigate gene content evolution in the closest algal relatives of embryophytes. The inclusion of algae also enabled us to investigate further the gene loss proposed to underpin the reduced body plan of the embryophyte sister lineage, Zygnematophyceae. In addition to algae, a subspecies of *Marchantia polymorpha (ruderalis*), was added to the dataset, allowing us to display values for DTL and copy number for the internal node defining liverworts. Single copy orthologs were determined using the same method as the main analysis (Supplementary Fig. 2) – 185 single copy gene families were inferred, leading to an alignment of 71855 sites. A maximum likelihood tree (Supplementary Figure 11) was inferred from the dataset under the LG+C60+G+F model in IQ-Tree (Nguyen *et al.,* 2015). The tree was rooted in accordance with the previous analysis. Again, even with this reduced dataset, we recover monophyletic bryophytes with high support (97% Bootstrap support). Due to ALEml requiring a dated tree for reconstruction of ancestral gene content, a time scale was inferred using a subset of the calibrations from the transcriptomic analysis (Supplementary Fig. 13). Divergence time analyses were performed as described for the transcriptomic analysis.

Gene family clusters were inferred using the same methods as the rooting analysis. The pipeline yielded 20822 gene family clusters with 1000 bootstrap samples for each family. The gene families were reconciled with the rooted species tree, using the ALEml algorithm with the fraction missing of each genome also incorporated. The presence and absence of each gene family across the tree was inferred using the *Ancestral_reconstruction_copy_number.py* python script – we determined that a gene family was present if it had more than 0.5 copies at the specified branch i.e., was present at the branch in more than 50% of the reconciliations. The summed number of DTL evolutionary events was determined with *branchwise_number_of_events.py*. In addition to this, we scaled the number of events by the branch length of the time scale analysis, to determine the rate of DTL for important sections of the tree. The number of DTL evets and the summed gene copy number for each branch of the tree were calculated (Supplementary Table 6). The presence and absence of each gene family cluster across the embryophyte tree, and whether an *Arabidopsis* gene is present in said cluster is displayed in Supplementary Table 6.

To functionally annotate the gene families, the consensus sequence of each gene family was determined using HMMER3 (Eddy, 2011). This process summarises the gene family into a single sequence most indicative of the ancestral state. The consensus sequences were functionally annotated using eggNOG-mapper 2 (Huerta-Cepas *et al.,* 2017). The gene families present at each branch were determined as described above, and the functional annotation for their corresponding consensus sequence were retrieved. The frequency of GO terms associated with the consensus sequences present at each branch were calculated using the custom python script *make_go_term_dictionary.py.* In doing so, the change in the frequency of GO terms could be tracked from node to node, allowing us to identify changes in function across the embryophyte tree. Changes in the frequency of GO terms from the embryophyte ancestor to bryophytes, and to tracheophytes was calculated – this is termed the GO term differential i.e., if there was a frequency of 10 for the GO term “Shoot Development” at the ancestral embryophyte branch, but only a frequency of 6 for the bryophyte branch, the GO term differential would be -4. The GO term differentials are presented in Supplementary Fig. 14: on the left-hand side, all GO term differentials are plotted, i.e., the change in every GO term differential between the ancestral branch and the subsequent tracheophyte and bryophyte branch is plotted. Overall, there has been a reduction in the frequency of GO terms in bryophytes in comparison to the ancestral embryophyte, suggesting there has been a wide scale genomic reduction in this linage. This was also suggested by the analysis of DTL rates which suggested a vast amount of loss on this branch. On the right, changes in select GO terms/functional processes are highlighted. Reduction in the frequency of GO terms associated with reproductive shoot and vegetative shoot development are observed between the ancestral embryophyte and the bryophytes, whilst the inverse is true for tracheophytes (Supplementary Tables 7, 9 and 10).

One limitation of GO term analysis is that differences on the bryophyte and tracheophyte stems might reflect, in part, differences in the completeness of functional annotation for the extant members of the two clades. To investigate the broad-scale patterns identified in the GO term analysis in more detail, we investigated the evolutionary histories of 50 key genes involved in *Arabidopsis* stomatal function/development (Supplementary Table 11), vasculature development and symbiosis/defence (Supplementary Fig. 15). We were able to determine where the gene families that these genes belong to originated, were present, and were lost across the tree. Many of the orthologous gene clusters that harbour critical constituents of the stomatal development pathway such as SPCH/MUTE/FAMA, TMM, and ERECTA originated on the ancestral embryophyte branch, suggesting that stomata evolved at this point. Moreover, orthologous gene families of master-regulators of stomatal function such as HT1 and OST1 were also present. In addition to this, gene families which contain key *Arabidopsis* vasculature genes such as WOX4 and WER (MYB66) originated on the branch defining Embryophytes – highlighting a possible increase in vasculature complexity. Interestingly, our analysis suggests that these gene families were lost on the branch defining bryophytes (Supplementary Fig. 15), suggesting a reduction in vasculature complexity in this lineage.

To further assess the losses of WOX4, WER and STM on the branch defining bryophytes, manually curated phylogenetic trees were constructed for their respective gene families (Supplementary Figure 16). The gene trees were inferred by first undertaking a BLAST search of the 177 species dataset to retrieve homologous sequences. The homologues were then aligned with MAFFT and trimmed with BMGE. Phylogenetic trees were inferred from the alignments in IQ-Tree under the best fitting model determined by BIC. The trees were then rooted in accordance with the previous analysis (Fig. 1). It appears for all three genes that the origination of the orthologous gene families in which they belong predated the divergence of bryophytes from tracheophytes. In bryophytes, the ancestral gene of WOX4, WER and STM was subsequently lost, but it was retained in tracheophytes. Additionally, we see wide scale loss of gene families which harbour critical stomata and vasculature associated genes on the branches that define hornworts and liverworts (Supplementary Fig. 16). The loss of stomatal- associated gene families in these lineages is congruent with the findings of Harris *et al.,* (2020). Presently, the number of available genomes for non-flowering plants is limited and is a potential source of bias for estimating ancestral genome content in deeper nodes of the tree. We expect, with the increasing volume of sequence data and projects specifically targeting non-flowering plants (e.g. 10000 genome project, Sphagnum genome project), that it will be possible to develop a still more detailed reconstruction of the ancestral embryophyte.

### 5. Molecular clock estimates of the timing of land plant evolution

The age of land plants has become a recent controversy, with some studies favouring a far older (pre-Cambrian) origin of crown embryophytes (Hedges et al. 2018; Su et al. 2021), while others limit the origin of land plants to the late Cambrian. These differences reflect diverging approaches to the implementation of maximum age constraints derived from the fossil record. Previous studies have employed a maximum age derived from the earliest record of cryptospores. Cryptospores are spores that possess certain characteristics of embryophyte spores, but are sufficiently different to make their assignment to any extant clade difficult. However, the possibility of an embryophyte affinity cannot be disregarded. Cryptospores possess a similarly high preservation potential to the spores of extant land plants, and indeed from the Ordovician onwards there is a good fossil record of spores that are assigned to extant land plant lineages. This provides the basis of the maximum age calibration, following the reasoning that if land plant spores did exist prior to the Cambrian, they would likely be preserved.

To address recent discussions about the value of these maxima, we implemented two additional calibration strategies. In the first (Strategy B), we relaxed the late Cambrian maximum age calibration on crown embryophytes (and all internal nodes that share the same calibration). This was achieved by setting an arbitrary ancient maxima of 1 Ga. The maximum age on tracheophytes (derived from similar reasoning regarding the presence of trilete spores), was retained. In the second strategy, we relaxed the maximum calibration on tracheophytes also to 1 Ga, provided no effective maxima on the major lineages of land plants. Analyses were performed as described in the methods of the main text and the impact on the age of crown embryophytes across calibration strategies is compared in Supplementary Figure 10.

### 6. Fossil Calibrations for Molecular Clock Analyses

#### C_1MRCA Viridiplantae/Chloroplastida

***Fossil Taxon and Specimen:*** *Proterocladus antiquus* [VPIGM-4762, deposited at Virginia Polytechnic Institute Geoscience Museum], from the lower Nanfen Formation in North China [1].

***Phylogenetic justification:*** Macrofossils from the Nanfen Formation, Northern China, have reconstructed an erect, epibenthic, multicellular alga. Tang *et al.* [1] argue that the overall morphology of *P. antiquus* is comparable to extant members of Ulvophyceae, yet no diagnostic features are described. As such, *P. antiquus* can conservatively be used to provide a minimum calibration on the origin of Viridiplantae.

***Minimum age:*** 940.4 Ma

***Maximum age:*** 1891 Ma

***Age justification:*** The Nanfen Formation is positioned within the Xihe group in southern Liaoning Province, North China [2]. The Nanfeng Formation has not been dated directly. The age of the underlying Diaoyutai Formation can only be constrained by detrital zircons, the youngest population of which has been dated to 1056 Ma ± 22 Ma [3]. A sill in the overlying Qiaotou Formation, which overlies the Nanfeng Formation, has a zircon secondary-ion mass spectrometry U-Pb age of 947.8 Ma ± 7.4 Myr [4]. Thus, the minimum age of the Nanfeng Formation can be established on the minimum age interpretation of this sill viz. 940.4 Ma, although it is presumably significantly older. The soft maximum age is established following Morris *et al.* [5] and is based on the earliest record of eukaryotes, where despite the presence of simple eukaryotes, there is no evidence of complex or multicellular eukaryotes such as green algae.

***Discussion:*** *Proterocladus* is determined as a crown group member of green algae based on the cellular differentiation, presence of a holdfast, inferred siphonocladous nature, and lack of an outer sheath. The absence of an outer sheath and siphonocladous nature precludes an affinity with stigonematalean cyanobacteria. A fungal affinity was also rejected based on the structure of the septae, lack of mycelium-like branching and the structure of the reproductive organs. The septation in *Proterocladus* also suggests that it is not similar to the xanthophycean alga *Vaucheria* or the rhodophyte *Griffithsia*. Nie *et al.* consider a Mesoproterozoic specimen identified as *Pterospermella* as a member of Prasinophyceae [6, 7], however the lack of clearly described structures prevents their unequivocal assignment to the green algae [5].

#### C_2: Chlorophyceae – Prasinophyaceae

***Fossil taxon and specimen:*** *Palaeocymopolia silurica*

***Phylogenetic justification:*** Following Morris *et al.* [5]

***Minimum age:*** 438.3 Ma

***Maximum age:*** 1891 Ma

***Age justification***: Following Morris *et al.* [5]

#### C_3: Steptrophyta: “Streptophytes” - Embryophytes

***Fossil taxon and specimen:*** *Tetrahedraletes cf. medinensis*

***Phylogenetic justification:*** Following Morris *et al.*, [5]

***Minimum age:*** 469 Ma

***Maximum age:*** 1891 Ma

***Age justification***: Following Morris *et al.* [5]

#### C_4: Embryophyta: Bryophyta – Tracheophyta

***Fossil taxon and specimen:*** *Cooksonia cf. pertoni*

***Phylogenetic justification:*** Following Morris *et al.* [5]

***Minimum age:*** 426.9 Ma

***Maximum age:*** 515.5 Ma

***Age justification***: Following Morris *et al.* [5]

#### C_5: Marchantiophyta: Jungermanniopsida - Marchantiopsida

***Fossil taxon and specimen:*** *Metzergiothallus sharonae* [NYSM 17656, Paleontology Collection of the New York State Museum, Albany, USA], from the Sample CHD-4.6, Plattekill Formation of the eastern Hamilton Group, Givetian (late Middle Devonian) at Cairo Highway Department quarry, just south of New York State Route 145, 3.22 km NW of Cairo, Greene County, New York (42.32° N and 74.04°W, NAD 83).

***Phylogenetic justification:*** Morris *et al.* [5] identified *Riccardiothallus devonicus* as the oldest liverwort but its compression-preservation precludes the identification of anatomical characters that might corroborate a liverwort affinity beyond its general form [8]. *Metzergiothallus sharonae,* preserves anatomical detail including putative oil body cells [9, 10], bifurcating thalloid habit with an axial costa, dorso-ventral cellular differentiation, ventral ribbon-like rhizomes, and associated sporophyte that has a multivalve structure [11]. These characteristics support the interpretation of *Metzergiothallus sharonae* as a crown group liverwort, as supported by a cladistic analysis of morphological characters [12]. Flores *et al.* placed *M. sharonae* as sister to the extant genus *Metzgeria*, albeit with very low bootstrap support (50%). As such, we conservatively consider *M. sharonae* as a member of the Jungermanniopsida and thus capable of calibrating the divergence between Jungermanniopsida and Marchantiopsida.

***Minimum age:*** 377.7 Ma

***Maximum age:*** 515.5 Ma

***Age justification***: Attempts to resolve the age of the Plattekill Formation at Cairo Quarry through palynostratigraphy have proven unsuccessful due to poor preservation [13], however, the age of the Plattekill Formation has been established elsewhere. Willis and Bridge [14] placed the Plattekill and overlying Manorkill Formations in the mid *lemurata magnificus*– mid *optivus triangulatus* zones of Richardson and McGregor [15], indicating a latest Eifelian- early Givetian age [16]. In the absence of better stratigraphic constraint, this allows us to establish a minimum constraint on the age of the Plattekill Formation based on the Givetian-Frasnian boundary, dated to 378.9 Ma ± 1.2 Myr [16], thus 381.7 Ma.

#### C_6: Anthocerotophyta: Anthocerotaceae

*Fossil taxon and specimen: Anthoceros sp.* (spore type A) [BA Pb Pm 5034, Museo Argentino de Ciencias Naturales] from the Albian Anfiteatro de Ticó Formation, Baqueró Group, Santa Cruz Province, Argentina

*Phylogenetic justification:* This spore was assigned to the extant genus *Anthoceros* based on the similar morphology [17, 18].

*Minimum age:* 116.59 Ma

*Maximum age:* 515 Ma

*Age justification:* The spores were recovered from the Anfiteatro de Ticó Formation [18], which is overlain by the Bajo Tigre Formation, constrained based on U/Pb dating to 116.85 ± 0.26 Ma [19]. Thus, a minimum age of 116.59 Ma is proposed.

#### C_7: Anthocerotophyta: Notothyladaceae

*Fossil taxon and specimen: Notothylites nirulai* from the Deccan Intertrappean beds of Mohgaon-Kalan, Madhya Pradesh, India [20].

*Phylogenetic justification:* The overall morphology has a similar thallus size, sporophyte size and elater shape to extant *Notothylas*, as well as lacking stomata on the sporophyte [17].

*Minimum age:* 65.81 Ma

*Maximum age:* 515 Ma

*Age justification:* The age of the Intertrappean beds at Mohgaon-Kalan was determined to be Maastrichtian on the palynological assemblage being characterised as Maastrichtian and the magnetostratigraphy indicating the flows to be of C30N (Maastrichtian) [21]. Thus, a minimum age based on the base of the C30N polarity chron of 66.31 ± 0.5 is proposed [22].

#### C_8: Anthocerotophyta: *Phaeomegaceros*

*Fossil taxon and specimen: Phaeoceros sp.* from the Uscari Formation, Costa Rica *Phylogenetic Justification:* Though originally assigned to the extant genus *Phaeoceros* [23], the spore is highly similar to the extant genus *Phaeomegaceros* in size and morphology (Villareal and Renner)

*Minimum age:* 13.82

*Maximum age:* 515 Ma

*Age justification:* Material was deposited in the Uscari Formation in the Limon Basin of Costa Rica. Analysis of planktonic formanifera identified an overlap of *Orbulina universa* and *Globorotalia foshi peripheroronda*, placing the the Formation in the early Middle Miocene (formaniferal zones N9 to N10) [24]. Following Hilgen et al. [25], the base of the N10 formaniferal zone correlates with the M7 zone and the base of the Langhian and thus a minimum age of 13.82 Ma is derived.

#### C_9: Marchantiophyta: Pelliidae - [Jungermannidae + Metzgeriidae]

***Fossil taxon and specimen:*** *Pellites hamiensis* [SDL-98-4-91, Institute of Paleontology and Stratigraphy of Lanzhou University, Gansu Province, China] from the Sandaoling coal mine, Hami City, Xinjiang Uygur Autonomous Region, China.

***Phylogenetic justification:*** The gross morphology and anatomy of the thalli support a placement within Pelliaceae [26], including ribbon-like segments, thinning wings, a unistratose margin, a conspicuous costa with elongate cells and thick rectangular cells near the margin. The absence of reproductive structures led the authors to assign the samples to the fossil genus *Pellites* rather than *Pellia* [27].

***Minimum age:*** 167.0 Ma

***Maximum age:*** 515.5 Ma

***Age justification:*** Material was collected from the Xishanyao Formation of the Sandaoling coal mine, which overlies the lower Jurassic Sangonghe Formation and underlies the Palaeogene Shansan Group strata [28]. Based on biostratigraphy and plant assemblages, an Aalenian to Bajocian age is proposed for the Xishanyao formation [29]. Thus, we propose a minimum age according to the upper boundary of the Bajocian, 168.2 ± 1.2 Ma [30].

#### C_10: Marchantiophyta: Sphaerocarpales – Marchantiales

***Fossil taxon and specimen:*** *Marchantites cyathodoides*

***Phylogenetic justification:*** Following Morris *et al.* [5]

***Minimum age:*** 227.6 Ma

***Maximum age:*** 515.5 Ma

***Age justification***: Following Morris *et al.* [5]

#### C_11: Marchantiophyta: Ricciaceae – Conocephalaceae

***Fossil taxon and specimen:*** *Ricciopsis ferganica* [FG 596/X/721 in the Palaeontological collection of the Geologisches Institut, Technische Universität Bergakademie Freiberg, Germany] from the village of Madygen, southwest Kyrgyzstan, Central Asia

***Phylogenetic justification:*** *R. ferganica* was assigned to the fossil genus *Ricciopsis* (Ricciaceae) [31] based on resemblance to the thalloid floating forms of the extant genus *Riccia.* No diagnostic features are available for the genus, however a flattened, small branching thallus and the presence of a longitudinal median furrow or groove on the dorsal side of the thallus is characteristic of Ricciaceae [32]. Further, a cladistic analysis of morphological characters supported the placement of *Ricciopsis* as sister to the extant genus *Riccia* [12].

***Minimum age:*** 227.6

***Maximum age:*** 515.5 Ma

***Age justification***: Material was collected from the lower and middle parts of the uppermost lacustrine unit at Urochishche Madygen. The Madygen Lagerstätten is deemed to be Ladinian-Carnian (Middle to Late Triassic) in age based on comparative studies of the flora [33] and insect assemblages [34]. Thus, we propose a minimum age based on the top of the Carnian of 227.3 ± 0.1 Ma [30].

#### C_12: Marchantiophyta: Jungermanniidae – Metzgeriidae

**Fossil taxon and specimen:** *Cheirorhiza brittae* [515-38a, Far East Geological Institute, Far East Branch of the Russian Academy of Science] from the Bureja basin, Russia

**Phylogenetic justification:** *C. brittae* has been compared to extant members of both Jungermanniales and Porelliales on the basis of morphology [35, 36], and so we follow Heinrichs et al. [37] and treat it as a stem lineage member of Jungermanniidae.

**Minimum age:** 143.1

**Maximum age:** 515.5

**Age justification:** Material was collected from the Talyndzhan formation of the Bureja basin. The Talyndzhan formation was initially determined to be of Callovian or Callovian- Oxfordian age based on floral assemblage. This has been revised in light of palynological evidence and the top of the Talyndzhan is believed to correspond to the Jurassic-Cretaceous boundary [38, 39]. Thus, a minimum age based on the upper boundary of the Jurassic 143.1 [30], and so 144.9 Ma, is proposed.

#### C_13: Marchantiophyta: Jungermanniales – Porellales

***Fossil taxon and specimen:*** *Diettertia montanensis* [no. 281 of the University of Montana Paleontological Collection (UMPC)] from the Missouri river, Montana, USA.

***Phylogenetic justification:*** *Dietteria montanensis* has been assigned as a member of Jungermanniales stem lineage based on its overall morphology and anatomy, including complicate-bilobed, subtransverely inserted leaves, stems with a small merophyte with narrow elongated cortical cells [37, 40].

***Minimum age:*** 112.8 Ma

***Maximum age:*** 515.5 Ma

***Age justification*:** Material was collected from the Kootenai Formation, Great Falls, Montana. Fossil assemblages, particularly molluscs, ostracods and charophyte algae indicate an Aptian age for the formation and an Albian age for the overlying Blackleaf formation [41, 42]. Thus, a minimum age is derived from the Aptian-Albian boundary, estimated at 113.10 ± 0.3 Ma, of 112.8 Ma.

#### C_14: Marchantiophyta: Porellineae – [Radulineae + Jubulineae

***Fossil taxon and specimen:*** *Gackstroemia cretacea* [AMNHBU ASJH-1, American Museum of Natural History, New York], from Myanmar [43].

***Phylogenetic justification:*** Heinrichs *et al.* [43] placed *G. cretacea* within the Lepidolaenaceae based on a sterile gametophyte, including incubous foliation, complicate bilobed leaves, splitting to a dorsal lobe and a ventral *Frullania*-type water sac. Based on the presence of bifurcate underleaves, the fossil was assigned to the extant genus *Gackstroemia.* Lepidolaenaceae are placed within the Porellineae, and so *G. cretacea* can calibrate the divergence of *Porella* (Porellineae) from *Radula* (Radulineae) and Frullania (Jubulineae).

***Minimum age:*** 98.17

***Maximum age:*** 515.5 Ma

***Age justification:*** The age of Burmese amber has long been contested, with estimates ranging between the Miocene and Cretaceous. Claims of an upper Albian age based on ammonite biostratigraphy [44] have not been verified [45]. However, a U-Pb age of 98.79 Ma ± 0.62 Myr was obtained from zircons in volcanoclastics deposited contemporaenously with the amber [46]. Shi et al. (2012) and Yu et al. (2019) interpret this as a minimum age for the amber since it is bioeroded. Nevertheless, in the absence of other evidence, this U-Pb zircon age can be used to established a minimum constraint of 98.17 Ma [44].

#### C_15: Marchantiophyta: Radulaceae – [Frullaniaceae – Lejeuneaceae]

***Fossil taxon and specimen:*** *Radula cretacea* [PB22484, Nanjing Institute of Geology and Palaeontology, Chinese Academy of Sciences, Nanjing], from amber localities near the village of Tanai, Kachin State, Myanmar [47].

***Phylogenetic justification:*** The specimen was assigned to the genus *Radula* (Radulaceae, Porellales) based acute to acuminate leaf apices, gemmae exclusively produced from leaf- lobe marginal cells and female bracts present in two pairs [47].

***Minimum age:*** 98.17

***Maximum age:*** 515.5 Ma

***Age justification:*** The age of Burmese amber has long been contested, with estimates ranging between the Miocene and Cretaceous. Claims of an upper Albian age based on ammonite biostratigraphy [44] have not been verified [45]. However, a U-Pb age of 98.79 Ma ± 0.62 Myr was obtained from zircons in volcanoclastics deposited contemporaenously with the amber [46]. Shi et al. (2012) and Yu et al. (2019) interpret this as a minimum age for the amber since it is bioeroded. Nevertheless, in the absence of other evidence, this U-Pb zircon age can be used to established a minimum constraint of 98.17 Ma [44].

#### C_16: Marchantiophyta: Frullaniaceae – Lejeuneaceae

***Fossil taxon and specimen:*** *Frullania cretacea* [AMNH B-011, American Museum of Natural History, New York], from Myanmar [48]

***Phylogenetic justification:*** *F. cretacea* was initially placed in the extant genus *Frullania* based on the reddish leaf colour, leaf cells with a single mammilla, lateral branching and helmet-shaped water sacs [49]. More recent findings support this placement, with the identification of a stylus shape similar to extant *Frullania,* bilobed underleaves and more elongate and toothed subgynoecial underleaves [48].

***Minimum age:*** 98.17

***Maximum age:*** 515.5 Ma

***Age justification:*** The age of Burmese amber has long been contested, with estimates ranging between the Miocene and Cretaceous. Claims of an upper Albian age based on ammonite biostratigraphy [44] have not been verified [45]. However, a U-Pb age of 98.79 Ma ± 0.62 Myr was obtained from zircons in volcanoclastics deposited contemporaenously with the amber [46]. Shi et al. (2012) and Yu et al. (2019) interpret this as a minimum age for the amber since it is bioeroded. Nevertheless, in the absence of other evidence, this U-Pb zircon age can be used to established a minimum constraint of 98.17 Ma [44].

#### C_17: Bryophyta: Sphagnopsida – [Polytrichopsida – Bryopsida]

***Fossil taxon and specimen:*** Sphagnales

***Phylogenetic justification:*** Following Morris *et al.* [5]

***Minimum age:*** 330.7 Ma

***Maximum age:*** 515.5 Ma

***Age justification***: Following Morris *et al.* [5]

#### C_18: Bryophyta: Polytrichopsida – Bryopsida

***Fossil taxon and specimen:*** *Palaeocampylopus buragoase*

***Phylogenetic justification:*** Following Morris *et al.* [5] ***Minimum age:*** 271.8 Ma

***Maximum age:*** 515.5 Ma

***Age justification***: Following Morris *et al.* [5]

#### C_19: Bryophyta: Polytrichaceae – Tetraphidaceae

***Fossil Taxon and Specimen:*** *Meantoinea alophosioides* [Gemmiferous gametophyte shoot in rock slab UAPC-ALTA P15393 B (slides B bot series a), University of Alberta Paleobotanical Collections (UAPC-ALTA), Edmonton, Alberta, Canada] from the Lower Cretaceous deposits at Apple Bay (Vancouver Island, British Columbia, Canada) [50].

***Phylogenetic Justification:*** Bippus *et al.* [50] placed *M. alophosioides* within the extant family Polytrichaceae based on the presence of leaves with diagnostic anatomy, differentiated into sheathing base and free lamina, bearing photosynthetic lamellae, and a conducting strand present in the stem. A phylogenetic analysis of 91 morphological characters confirmed the position of *M. alophosioides* as a stem group member of Polytrichaceae [51].

***Minimum age:*** 132 Ma

***Maximum age:*** 515.5 Ma

***Age justification:*** Sediments of the Apple Bay locality are regarded as Longarm Formation equivalent and originally proposed as Barremian based on the presence of monosulcate reticulate pollen grains [52]. Klymiuck and Stockey [53] argued that these grains are more likely associated with extinct gymnosperms, and instead propose a Valanginian age, supported by oxygen isotope analysis (D. Gröcke, personal communication, 2005, 2013).

Thus, we propose the upper boundary of the Valanginian 132.6 ± 0.6 Ma and thus, 132 Ma as a minimum age constraint.

#### C_20: Bryophyta: Atrichum – Polytrichum

***Fossil Taxon and Specimen:*** *Eopolytrichum antiquum* [PP44714, Department of Geology, Field Museum, Chicago] from Buffalo Creek member of the Gaillard Formation (Upper Cretaceous), Crawford County, Georgia, USA [54].

***Phylogenetic Justification:*** Konopka et al. [54] determined that *Eopolytrichum* was a member of the extant family Polytrichaceae based on several features including the epiphragm, broadened columella apex, peristomial membrane structure, and echinulate spore sculpture, as well as the overall lamellae and leaf anatomy associated with Polytrichaceae.

Based on a cladistic analysis of 99 characters Bippus et al. [51] recovered *Eopolytrichum* nested within the extant genus *Polytrichum*.

***Minimum age***: 82.9 Ma

***Maximum age:*** 515.5 Ma

***Age justification:*** *Eopolytrichum* was recovered from the sediments of the Allon quarry, assigned to the Buffalo Creek Member of the Gaillard Formation. Based on palynological analyses, a Campanian age was initially inferred [55]. However, this was later revised to a late Santonian age based on the correlation of the Allon locality palynological assemblage and that of the Coastal Plain Province [56], which was assigned to Burnett’s calcareous nannofossil zone CC17 [57], the base of which correlates with the Santonian/Campanian boundary. Thus, a minimum age constraint of 83.6 ± 0.7 is proposed.

#### C_21: Bryophyta: Funariidae – [Dicraniidae + Bryidae]

***Fossil taxon and specimen:*** *Capimirinus riopretensis*

***Phylogenetic justification:*** Following Morris *et al.* [5]

***Minimum age:*** 268.3 Ma

***Maximum age:*** 515.5 Ma

***Age justification***: Following Morris *et al.* [5]

#### C_22: Bryophyta: Dicraniidae – Bryiidae

***Fossil Taxon and Specimen:*** *Tricarinella crassiphylla* [Gametophyte shoot in rock slab UAPC-ALTA P13311, University of Alberta Paleobotanical Collections, Edmonton, Alberta, Canada], from the Apple Bay locality, Longarm Formation equivalent, Quatsino Sound, northern Vancouver Island, British Columbia, Canada (latitude 50°36′21″N, 127°39′25″W; UTM 9U WG 951068) [58].

***Phylogenetic justification:*** *Tricarinella crassiphylla* is distinguished predominantly based on 3 characters: tristichous phyllotaxis, bistratose leaf lamina and a homogeneous costa. This set of characters, along with the clear implication that *T. crassiphylla* was an acrocarpous species, leads to its placement within Grimmiales [58]. Grimmiales + Dicranales form part of the Dicraniidae, which is in turn sister to Bryiidae, and so was used to establish a minimum age constraint on the divergence of *Ceratodon* (Dicranales) and Bryiidae.

***Minimum age:*** 133.3 Ma

***Maximum age:*** 515.5 Ma

***Age justification:*** Sediments of the Apple Bay locality are regarded as Longarm Formation equivalent and orginially proposed as Barremian based on the presence of monosulcate reticulate pollen grains [52]. Klymiuck and Stockey [53] argued that these grains are more likely associated with extinct gymnosperms, and instead propose a Valanginian age, supported by oxygen isotope analysis (D. Gröcke, personal communication, 2005, 2013). Thus, we propose the upper boundary of the Valanginian 133.9 ± 0.6 Ma and thus, 133.3 Ma as a minimum age constraint.

#### C_23: Bryophyta: Hypnanae - Brynanae

***Fossil taxon and specimen:*** *Krassiloviella limbelloides* [Gametophyte shoot in rock slabs UAPC-ALTA P17596 C bot and D, University of Alberta Paleobotanical Collections (UAPC-ALTA), Edmonton, Alberta, Canada] from the northern shore of Apple Bay, Quatsino Sound, on the west side of Vancouver Island, British Columbia, Canada (lat. 507360 2100N, long. 1277390 2500W; UTM 9U WG 951068) [59].

***Phylogenetic justification:*** *K. limbelloides* was described as a pleurocarpous moss based on the presence of pinnate branching, homocostate and pluricostate leaves [59]. This was further supported by features including a monopodial main stem, second-order branching, the lack of conducting strands in the stems, and thin laminal cell walls. This combination of features indicates membership within the superorder Hypnanae, and the extinct family Tricostaceae.

***Minimum age:*** 132 Ma

***Maximum age:*** 515.5 Ma

***Age justification:*** Sediments of the Apple Bay locality are regarded as Longarm Formation equivalent and orginially proposed as Barremian based on the presence of monosulcate reticulate pollen grains [52]. Klymiuck and Stockey [53] argued that these grains are more likely associated with extinct gymnosperms, and instead propose a Valanginian age, supported by oxygen isotope analysis (D. Gröcke, personal communication, 2005, 2013). Thus, we propose the upper boundary of the Valanginian 132.6 ± 0.6 Ma and thus, 132 Ma as a minimum age constraint.

#### C_24: Tracheophyta: Lycopodiophyta – Euphyllophyta

***Fossil taxon and specimen:*** *Zosterophyllum sp.* and *Chelinospora? sp.* from the Ordovician successions of central Sweden.

***Phylogenetic justification:*** The minimum age is derived following Morris *et al.* [5].

***Minimum age:*** 420.7 Ma

***Maximum age:*** 458.88 Ma

***Age justification***: The minimum age is derived following following Morris *et al.* [5]. The soft maximum is based on a probably trilete spore (*Chelinospora? sp*.) from the Ordovician successions of central Sweden [60]. Spores were sampled from the Borenshult-1 drillcore, comprising a well-dated succession of the Middle to Upper Ordovician. The successions are dated based on conodont biostratigraphy [61] and occur below the Kinnekulle K-bentonite, itself dated using U/Pb zircons to 454.41 ± 0.17 Ma [62, 63], confirming a Sandbian age. The spore was sampled from 48.2m, corresponding to the *Baltoniodus variabilis* biozone, with a maximum age corresponding to the top of the Sandbian, 458.18 ± 0.7 Ma [61, 64].

#### C_25: Lycopodiophyta: Isoetales – Lycopodiales

***Fossil taxon and specimen:*** *Leclercqia complexa*

***Phylogenetic justification:*** Following Morris *et al.* [5]

***Minimum age:*** 392.1 Ma

***Maximum age:*** 451 Ma

***Age justification***: Following Morris *et al.* [5]

#### C_26: Lycopodiophyta: Isoetales – Selaginellaes

***Fossil taxon and specimen:*** *Yuguangia ordinata* [Specimen BUP.H-y07, Department of Geology of Peking University, PRC], from the Haikou Formation of Yuguang village, Zhanyi District, Yunnan Province, southwestern China [65].

***Phylogenetic justification:*** *Yuguangia* is placed within Lycophytina based on its overall morphology, including external and internal anatomy and reproductive structures. A cladistic analysis by Hao *et al.* [65] based on 19 morphological characters placed *Yuguangia* as more closely related to the extant order Isoetales than Selaginellales, and thus it allows a minimum age constraint on the divergence of *Isoetes* and *Selaginella*.

***Minimum age:*** 377.7 Ma

***Maximum age:*** 451 Ma

***Age justification:*** A Givetian age for the Haikou Formation was determined based on the presence of a miospore assemblage of *Cirratriradites monogrammos, Archaeozonotriletes variabilis*, *Cymbosporites magnificus* rm \**Geminospora lemurata* [66]. Thus, we propose a minimum age constraint coinciding with the upper boundary of the Givetian (388.9 ± 1.2 Ma)

#### C_27: Lycopodiophyta: *Selaginella selaginoides* – [rhizophoric *Selaginella*]

***Fossil Taxon and Specimen:*** *Selaginella resimus* [BMNH V.62303-V.62330, Palaeontology Department, Natural History Museum, London] from the Puddlebrook assemblage (Lower Carboniferous), Drybrook Sandstone of the Forest of Dean, Gloucestershire [67].

***Phylogenetic Justification:*** Rowe [67] described small lycophyte leafy shoots with consistently narrow stems and terminal strobili, placing it within *Selaginella.* The leaves are isophyllous and decussately arranged, supporting a placement within the subgenus *Boreoselaginella.* Thus, *S. resimus* provides a minimum age for the divergence of *S. selaginoides* and the rhizophoric clade, following Westrand and Korall [68, 69].

***Minimum age:*** 322.8 Ma

***Maximum age:*** 451 Ma

***Age Justification:*** The plant-bearing outcrop of the Puddlebrook assembly is part of the Drybrook Sandstone at Plump Hill. The correlation between Puddlebrook and Plump Hill is supported by macrofossil remains of the lycophyte *Lepidostrobophyllum fimbriatum* and the pteridosperm *Diplopteridium*, both commonly occurring at Puddlebrook [70, 71]. Plump Hill is characterised by the miospores *Perotriletes tesselatus, Schultzospora* and *Carbaneuletes circularis,* indicative of an Asbian/Holkerian age. An Upper Visean age for the Puddlebrook assembly is further supported by the presence *Tetrapterites visensis,* which is known from the Upper Visean of Carboniferous limestome of the Menai region of Wales, and the presence of *Cribosporites cribellatus* [72],which supports and Asbian level. Following Poty *et al.* [73], the Asbian correlates with the Chesterian and the Warnantian, which both share an upper boundary with the Serpukhovian (323.2 Ma ± 0.4) and thus a minimum age of 322.8 Ma is proposed.

#### C_28: Lycopodiophyta: Selaginella subg. Stachygynandrum – Selaginella kraussiana

***Fossil taxon and specimen:*** *Selaginella sp.* [GZG.BST.21966-2002, Geoscience Centre, University of Göttingen] from the mid-Cretaceous Kachin Burmese amber.

***Phylogenetic justification:*** Based on the presence of bilateral strobili, Schmidt et al [74] were able to confidently assign six fossil species to the subgenus *Stachygynandrum*.

***Minimum age:*** 98.17 Ma

***Maximum age:*** 451 Ma

***Age justification:*** The age of Burmese amber has long been contested, with estimates ranging between the Miocene and Cretaceous. Claims of an upper Albian age based on ammonite biostratigraphy [44] have not been verified [45]. However, a U-Pb age of 98.79 Ma ± 0.62 Myr was obtained from zircons in volcanoclastics deposited contemporaenously with the amber [46]. Shi et al. (2012) and Yu et al. (2019) interpret this as a minimum age for the amber since it is bioeroded. Nevertheless, in the absence of other evidence, this U-Pb zircon age can be used to established a minimum constraint of 98.17 Ma [44].

#### C_29: Lycopodiophyta: Lycopodioideae – Lycopodielloideae

***Fossil Taxon and Specimen:*** *Retitriletes austroclavatidites* [GIX-SHA08-5-$2; R32/1, orschungsstelle für Paläobotanik at the Institut für Geologie und Paläontologie, Westfälische Wilhelms-Universität Münster, Germany], from the Section Peak Formation

***Phylogenetic Justification:*** *Retitriletes* is a spore genus which bears strong resemblance with the spores of extant *Lycopodium* on the basis of their isotetrahedral structure and distinctive reticulate ornamentation on the distal surface [75–77]. They are the first spores which can reliably be assigned to the Lycopodoioideae, and thus we follow Bauret *et al.* [76] in providing a minimum age for the divergence of the Lycopodioideae and Lycopodielloideae.

***Minimum age:*** 199.2 Ma

***Maximum age:*** 451 Ma

***Age justification:*** The earliest record of *Retitriletes* is recorded by Bomfleur *et al.* [78] in the upper part of the Section Peak at Priestly Glacier cliff. The assemblage is assigned to Price’s APJ1 assemblage based on the presence of intrastriate *Classopollis ssp.* and the absence of *Ichyosporites crateris-punctatus* [79], advocating and early-middle Sinemurian age. Radiometric ages of detrital zircons from c. 10m below the sample indicated ages of 214 ± 3 Ma and from c. 15m above the sample of 198 ± 2 Ma [80] which is congruent with a Sinemurian assignment. Thus, a minimum age constraint of 199.5 ± 0.3 Ma was proposed [81].

#### C_30: Euphyllophyta

***Fossil taxon and specimen:*** *Kenrickia bivena* [Axis in slab 557820-5, Smithsonian Institution, NMNH] from the Battery Point Formation (Early Devonian), Quebec, Canada

***Phylogenetic justification:*** A cladistic analysis by Toledo et al. [82] of 50 morphological characters supported *Kenrickia* as sister to all other radiatopsids, a lineage within euphyllophytes and outgroup to remaining progymnosperms and seed plants.

***Minimum age:*** 393.2 Ma

***Maximum age:*** 451 Ma

***Age justification***: Material was collected from the south shore of Gaspé Bay, near Douglastown, QC, Canada. The fossils form part of an allochthonous assemblage preserved in fluvial to coastal deposits [83, 84]. Palynological assemblages indicate the deposits near Douglastown belong to the late Emsian [85] and so our minimum age is based on the boundary of the Emsian, 394.3 ± 1.1 Ma [86].

#### C_31: Monilophyta: *Equisetum -* Pteridophyta

***Fossil taxon and specimen:*** *Ibyka amphikoma*

***Phylogenetic justification:*** Following Morris *et al.* [5]

***Minimum age:*** 384.7 Ma

***Maximum age:*** 451 Ma

***Age justification***: Following Morris *et al.* [5]

#### C_32: Monilophyta: Equisetum subg. Equisetum – Equisetum subg. Hippochaete

***Fossil taxon and specimen:*** *Equisetum fluviatoides* [US1-103] from 34 metres above the base of the Ravenscrag Formation at Ravenscrag Butte, 17 Km southwest of Eastend, Saskatchewan, Canada [87].

***Phylogenetic justification:*** *Equisetum fluviatoides* is identified as belonging to the crown group *Equisetum* based on vegetative and reproductive similarities to the extant species *E. fluviatile* [87]. A phylogenetic analysis by Elgorriaga et al. [88] placed *E. fluviatoides* as sister to *E. fluviatile* or *E. diffusum* within the subgenus *Equisetum*. The present study analysed the same phenotype and molecular data in a Bayesian framework and was not able to resolve the position of *E. fluviatoides* within the subgenus *Equisetum* but in a more basal position as sister to clade containing *E. fluviatile, E. diffusum* and *E. arvense*. Following our results, *E. fluviatoides* was used to calibrate the divergence of subgenus *Equisetum* from subgenus *Hippochaete*, here representing the crown group of *Equisetum*.

***Minimum age:*** 64.86 Ma

***Maximum age:*** 451.0 Ma

***Age justification*:** The minimum age constraint is based on the holotype of *Equisetum fluviatoides,* described from 34 metres above the base of the Ravenscrag Formation at Ravenscrag Butte, Saskatchewan, Canada by McIver and Basinger [87] who estimate an early Palaeocene age on the basis that the Ravenscrag Formation conformably overlies the Ferris/No.1 Coal Zone which at this site is established to approximate the Cretaceous- Palaeocene Boundary based on palynostratigraphy, about 3 metres below the top of magnetostratigraphic zone 29r [89]. Thus, the horizon from which the holotype of *Equisetum fluviatoides* was derived falls within the 29n magnetozone, the minimum age of which can be constrained by the 29n-28r boundary, which is dated to 64.86 Ma [22].

#### C_33: Monilophyta: Marratiopsida – Psilotopsida

***Fossil taxon and specimen:*** *Psaronius simplicaulis*

***Phylogenetic justification:*** Following Morris *et al.* [5] ***Minimum age:*** 318.71 Ma

***Maximum age:*** 451.0 Ma

***Age justification***: Following Morris *et al.* [5]

#### C_34: Monilophyta: *Ophioglossum – Sceptridium*

***Fossil taxon and specimen:*** *Botrychium wightonii* [S1333 in the Palaeobotanical Herbarium, Department of Botany, University of Alberta] from the Palaeocene Genesee locality, North Saskatchewan River, Canada.

***Phylogenetic justification:*** Based on the morphology of sporophores and trophophores, Rothwell & Stockey [90] concluded that *B. wightonii* could be assigned to the extant genus *Botrychium* with high confidence.

***Minimum age***: 65.5

***Maximum age:*** 451.0 Ma

***Age justification:*** Material was collected from the Genesee locality, along the bank of the North Saskatchewan River 45 miles southwest of Edmonton, Canada. The megafloral assemblage at the Genesee locality indicates a Tiffanian age [91]. More recently, Stockey *et al.* [92] indicated that the Genesee flora belonged to the upper Scollard Formation, pollen zone P1, which correlates with the *Wodehouseia fimbriata* biozone [93], indicating an earliest Palaeocene age. U-Pb dating of the Scollard Formation and the overlying Battle Formation indicates an lower bound of 65.5 Ma for the *Wodehouseia fimbriata* biozone [94].

#### C_35: Monilophyta: Gleicheniales – [Cyatheales – Polypodiales]

***Fossil taxon and specimen****: Oligocarpia gothanii* [S138027–S138989, S147989 and KC 1983–1, Department of Palaeobotany of the Swedish Natural History Museum, Stockholm] from the Mid Permian of the Shihhotse valley near Ch’en-chia-yu near Taiyuan City in Shanxi Province, China [95].

***Phylogenetic justification:*** *Oligocarpia* has been associated with gleicheniaceous ferns on the basis of both spore and leaf morphology [95, 96].

***Minimum age:*** 268.3 Ma

***Maximum age:*** 451.0 Ma

***Age justification:*** *Oligocarpia gothanii* was collected from the beds of the south side of the Shihhotse valley near Ch’en-chia-yu near Taiyuan City in Shanxi Province, China [95]. These beds belong to the Lower Shihhotse Formation, which has been correlated with the Kungarian-Roadian stages of the Mid Permian [97, 98]. Thus, our minimum age is based on the upper boundary of the Roadian stage, estimated at 268.8 ± 0.5 Ma.

#### C_36: Monilophyta: Cyatheaceae – Thyrsopteridaceae

***Fossil Taxon and Specimen:*** *Oguracaulis banksii* [Z2286, Tasmania Museum and Art Gallery, Hobart, Tasmania] from the Lune River Fossil Site (Early to Middle Jurassic), Ida Bay, Tasmania [99].

***Phylogenetic Justification:*** Phylogenetic analysis based on characters derived from fossilized stems placed *Oguracaulis banksii* as sister to *Thyrsopteris* [100].

***Minimum age:*** 178 Ma

***Maximum age:*** 451 Ma

***Age justification:*** The flora of the Lune River area, Tasmania was initially estimated as Tertiary based on the similarity of the assemblage to others across Tasmania [101]. However, a revision of the taxonomic affinities of the flora suggested a Late Jurassic to Early Cretaceous age [102]. A basalt outcrop upslope of the fossil localities was shown to be comagmatic with Jurassic dolerite [103] and volcanolithic sandstone containing the fossils was recently dated at 182 ± 4 Ma (U-Pb zircon) [104], indicating a Pliensbachian-Toarcian age. Thus, we propose a minimum age constraint of 178 Ma.

#### C_37: Monilophyta: Lindseaceae – Lonchitidaceae

***Fossil Taxon and Specimen:*** Lindsaceae gen. sp. indet. [GZG.BST.21952 in the Geoscientific Collections of the Georg August University Göttingen, Germany] from the Burmese Amber, Myanmar.

***Phylogenetic Justification:*** The fusion of multiple veins below elongate terminal sori allows unambiguous assignment to Lindsaceae [105].

***Minimum age:*** 98.17

***Maximum age*: *451***

***Age Justification:*** The age of Burmese amber has long been contested, with estimates ranging between the Miocene and Cretaceous. Claims of an upper Albian age based on ammonite biostratigraphy [44] have not been verified [45]. However, a U-Pb age of 98.79 Ma

± 0.62 Myr was obtained from zircons in volcanoclastics deposited contemporaenously with the amber [46]. Shi et al. (2012) and Yu et al. (2019) interpret this as a minimum age for the amber since it is bioeroded. Nevertheless, in the absence of other evidence, this U-Pb zircon age can be used to established a minimum constraint of 98.17 Ma [44].

#### C_38: Monilophyta: Pteridiinae + Polypodiinae

***Fossil Taxon and Specimen:*** *Pteris sp.* [Palaeontological Institute, Moscow, Catalogue no. 4799-10] from the mid-Cenomanian of the Nammoura locality, Mount-Liban, Lebanon [106].

***Phylogenetic Justification:*** The specimen was assigned to the extant genus *Pteris* based on strong similarities in pinnule shape and coenosori position [106]. Schneider *et al.* supported its position as a pteridoid fern, but thought that no further assignment was possible and it was used as a minimum age for the divergence of Pteridaceae and Polypodiinae [107].

***Minimum age:*** 100.3 Ma

***Maximum age:*** 451 Ma

***Age justification:*** The Nammoura lagerstätte is estimated as latest middle Cenomanian in age based on the foraminiferan biostratigraphy and the characteristic species *Rhapydionina laurinensis* [106]. The upper units of the bed are marked by the presence of orbitolinids, indicating a middle Cenomanian age, which are absent in the lower units. As the unit from which the specimen was collected is not apparent, we took the upper boundary of the Cenomanian as the minimum age constraint, estimated at 100.5 ± 0.1, thus 100.3 Ma

#### C_39: Monilophta: Eupolypods I + Eupolypods II

***Fossil Taxon and Specimen:*** *Woodwardia sp.* from the Jose Creek Member, McRae Formation, New Mexico, USA [108].

***Phylogenetic Justification:*** Leaf impressions demonstrating diagnostic traits including a toothed margin, sickle-shaped pinnules, veins that anastomose at the midvein to form costal areoles, veins outside the costal areoles that run to the margin, and the absence of freely ending veinlets indicate a membership to the extant genus *Woodwardia*, a member of the Eupolypods II clade (Aspleniineae) [108].

***Minimum age:*** 72 Ma

***Maximum age: 451***

***Age justification:*** Leaf impression were collected from the lower Jose Creek Member of the McRae Formation. U-Pb age estimates from multiple horizons within the McRae formation have confirmed a Late Campanian age, rather than the previously suggested Maastrichtian [109]. We use the upper boundary of the Campanian, 72.2 ± 0.2 Ma, thus 72 Ma, as a minimum age constraint.

**Discussion:** *Holttumopteris burmensis*, a putative crown member of Eupolypods II was recently describe from Early Cretaceous amber deposits in Myanmar [110]. The authors decided that it was likely a member of the derived family Thelypteridaceae, but based on the lack of preservation of key characters could not confidently assign it to an extant family.

#### C_40: Spermatophyta: Acrogymnospermae – Angiospermae

***Fossil taxon and specimen:*** *Cordaixylon iowensis*

***Phylogenetic justification:*** Following Morris *et al.* [5]

***Minimum age:*** 308.14 Ma

***Maximum age:*** 365.63 Ma

***Age justification***: Following Morris *et al.* [5]

#### C_41: Acrogymnospermae: [Ginkgoopsida + Cycadopsida] – Pinopsida

***Fossil taxon and specimen:*** *Cordaixylon iowensis*

***Phylogenetic justification:*** Following Morris *et al.* [5]

***Minimum age:*** 308.14 Ma

***Maximum age:*** 365.63 Ma

***Age justification***: Following Morris *et al.* [5]

#### C_42: Acrogymnospermae: Ginkgoopsida – Cycadopsida

***Fossil taxon and specimen:*** *Iratinia australis* [MCT1487-PB, Museu de Ciências da Terra, Rio de Janeiro, Brazil] from the Irati Formation of the Kungurian of the Paraná Basin, Brazil

***Phylogenetic justification:*** Though initially proposed as a lycopsid [111], based on multiple morpho-anatomical traits Spiekermann *et al.* [112] determined that *I. australis* belonged to the extant order Cycadales.

***Minimum age:*** 274 Ma

***Maximum age:*** 365.63 Ma

***Age justification***: Material was collected from the Irati Formation, the lowermost unit of the Passa Dois Group. The Assistência Member from the upper part of the Irati Formation was U-Pb zircon dated to 279.9 ± 4.8 to 280 ± 3 Ma [113], placing it within the Kungurian. The Taquaral Member at the base of the formation was also placed in the Kungurian based on palynological evidence [114]. Thus, a Kungurian age is determined for the Irati Formation and a minimum age of 274.4 ± 0.5 Ma.

#### C_43: Pinopsida: Taxaceae - Cupressaceae

***Fossil taxon and specimen:*** *Palaeotaxus rediviva* from the Rhaetian of southern Sweden.

***Phylogenetic justification:*** *Palaeotaxus* forms the first credible record of the family Taxaceae and was placed within Taxaceae in a phylogenetic analysis of morphological characters [115, 116].

***Minimum age:*** 201.2 Ma

***Maximum age:*** 321.48 Ma

***Age justification***: Material was collected from the upper coal bed of the Skromberga Colliery in Scania, Sweden. The Skromberga locality has been assigned to the Bjuv member [117] and Pott & McLoughlin and McElwain et al indicate a Rhaetian age based on the correlation of the *Thaumatopteris-Lepidopteris* flora [118–120]. Thus, a minimum age is established based on the upper boundary of the Rhaetian 201.4 ± 0.2 Ma.

#### C_44: Pinopsida: Cunninghamia – Juniperus

***Fossil taxon and specimen:*** *Chamaeocyparis corpulenta* from the Upper Cretaceous Comox Formation of Vancouver Island, British Columbia.

*Phylogenetic justification:* Based on morphological similarities of the seed cone structure and the cones on the axes, *C. corpulenta* is considered the oldest known representative of the Cupresseae [121, 122].

***Minimum age:*** 83.2

***Maximum age:*** 321.48 Ma

***Age justification:*** Material was recovered from the carbonaceous shales from the No.2 coal seam within the Cumberland Member of the Comox Formation, Vancouver Island [123].

Marine fauna above and below the Comox formation indicates a Santonian age and thus a minimum of 83.7 ± 0.5 Ma.

#### C_45: Pinopsida: Pinales – Gnetales

***Fossil taxon and specimen:*** *Eathiestrobus mackenziei*

[Specimen Pb483, Hunterian Museum, Glasgow, Scotland] from Eathie, Black Isle, Scotland. ***Phylogenetic justification:*** Based on a combination of diagnostic seed, cone and scale morphologies, *Eathiestrobus* was considered a member of Pinaceae [124]. However, the wide bract is a feature that distinguishes *Eathiestrobus* from all extant genera.

***Minimum age:*** 153.62 Ma

***Maximum age:*** 321.48 Ma

***Age justification:*** Coastal Jurassic exposures at Eathie are from the Kimmeridge Clay Formation [125]. Ammonite stratigraphy indicates belonging to the *Cymodoce* chronozone [126], yet dinoflagellate stratigraphy supports a placement in the *Baylei* zone (Lower Kimmerigian). The top of the *Cymodoce* zone correlates with the magnetostratigraphic zone M24, the top of which is dated to 153.62 Ma [127].

#### C_46: Pinopsida: *Pinus – Cedrus*

***Fossil taxon and specimen:*** *Pinus yorkshirensis* [BU4737 at the Lapworth Museum, University of Birmingham, United Kingdom] from the early Cretaceous Wealden of Yorkshire, United Kingdom.

***Phylogenetic justification:*** A cladistic analysis of morphological characters resolved *P. yorkshirensis* as a crown-group member of *Pinus* [128].

***Minimum age:*** 129 Ma

***Maximum age:*** 321.48 Ma

***Age justification:*** Material comes from the Speeton Clay Formation, north of Speeton, Yorkshire. Palynological samples collected from the same material include the dinoflagellate species *Callaiosphaeridium asymmetricum, Odontochitina operculata* and *?Cribroperidinium confossum* [128] which constrains the sediment to Late Hauterivian to Early Barremian [129]. Based on the range bases of these taxa, Ryberg *et al.* conclude that the sediment was formed during the Hauterivian-Barremian transition (131-129 Ma) and thus, we propose a minimum age of 129 Ma.

#### C_47: Pinopsida: *Pinus* subg. *Pinus – Pinus* subg. *Strobus*

***Fossil taxon and specimen:*** *Pinus triphylla* [E.C Jeffrey Collection, Harvard University, USA] from the Late Cretaceous Raritan Formation, Staten Island, USA.

***Phylogenetic justification:*** *P. triphylla* has been considered a stem lineage member of the subgenus *Pinus* based on the synapomorphy of possessing two vascular bundles in the needles [130, 131].

***Minimum age:*** 89.37 Ma

***Maximum age:*** 321.48 Ma

***Age justification:*** Material was collected from the Androvette Clay Pit, Staten Island, part of the Raritan Formation. Based on palynological evidence, a Turonian age is proposed for the formation [55, 132, 133]. Thus, we propose a minimum age based on the lower boundary of the Turonian of 89.75 ± 0.38.

#### C_48: Pinopsida: *Pinus* subsection *Australes*

***Fossil taxon and specimen:*** *Pinus storeyana* [University of California, Museum of Paleontology, holotype 7234] from the Coal Valley Formation, Celetom Quarry, Nevada, USA

***Phylogenetic justification:*** Based on seed cone morphology, Saladin *et al.* placed *P. storeyana* within the *Attenuatae* subclade [130, 134].

***Minimum age:*** 9 Ma

***Maximum age:*** 321.48 Ma

***Age justification:*** K-Ar dating of the Coal Valley Formation, Wassuk Group, Nevada suggests it was deposited between 13 and 9 Ma [135]. Thus, a minimum age of 9 Ma is proposed

#### C_49: Angiospermae

***Fossil taxon and specimen:*** Tricolpate pollen grain

***Phylogenetic justification:*** Following Morris *et al.* [5]

***Minimum age:*** 125 Ma

***Maximum age:*** 247Ma

***Age justification***: Following Morris *et al.* [5]

#### C_50: Angiospermae: Nympheales

***Fossil taxon and specimen:*** Tricolpate pollen grain

***Phylogenetic justification:*** Following Morris *et al.* [5]

***Minimum age:*** 125 Ma

***Maximum age:*** 247Ma

***Age justification***: Following Morris *et al.* [5]

#### C_51: Angiospermae: Austrobaileyales

***Fossil taxon and specimen:*** Tricolpate pollen grain

***Phylogenetic justification:*** Following Morris *et al.* [5]

***Minimum age:*** 125 Ma

***Maximum age:*** 247Ma

***Age justification***: Following Morris *et al.* [5]

#### C_52: Mesangiospermae

***Fossil taxon and specimen:*** Tricolpate pollen grain

***Phylogenetic justification:*** Following Morris *et al.* [5]

***Minimum age:*** 125 Ma

***Maximum age:*** 247Ma

***Age justification***: Following Morris *et al.* [5]

#### C_53: Magnoliids: [Magnoliales + Laurales] + Piperales

***Fossil taxon and specimen:*** *Endressinia brasiliana*

***Phylogenetic justification:*** Following Morris *et al.* [5]

***Minimum age:*** 110.87 Ma

***Maximum age:*** 247Ma

***Age justification***: Following Morris *et al.* [5]

**C_54:** Piperales: *Houttuynia – Saruma*

***Fossil taxon and specimen:*** *Saururus tuckerae*

***Phylogenetic justification:*** Following Morris *et al.* [5] ***Minimum age:*** 44.3 Ma

***Maximum age:*** 247Ma

***Age justification***: Following Morris *et al.* [5]

#### C_55: Eudicotyledonae

***Fossil taxon and specimen:*** *Hyrcantha decussata*

***Phylogenetic justification:*** Following Morris *et al.* [5]

***Minimum age:*** 119.6 Ma

***Maximum age:*** 128.63 Ma

***Age justification***: Following Morris *et al.* [5]

#### C_56: Eudicotyledonae: Vitales

***Fossil taxon and specimen:*** *Paleoclusia chevalieri*

***Phylogenetic justification:*** Following Barba-Montoya *et al.* [136]

***Minimum age:*** 85.84 Ma

***Maximum age:*** 128.63 Ma

***Age justification***: Following Barba-Montoya *et al.* [136]

#### C_57: Eudicotyledonae: Rosids minus Vitales

***Fossil taxon and specimen:*** *Paleoclusia chevalieri*

***Phylogenetic justification:*** Following Barba-Montoya *et al.* [136]

***Minimum age:*** 85.8 Ma

***Maximum age:*** 128.63 Ma

***Age justification***: Following Barba-Montoya *et al.* [136]

#### C_58: Eudicotyledonae: Ericales

***Fossil taxon and specimen:*** *Paleoenkianthus sayrevillensis*

***Phylogenetic justification:*** Following Barba-Montoya *et al.* [136]

***Minimum age:*** 85.8 Ma

***Maximum age:*** 128.63 Ma

***Age justification***: Following Barba-Montoya *et al.* [136]

#### C_59: Eudicotyledonae: Myrtales

***Fossil taxon and specimen:*** *Esqueiria futabensis*

***Phylogenetic justification:*** Following Barba-Montoya *et al.* [136]

***Minimum age:*** 83.3 Ma

***Maximum age:*** 128.63 Ma

***Age justification***: Following Barba-Montoya *et al.* [136]

#### C_60: Eudicotyledonae: Asteraceae

***Fossil taxon and specimen:*** *Tubulifloridites antipodica **Phylogenetic justification:*** Following Barba-Montoya *et al.* [136] ***Minimum age:*** 41.5 Ma

***Maximum age:*** 128.63 Ma

***Age justification***: Following Barba-Montoya *et al.* [136]

#### C_61: Eudicotyledonae: Salicaceae

***Fossil taxon and specimen:*** *Pseudosalix handleyi*

***Phylogenetic justification:*** Following Barba-Montoya *et al.* [136]

***Minimum age:*** 48.57 Ma

***Maximum age:*** 128.63 Ma

***Age justification***: Following Barba-Montoya *et al.* [136]

#### C_62: Eudicotyledonae: Solanales

***Fossil taxon and specimen:*** *Solanites crassus*

***Phylogenetic justification:*** Following Barba-Montoya *et al.* [136]

***Minimum age:*** 37.3 Ma

***Maximum age:*** 128.63 Ma

***Age justification***: Following Barba-Montoya *et al.* [136]

#### C_63: Monocotyledonae

***Fossil taxon and specimen:*** *Liliacidites sp*.

***Phylogenetic justification:*** Following Barba-Montoya *et al.* [136]

***Minimum age:*** 113 Ma

***Maximum age:*** 128.63 Ma

***Age justification***: Following Barba-Montoya *et al.* [136]

#### C_64: Monocotyledonae: Dioscoreales + Core Monocots

***Fossil taxon and specimen:*** *Cratolirion bognerianum* [MB.Pb. 1997/1233 (repository: Museum für Naturkunde, Berlin)] from the Crato Formation (Early Cretaceous), Santana do Cariri, Brazil [137].

***Phylogenetic justification:*** Phylogenetic analyses indicated a membership within the Pandanales + Dioscoreales lineage (Petrosaviidae) [137].

***Minimum age:*** 119.5 Ma

***Maximum age:*** 128.63 Ma

***Age justification:*** The age of the [5, 138]Crato Formation has been much debated but a 123 Ma ± 3.5 Myro been obtained for the overlying Ipubi Formation using Re-Os isotope dating [139], providing for a minimum constraint of 119.5 Ma [138].

#### C_65: Monocotyledonae: Riponogaceae (*Colcichum – Smilax*) ** NEW **

***Fossil taxon and specimen:*** *Ripogonum tasmanicum* from the Macquarie Harbour Formation (Early Eocene) from the central west coast of Tasmania [140]

***Phylogenetic justification:*** Based on a phylogenetic analysis of morphological characters, *Ripogonum tasmanicum* was placed within the extant genus *Ripogonum* (Ripogonaceae, Liliales). Phylogenetic analyses identify Riponogaceae and Smilaceae being sister lineages respective to Colchicaceae. ***Minimum age:*** 50.5 Ma

***Maximum age:*** 128.63 Ma

***Age justification:*** Fossil leaves were collected from mudstone deposits at the Lowana Road site, the Macquarie Harbour Formation, Tasmania. Spores, pollen and dinoflagellates from the site were attributed to the Planktonic Foraminiferal Zone P8-9 [141]corresponding to a

Mid-Ypresian age. The minimum age of the deposit is determined based on correlation of the Upper *Malvacipollis diversus* spore-pollen zone with zones P7 to P8 of Partridge [142] (52.5-50.5 Ma). Thus, a minimum age of 50.5 Ma is provided.

#### C_66: Monocotyledonae: Arecales – Poales

***Fossil taxon and specimen:*** *Sabalites carolinensis*

***Phylogenetic justification:*** Following Barba-Montoya *et al.* [136]

***Minimum age:*** 83.41 Ma

***Maximum age:*** 128.63 Ma

***Age justification***: Following Barba-Montoya *et al.* [136]

#### C_67: Poales: BOP-PACMAD

***Fossil taxon and specimen:*** *Changii indicum* [Slide 13160, Q-14-3, Birbal Sahni I. Palaeobotany, Lucknow, India] recovered from Coprolites from red clays in Lameta Formation, associated with the Deccan Volcanics (infra- and inter-trappan) at Pisdura (East and South sections), Central India. [143].

***Phylogenetic justification:*** Based on phylogenetic analyses, Prasad *et al.* [143] determined *C. indicum* as a member of Erhartoideae (Poaceae). More recent analyses have questioned this placement, and have suggested that it could belong to a number of tribes. As such, we follow Christin *et al.* [144] in a more conservative placement as a stem group member of the BOP clade.

***Minimum age:*** 66.261 Ma

***Maximum age:*** 128.63 Ma

***Age justification:*** Prasad *et al.* [61] interpreted the minimum age of *Changii indicum* as 65.0 Ma, while Iles *et al.* [145] adopted the radiometric and magnetostrategraphic dating of the Deccan Traps by Courtillot and Ren [146], supportted by age constraints of associated dinosaur coprolites to infer a minimum constraint of 66.0 Ma, based on the dating of the Maastrichtian-Palaeocene Boundary; this would revise to 65.98 Ma following Ogg & Hinnov [138]. However, the dating of the Deccan Traps has been revised [147] and, given the fossil occurrences predate the volcanic sequences [61] we can establish a minimum constraint based on the age of the oldest phase of volcanism, dated to 66.288 Ma ± 0.027 Myr, thus 66.261 Ma.

#### C_68: Poales: *Brachypodium – Oryza* ** NEW **

***Fossil taxon and specimen:*** *Stipa florissanti* [Syntypes: 34750, 34751 (USNM)] from Florissant and Florissant Fossil Beds National Monument, CO, USA [148]

***Phylogenetic justification:*** Based on morphology this fossil was assigned the extant grass lineage *Stipa.* Extant *Stipa* has a strictly Old World distribution [149], so the generic assignment has been questioned [150]. We follow Iles *et al.* [145] and their assignment to the stem group Stipeae, itself nested within Pooideae [151].

***Minimum age:*** 33.705 Ma

***Maximum age:*** 128.63 Ma

***Age justification:*** The age interpretation of the Florissant Formation is usually based on Evanoff et al. [152] who used ^40^Ar/^39^Ar dating of sanidine crystals in detrital pumice, yielding an age of 34.07 Ma ± 0.1 Myr. However, its detrital origin renders this a maximum constraint on the age of the fossil. Nevertheless, Florissant Formation falls fully within the C13r magnetozone [153] and the base of the succeeding C13n magnetozone is dated to 33.705 [22]

## Notes

### Competing Interest Statement

The authors have declared no competing interest.

## References

Albalat, R. and Cañestro, C. (2016) ‘Evolution by gene loss’, Nature Reviews Genetics, 17(7), pp. 379–391. doi: 10.1038/nrg.2016.39.

Bauer, H. et al. (2013) ‘The stomatal response to reduced relative humidity requires guard cell-autonomous ABA synthesis’, Current Biology. doi: 10.1016/j.cub.2012.11.022.

Bell, D. et al. (2020) ‘Organellomic data sets confirm a cryptic consensus on (unrooted) land-plant relationships and provide new insights into bryophyte molecular evolution’, American Journal of Botany, pp. 91–115. doi: 10.1002/ajb2.1397.

Bergsten, J. (2005) ‘A review of long-branch attraction’, Cladistics. John Wiley & Sons, Ltd, pp. 163–193. doi: 10.1111/j.1096-0031.2005.00059.x.

Berry, J. A., Beerling, D. J. and Franks, P. J. (2010) ‘Stomata: Key players in the earth system, past and present’, Current Opinion in Plant Biology. doi: 10.1016/j.pbi.2010.04.013.

Blanquart, S. and Lartillot, N. (2008) ‘A site- and time-heterogeneous model of amino acid replacement’, Molecular Biology and Evolution. doi: 10.1093/molbev/msn018.

Bowles, A. M. C., Bechtold, U. and Paps, J. (2020) ‘The Origin of Land Plants Is Rooted in Two Bursts of Genomic Novelty’, Current Biology, 30, pp. 530–536. doi: 10.1016/j.cub.2019.11.090.

Bowman, J. L. et al. (2017) ‘Insights into Land Plant Evolution Garnered from the Marchantia polymorpha Genome’, Cell, 171(2), pp. 287–304.e15. doi: 10.1016/j.cell.2017.09.030.

Buchfink, B., Xie, C. and Huson, D. H. (2014) ‘Fast and sensitive protein alignment using DIAMOND’, Nature Methods. Nature Publishing Group, pp. 59–60. doi: 10.1038/nmeth.3176.

Cannell, N. et al. (2020) ‘Multiple Metabolic Innovations and Losses Are Associated with Major Transitions in Land Plant Evolution’, Current Biology, 30(10), pp. 1783–1800.e11. doi: 10.1016/j.cub.2020.02.086.

Chanderbali, A. S. et al. (2017) ‘Evolution of floral diversity: genomics, genes and gamma.’, Philosophical transactions of the Royal Society of London. Series B, Biological sciences, 372(1713). doi: 10.1098/rstb.2015.0509.

Chen, J. and Wang, N. (2019) ‘Tissue cell differentiation and multicellular evolution via cytoskeletal stiffening in mechanically stressed microenvironments’, Acta Mechanica Sinica/Lixue Xuebao, 35(2), pp. 270–274. doi: 10.1007/s10409-018-0814-8.

Clark, J. W. and Donoghue, P. C. J. (2018) ‘Whole-Genome Duplication and Plant Macroevolution’, Trends in Plant Science. Elsevier Ltd, pp. 933–945. doi: 10.1016/j.tplants.2018.07.006.

Coleman, G. A. et al. (2021) ‘A rooted phylogeny resolves early bacterial evolution’, Science, 372(6542). doi: 10.1126/science.abe0511.

Cox, C. J. et al. (2014) ‘Conflicting phylogenies for early land plants are caused by composition biases among synonymous substitutions’, Systematic Biology, 63(2), pp. 272–279. doi: 10.1093/sysbio/syt109.

Criscuolo, A. and Gribaldo, S. (2010) ‘BMGE (Block Mapping and Gathering with Entropy): A new software for selection of phylogenetic informative regions from multiple sequence alignments’, BMC Evolutionary Biology. doi: 10.1186/1471-2148-10-210.

Donoghue, P. et al. (2021) ‘The evolutionary emergence of land plants’, Current Biology, 31(19), pp. R1281–R1298. doi: https://doi.org/10.1016/j.cub.2021.07.038.

Eddy, S. R. (2011) ‘Accelerated profile HMM searches’, PLoS Computational Biology, 7(10), p. 1002195. doi: 10.1371/journal.pcbi.1002195.

Edwards, D. et al. (2014) ‘Cryptospores and cryptophytes reveal hidden diversity in early land floras’, New Phytologist, 202(1), pp. 50–78. doi: 10.1111/NPH.12645.

Emms, D. M. and Kelly, S. (2017) ‘STRIDE: Species tree root inference from gene duplication events’, Molecular Biology and Evolution, 34(12), pp. 3267–3278. doi: 10.1093/molbev/msx259.

Emms, D. M. and Kelly, S. (2019) ‘OrthoFinder: Phylogenetic orthology inference for comparative genomics’, Genome Biology. doi: 10.1186/s13059-019-1832-y.

Enright, A. J., Van Dongen, S. and Ouzounis, C. A. (2002) ‘An efficient algorithm for large-scale detection of protein families’, Nucleic Acids Research. Oxford University Press, pp. 1575–1584. doi: 10.1093/nar/30.7.1575.

Federhen, S. (2012) ‘The NCBI Taxonomy database’, Nucleic Acids Research. doi: 10.1093/nar/gkr1178.

Feldberg, K. et al. (2021) Checklist of fossil liverworts suitable for calibrating phylogenetic reconstructions, Bryophyte Diversity and Evolution. doi: 10.11646/bde.43.1.6.

Flores, J. R. et al. (2021) ‘Defying death: incorporating fossils into the phylogeny of the complex thalloid liverworts (Marchantiidae, Marchantiophyta) confirms high order clades but reveals discrepancies in family-level relationships’, Cladistics, 37(3), pp. 231–247. doi: 10.1111/cla.12442.

Floyd, S. K. and Bowman, J. L. (2007) ‘The ancestral developmental tool kit of land plants’, International Journal of Plant Sciences, 168(1), pp. 1–35. doi: 10.1086/509079.

Gao, B. et al. (2020) ‘Phylogenomic synteny network analyses reveal ancestral transpositions of auxin response factor genes in plants’, Plant Methods (2020), 16(70). doi: 10.1186/s13007-020-00609-1.

Geyer, C. J. (1991) ‘Markov Chain Monte Carlo Maximum Likelihood’, Proceedings of the 23rd Symposium on the Interface.

Guijarro-Clarke, C., Holland, P. W. H. and Paps, J. (2020) ‘Widespread patterns of gene loss in the evolution of the animal kingdom’, Nature Ecology & Evolution, 4(4), pp. 519–523. doi: 10.1038/s41559-020-1129-2.

Harris, B. J. et al. (2020) ‘Phylogenomic Evidence for the Monophyly of Bryophytes and the Reductive Evolution of Stomata’, Current Biology. doi: 10.1016/j.cub.2020.03.048.

Harrison, C. J. and Morris, J. L. (2018) ‘The origin and early evolution of vascular plant shoots and leaves’, Philosophical Transactions of the Royal Society B: Biological Sciences. doi: 10.1098/rstb.2016.0496.

Hedges, S. B. et al. (2018) ‘Accurate timetrees require accurate calibrations’, Proceedings of the National Academy of Sciences of the United States of America, 115(41), pp. E9510– E9511. doi: 10.1073/pnas.1812558115.

Helsen, J. et al. (2020) ‘Gene Loss Predictably Drives Evolutionary Adaptation’, Molecular Biology and Evolution, 37(10), pp. 2989–3002. doi: 10.1093/MOLBEV/MSAA172.

Home — The Plant List (no date). Available at: http://www.theplantlist.org/ (Accessed: 21 June 2021).

Huang, C. H. et al. (2020) ‘Recurrent genome duplication events likely contributed to both the ancient and recent rise of ferns’, Journal of Integrative Plant Biology, 62(4), pp. 433–455. doi: 10.1111/jipb.12877.

Huerta-Cepas, J. et al. (2017) ‘Fast genome-wide functional annotation through orthology assignment by eggNOG-mapper’, Molecular Biology and Evolution, 34(8), pp. 2115–2122. doi: 10.1093/molbev/msx148.

Kanehisa, M., Sato, Y. and Morishima, K. (2016) ‘BlastKOALA and GhostKOALA: KEGG Tools for Functional Characterization of Genome and Metagenome Sequences’, Journal of Molecular Biology, 428(4), pp. 726–731. doi: 10.1016/j.jmb.2015.11.006.

Katoh, K. and Toh, H. (2008) ‘Recent developments in the MAFFT multiple sequence alignment program’, Briefings in Bioinformatics. doi: 10.1093/bib/bbn013.

Lartillot, N. et al. (2013) ‘Phylobayes mpi: Phylogenetic reconstruction with infinite mixtures of profiles in a parallel environment’, Systematic Biology. doi: 10.1093/sysbio/syt022.

Lartillot, N. and Philippe, H. (2004) ‘A Bayesian mixture model for across-site heterogeneities in the amino-acid replacement process’, Molecular Biology and Evolution, 21(6), pp. 1095–1109. doi: 10.1093/molbev/msh112.

Leebens-Mack, J. H. et al. (2019) ‘One thousand plant transcriptomes and the phylogenomics of green plants’, Nature. doi: 10.1038/s41586-019-1693-2.

Li, F. W. et al. (2014) ‘Horizontal transfer of an adaptive chimeric photoreceptor from bryophytes to ferns’, Proceedings of the National Academy of Sciences of the United States of America, 111(18), pp. 6672–6677. doi: 10.1073/pnas.1319929111.

Li, F. W. et al. (2015) ‘The origin and evolution of phototropins’, Frontiers in Plant Science. doi: 10.3389/fpls.2015.00637.

Li, F. W. et al. (2020) ‘Anthoceros genomes illuminate the origin of land plants and the unique biology of hornworts’, Nature Plants. doi: 10.1038/s41477-020-0618-2.

Matasci, N. et al. (2014) ‘Data access for the 1,000 Plants (1KP) project’, GigaScience. doi: 10.1186/2047-217X-3-17.

McDaniel, S. F. (2021) ‘Bryophytes are not early diverging land plants’, New Phytologist. Blackwell Publishing Ltd, pp. 1300–1304. doi: 10.1111/nph.17241.

Morris, J. L., Puttick, Mark N., et al. (2018) ‘Accurate timetrees do indeed require accurate calibrations’, Proceedings of the National Academy of Sciences of the United States of America. National Academy of Sciences, pp. E9512–E9513. doi: 10.1073/pnas.1812816115.

Morris, J. L., Puttick, Mark N, et al. (2018) ‘The timescale of early land plant evolution’, Proceedings of the National Academy of Sciences of the United States of America, 115(10), p. E2274. doi: 10.1073/PNAS.1719588115.

Nguyen, L. T. et al. (2015) ‘IQ-TREE: A fast and effective stochastic algorithm for estimating maximum-likelihood phylogenies’, Molecular Biology and Evolution. doi: 10.1093/molbev/msu300.

O’Malley, M. A., Wideman, J. G. and Ruiz-Trillo, I. (2016) ‘Losing Complexity: The Role of Simplification in Macroevolution.’, Trends in ecology & evolution, 31(8), pp. 608–621. doi: 10.1016/j.tree.2016.04.004.

Parham, J. F. et al. (2012) ‘Best Practices for Justifying Fossil Calibrations’, Systematic Biology, 61(2), pp. 346–359. doi: 10.1093/SYSBIO/SYR107.

Pires, N. D. and Dolan, L. (2012) ‘Morphological evolution in land plants: New designs with old genes’, Philosophical Transactions of the Royal Society B: Biological Sciences. doi: 10.1098/rstb.2011.0252.

Popper, Z. A. et al. (2011) ‘Evolution and Diversity of Plant Cell Walls: From Algae to Flowering Plants’, Annual Review of Plant Biology, 62(1), pp. 567–590.

Puttick, M. N. et al. (2018) ‘The Interrelationships of Land Plants and the Nature of the Ancestral Embryophyte’, Current Biology. doi: 10.1016/j.cub.2018.01.063.

Puttick, M. N., Clark, J. and Donoghue, P. C. J. (2015) ‘Size is not everything: Rates of genome size evolution, not C-value, correlate with speciation in angiosperms’, Proceedings of the Royal Society B: Biological Sciences, 282(1820). doi: 10.1098/rspb.2015.2289.

Radhakrishnan, G. V. et al. (2020) ‘An ancestral signalling pathway is conserved in intracellular symbioses-forming plant lineages’, Nature Plants, 6(3), pp. 280–289. doi: 10.1038/s41477-020-0613-7.

Rambaut, A. et al. (2018) ‘Posterior summarization in Bayesian phylogenetics using Tracer 1.7’, Systematic Biology, 67(5), pp. 901–904. doi: 10.1093/sysbio/syy032.

Raposo, G. and Stoorvogel, W. (2013) ‘Extracellular vesicles: Exosomes, microvesicles, and friends’, Journal of Cell Biology. The Rockefeller University Press, pp. 373–383. doi: 10.1083/jcb.201211138.

Raven, J. A. (2002) ‘Selection pressures on stomatal evolution’, New Phytologist. doi: 10.1046/j.0028-646X.2001.00334.x.

Dos Reis, M. et al. (2015) ‘Uncertainty in the Timing of Origin of Animals and the Limits of Precision in Molecular Timescales’, Current Biology, 25(22), pp. 2939–2950. doi: 10.1016/j.cub.2015.09.066.

Rensing, S. A. (2018) ‘Plant Evolution: Phylogenetic Relationships between the Earliest Land Plants’, Current Biology. Cell Press, pp. R210–R213. doi: 10.1016/j.cub.2018.01.034.

Rich, M. K. and Delaux, P. M. (2020) ‘Plant Evolution: When Arabidopsis Is More Ancestral Than Marchantia’, Current Biology, 30(11), pp. R642–R644. doi: 10.1016/j.cub.2020.04.077.

Sharma, V. et al. (2018) ‘A genomics approach reveals insights into the importance of gene losses for mammalian adaptations’, Nature Communications 2018 9:1, 9(1), pp. 1–9. doi: 10.1038/s41467-018-03667-1.

Shimodaira, H. (2002) ‘An approximately unbiased test of phylogenetic tree selection’, Systematic Biology. doi: 10.1080/10635150290069913.

Shimodaira, H. and Hasegawa, M. (2001) ‘CONSEL: for assessing the confidence of phylogenetic tree selection’, Bioinformatics. doi: 10.1093/bioinformatics/17.12.1246.

Simão, F. A. et al. (2015) ‘BUSCO: Assessing genome assembly and annotation completeness with single-copy orthologs’, Bioinformatics. doi: 10.1093/bioinformatics/btv351.

de Sousa, F. et al. (2019) ‘Nuclear protein phylogenies support the monophyly of the three bryophyte groups (Bryophyta Schimp.)’, New Phytologist. doi: 10.1111/nph.15587.

Stull, G. W. et al. (2021) ‘Gene duplications and phylogenomic conflict underlie major pulses of phenotypic evolution in gymnosperms’, Nature Plants, 7(8), pp. 1015–1025. doi: 10.1038/s41477-021-00964-4.

Su, D. et al. (2021) ‘Large-scale phylogenomic analyses reveal the monophyly of bryophytes and Neoproterozoic origin of land plants’, Molecular Biology and Evolution. doi: 10.1093/molbev/msab106.

Szöllosi, G. J. et al. (2013) ‘Efficient exploration of the space of reconciled gene trees’, Systematic Biology. doi: 10.1093/sysbio/syt054.

Szöllősi, G. J. et al. (2020) ‘Relative time constraints improve molecular dating Affiliations’, bioRxiv, p. 2020.10.17.343889. doi: 10.1101/2020.10.17.343889.

Szövényi, P., Gunadi, A. and Li, F.-W. W. (2021) Charting the genomic landscape of seed-free plants, Nature Plants. doi: 10.1038/s41477-021-00888-z.

Tomescu, A. M. F. et al. (2018) Why Are Bryophytes So Rare in the Fossil Record? A Spotlight on Taphonomy and Fossil Preservation, Transformative Paleobotan*y*.

Villarreal, J. C. and Renner, S. S. (2012) ‘Hornwort pyrenoids, carbon-concentrating structures, evolved and were lost at least five times during the last 100 million years’, Proceedings of the National Academy of Sciences of the United States of America, 109(46), pp. 18873–18878. doi: 10.1073/pnas.1213498109.

Villarreal, J. C. and Renner, S. S. (2014) ‘A review of molecular-clock calibrations and substitution rates in liverworts, mosses, and hornworts, and a timeframe for a taxonomically cleaned-up genus Nothoceros’, Molecular Phylogenetics and Evolution, 78(1), pp. 25–35. doi: 10.1016/j.ympev.2014.04.014.

de Vries, J. and Archibald, J. M. (2018) ‘Plant evolution: landmarks on the path to terrestrial life’, New Phytologist. doi: 10.1111/nph.14975.

Walden, N. et al. (2020) ‘Nested whole-genome duplications coincide with diversification and high morphological disparity in Brassicaceae’, Nature Communications, 11(1), p. 3795. doi: 10.1038/s41467-020-17605-7.

Wang, B. et al. (2010) ‘Presence of three mycorrhizal genes in the common ancestor of land plants suggests a key role of mycorrhizas in the colonization of land by plants’, New Phytologist, 186(2), pp. 514–525. doi: 10.1111/j.1469-8137.2009.03137.x.

Wang HC, Minh BQ, Susko E, R. A. (2014) ‘Modeling Site Heterogeneity with Posterior Mean Site Frequency Profiles Accelerates Accurate Phylogenomic Estimation’, Systems Biology, 67, pp. 216–235.

Warnock, R. C. M. et al. (2015) ‘Calibration uncertainty in molecular dating analyses: There is no substitute for the prior evaluation of time priors’, Proceedings of the Royal Society B: Biological Sciences, 282(1798). doi: 10.1098/rspb.2014.1013.

Warnock, R. C. M., Yang, Z. and Donoghue, P. C. J. (2012) ‘Exploring uncertainty in the calibration of the molecular clock’, Biology Letters, 8(1), pp. 156–159. doi: 10.1098/rsbl.2011.0710.

Wellman, C. H., Steemans, P. and Vecoli, M. (2013) ‘Chapter 29 Palaeophytogeography of Ordovician–Silurian land plants’, Geological Society, London, Memoirs, 38(1), p. 461.

Wellman, C. H. and Strother, P. K. (2015) ‘The terrestrial biota prior to the origin of land plants (embryophytes): A review of the evidence’, Palaeontology, 58(4), pp. 601–627. doi: 10.1111/pala.12172.

Wilhelmsson, P. K. I. et al. (2017) ‘Comprehensive Genome-Wide Classification Reveals That Many Plant-Specific Transcription Factors Evolved in Streptophyte Algae’, Genome Biology and Evolution, 9(12), pp. 3384–3397. doi: 10.1093/gbe/evx258.

Yang, Z. (2007) ‘PAML 4: Phylogenetic analysis by maximum likelihood’, Molecular Biology and Evolution, 24(8), pp. 1586–1591. doi: 10.1093/molbev/msm088.

Zhang, C. et al. (2018) ‘ASTRAL-III: polynomial time species tree reconstruction from partially resolved gene trees’, BMC Bioinformatics 2018 19:6, 19(6), pp. 15–30. doi: 10.1186/S12859-018-2129-Y.

## References

1. Tang Q., Pang K., Yuan X., Xiao S. 2020 A one-billion-year-old multicellular chlorophyte. Nature Ecology & Evolution 4(4), 543–549. (doi:10.1038/s41559-020-1122-9).

2. Zhang S.-H., Zhao Y., Ye H., Hu G.-H. 2016 Early Neoproterozoic emplacement of the diabase sill swarms in the Liaodong Peninsula and pre-magmatic uplift of the southeastern North China Craton. Precambrian Research 272, 203–225.

3. Yang D.-B., Xu W.-L., Xu Y.-G., Wang Q.-H., Pei F.-P., Wang F. 2012 U–Pb ages and Hf isotope data from detrital zircons in the Neoproterozoic sandstones of northern Jiangsu and southern Liaoning Provinces, China: implications for the Late Precambrian evolution of the southeastern North China Craton. Precambrian Research 216, 162-176.

4. Zhao H., Zhang S., Ding J., Chang L., Ren Q., Li H., Yang T., Wu H. 2019 New geochronologic and paleomagnetic results from early Neoproterozoic mafic sills and late Mesoproterozoic to early Neoproterozoic successions in the eastern North China Craton, and implications for the reconstruction of Rodinia. GSA Bulletin 132(3-4), 739–766. (doi:10.1130/b35198.1).

5. Morris J.L., Puttick M.N., Clark J.W., Edwards D., Kenrick P., Pressel S., Wellman C.H., Yang Z., Schneider H., Donoghue P.C.J. 2018 The timescale of early land plant evolution. Proceedings of the National Academy of Sciences 115(10), E2274–E2283. (doi:10.1073/pnas.1719588115).

6. Loiseau O., Weigand A., Noben S., Rolland J., Silvestro D., Kessler M., Lehnert M., Salamin N. 2020 Slowly but surely: gradual diversification and phenotypic evolution in the hyper-diverse tree fern family Cyatheaceae. Annals of Botany 125(1), 93–103.

7. Samuelsson J., Dawes P.R., Vidal G. 1999 Organic-walled microfossils from the Proterozoic Thule Supergroup, Northwest Greenland. Precambrian Research 96(1), 1–23. (doi:https://doi.org/10.1016/S0301-9268(98)00123-5).

8. Tomescu A., Bomfleur B., Bippus A., Savoretti A. 2018 Why Are Bryophytes So Rare in the Fossil Record? A Spotlight on Taphonomy and Fossil Preservation. (pp. 375–416.

9. Labandeira C.C., Tremblay S.L., Bartowski K.E., VanAller Hernick L. 2014 Middle Devonian liverwort herbivory and antiherbivore defence. New Phytologist 202(1), 247–258. (doi:10.1111/nph.12643).

10. he X., Sun Y., Zhu R.-L. 2013 The Oil Bodies of Liverworts: Unique and Important Organelles in Land Plants. Critical Reviews in Plant Sciences 32. (doi:10.1080/07352689.2013.765765).

11. Hernick L.V., Landing E., Bartowski K.E. 2008 Earth’s oldest liverworts— Metzgeriothallus sharonae sp. nov. from the Middle Devonian (Givetian) of eastern New York, USA. Review of Palaeobotany and Palynology 148(2), 154–162. (doi:https://doi.org/10.1016/j.revpalbo.2007.09.002).

12. Flores J.R., Bippus A.C., Suárez G.M., Hyvönen J. 2020 Defying death: incorporating fossils into the phylogeny of the complex thalloid liverworts (Marchantiidae, Marchantiophyta) confirms high order clades but reveals discrepancies in family-level relationships. Cladistics n/a(n/a). (doi:https://doi.org/10.1111/cla.12442).

13. Carpenter D.K. 2017 Charcoal, forests, and Earth’s palaeozoic geochemical oxygen cycle, University of Southampton.

14. Willis B., Bridge J. 1988 Evolution of Catskill river systems, New York state.

15. Richardson J.B. 1986 Silurian and Devonian spore zones of the Old Red Sandstone continent and adjacent regions. Geol Surv Canada, Bulletin 354, pl. 1-21.

16. Becker R., Gradstein F., Hammer O. 2012 The devonian period. In The geologic time scale (pp. 559–601, Elsevier.

17. Villarreal J.C., Renner S.S. 2012 Hornwort pyrenoids, carbon-concentrating structures, evolved and were lost at least five times during the last 100 million years. Proceedings of the National Academy of Sciences 109(46), 18873–18878. (doi:10.1073/pnas.1213498109).

18. Archangelsky S., Seoane L. 1996 Palynological studies of the baqueró formation (cretaceous), Santa Cruz Province, Argentina. VII. Ameghiniana 33, 307–313.

19. Passalia M.G., Llorens M., CÈsari S.N., Limarino C.O., Limarino C.O., Loinaze V.S.P., Vera E.I., Vera E.I. 2016 Revised stratigraphic framework of the Cretaceous in the Bajo Grande area (Argentinean Patagonia) inferred from new u Pb ages and palynology. Cretaceous Research 60, 152–166.

20. Chitaley S.D., Yawale N.R. 1980 On Notothylites nirulai gen. et sp. nov., a petrified Sporogonium from the Deccan Intertrappean beds of Mohgaonkalan, Madhya Pradesh. (

21. Thakre D., Samant B., Mohabey D., Sangode S.J., Srivastava P., Kapgate D., Mahajan R., Upreti N., Manchester S. 2017 A new insight into age and environments of intertrappean beds of Mohgaon Kalan, Chhindwara District, Madhya Pradesh using palynology, megaflora, magnetostratigraphy and clay mineralogy. Current science **VOL.** 112, 2193–2197.

22. Vandenberghe N., Hilgen F.J., Speijer R.P. 2012 The Paleogene Period. In The geologic timescale 2012 (eds. Gradstein F.M., Ogg J.G., Schmitz M., Ogg G.), pp. 855–921. Ansterdam, Elsevier.

23. Graham A. 1987 Miocene communities and paleoenvironments of southern Costa Rica. American Journal of Botany 74(10), 1501–1518.

24. Cassell D.T., Sen Gupta B.K. 1989 Foraminiferal stratigraphy and paleonvironments of the Tertiary Uscari Formation, Limon Basin, Costa Rica. Journal of Foraminiferal Research 19(1), 52–71. (doi:10.2113/gsjfr.19.1.52).

25. Hilgen F., Lourens L., Dam J.A. 2012 The Neogene Period. The Geologic Time Scale 2012 **2**, 923–978.

26. Seward A. 1904 The Jurassic Flora, 2: Liassic and Oolitic Floras of England. Catalogue of the Mesozoic Plants in the Department of Geology. Br Mus Nat Hist London 4(16), 192.

27. Li R., Li X., Deng S., Sun B. 2020 Morphology and microstructure of Pellites hamiensis nov. sp., a Middle Jurassic liverwort from northwestern China and its evolutionary significance. Geobios. (doi:https://doi.org/10.1016/j.geobios.2020.07.003).

28. Hongfu L. Early-Middle Jurassic Palynological Assemblages of Tmlufan-Hami Basin.

29. Deng S., Lu Y., Fan R., Pan Y., Cheng X., Fu G., Wang Q., Pan H., Shen Y., Wang Y. 2010 The Jurassic System of Northern Xinjiang, China. In Contributions to the 8th International Congress on the Jurassic System University of Science and Technology of China Press, Hefei (

30. Ogg J.G., Ogg G.M., Gradstein F.M. 2016 A concise geologic time scale: 2016, Elsevier.

31. Moisan P., Voigt S., Schneider J.W., Kerp H. 2012 New fossil bryophytes from the Triassic Madygen Lagerstätte (SW Kyrgyzstan). Review of Palaeobotany and Palynology 187, 29–37.

32. Schuster R.M. 1966 Hepaticae and Anthocerotae of North America east of the hundredth meridian.

33. Dobruskina I.A. 1995 Keuper (Triassic) Flora from Middle Asia (Madygen, Southern Fergana): Bulletin 5, New Mexico Museum of Natural History and Science.

34. Shcherbakov D. 2008 Madygen, Triassic Lagerstätte number one, before and after Sharov. Alavesia 2, 113–124.

35. Krassilov V.A. 1973 Mesozoic bryophytes from the Bureja Basin, Far East of the USSR. Palaeontographica Abteilung B, 95-105.

36. Krassilov V. 1970 Leafy hepatics from the Jurassic of the Bureja basin. Paleontol Zurn 3, 131–142.

37. Heinrichs J., Hentschel J., Wilson R., Feldberg K., Schneider H. 2007 Evolution of Leafy Liverworts (Jungermanniidae, Marchantiophyta): Estimating Divergence Times from Chloroplast DNA Sequences Using Penalized Likelihood with Integrated Fossil Evidence. Taxon 56(1), 31–44. (doi:10.2307/25065733).

38. Markevich V.S., Bugdaeva E.V. 2009 Palynological evidence for dating Jurassic-Cretaceous boundary sediments in the Bureya basin, Russian Far East. Russian Journal of Pacific Geology 3(3), 284–293. (doi:10.1134/S1819714009030087).

39. Markevich V., Bugdaeva E. 2014 Late jurassic-early cretaceous coal-forming plants of the Bureya Basin, Russian Far East. Stratigraphy and Geological Correlation 22(3), 239–255.

40. Schuster R.M., Janssens J.A. 1989 On Diettertia, an isolated Mesozoic member of the Jungermanniales. Review of palaeobotany and palynology 57(3-4), 277–287.

41. McGookey D., Haun J., Hale L., Goodell H., McCubbin D., Weimer R., Wulf G. 1972 Cretaceous system. Geologic Atlas of the Rocky Mountain Region: Rocky Mountain Association of Geologists, 190-228.

42. Decelles P.G. 1986 Sedimentation in a tectonically partitioned, nonmarine foreland basin: The Lower Cretaceous Kootenai Formation, southwestern Montana. Geological Society of America Bulletin 97(8), 911–931.

43. Heinrichs J., Schäfer-Verwimp A., Feldberg K., Schmidt A.R. 2014 The extant liverwort Gackstroemia (Lepidolaenaceae, Porellales) in Cretaceous amber from Myanmar. Review of Palaeobotany and Palynology 203, 48–52. (doi:https://doi.org/10.1016/j.revpalbo.2014.01.004).

44. Cruickshank R., Ko K. 2003 Geology of an amber locality in the Hukawng Valley, northern Myanmar. Journal of asian earth Sciences 21(5), 441–455.

45. Yu T., Kelly R., Mu L., Ross A., Kennedy J., Broly P., Xia F., Zhang H., Wang B., Dilcher D. 2019 An ammonite trapped in Burmese amber. Proceedings of the National Academy of Sciences 116(23), 11345–11350.

46. Shi G., Grimaldi D.A., Harlow G.E., Wang J., Wang J., Yang M., Lei W., Li Q., Li X. 2012 Age constraint on Burmese amber based on U–Pb dating of zircons. Cretaceous research 37, 155–163.

47. Bechteler J., Schmidt A.R., Renner M.A.M., Wang B., Pérez-Escobar O.A., Schäfer-Verwimp A., Feldberg K., Heinrichs J. 2017 A Burmese amber fossil of Radula (Porellales, Jungermanniopsida) provides insights into the Cretaceous evolution of epiphytic lineages of leafy liverworts. Foss Rec 20(2), 201–213. (doi:10.5194/fr-20-201-2017).

48. Heinrichs J., Reiner-Drehwald M.E., Feldberg K., von Konrat M., Hentschel J., Váňa J., Grimaldi D.A., Nascimbene P.C., Schmidt A.R. 2012 The leafy liverwort Frullania (Jungermanniopsida) in the Cretaceous amber forest of Myanmar. Review of Palaeobotany and Palynology 169, 21–28.

49. Hentschel J., Schmidt A.R., Heinrichs J. 2009 Frullania cretacea sp. nov.(Porellales, Jungermanniopsida), a leafy liverwort preserved in Cretaceous amber from Myanmar. Cryptogamie 30(3), 323.

50. Bippus A.C., Stockey R.A., Rothwell G.W., Tomescu A.M. 2017 Extending the fossil record of Polytrichaceae: Early Cretaceous Meantoinea alophosioides gen. et sp. nov., permineralized gametophytes with gemma cups from Vancouver Island. American journal of botany 104(4), 584–597.

51. Bippus A.C., Escapa I.E., Tomescu A.M.F. 2018 Wanted dead or alive (probably dead): Stem group Polytrichaceae. American Journal of Botany 105(8), 1243–1263. (doi:10.1002/ajb2.1096).

52. Sweet A.R. 2000 Applied research on two samples of Cretaceous age from Vancouver Island, British Columbia as requested by J. Haggart (GSC Pacific, Vancouver). In Geological Survey of Canada, Natural Resources Canada/Ressources Naturelles Canada, Ottawa, Ontario, Canada (

53. Klymiuk A.A., Stockey R.A. 2012 A Lower Cretaceous (Valanginian) seed cone provides the earliest fossil record for Picea (Pinaceae). American Journal of Botany 99(6), 1069–1082. (doi:10.3732/ajb.1100568).

54. Konopka A.S., Herendeen P.S., Merrill G.L.S., Crane P.R. 1997 Sporophytes and gametophytes of Polytrichaceae from the Campanian (Late Cretaceous) of Georgia, USA. International Journal of Plant Sciences 158(4), 489–499.

55. Christopher R.A. 1979 Normapolles and triporate pollen assemblages from the Raritan and Magothy Formations (Upper Cretaceous) of New Jersey. Palynology 3(1), 73–121. (doi:10.1080/01916122.1979.9989185).

56. Konopka A.S., Herendeen P.S., Crane P.R. 1998 Sporophytes and gametophytes of Dicranaceae from the Santonian (Late Cretaceous) of Georgia, USA. American Journal of Botany 85(5), 714–723. (doi:10.2307/2446542).

57. Burnett J.A. 1996 Nannofossils and Upper Cretaceous stage boundaries. Journal of Nannoplankton Research 18, 23–32.

58. Savoretti A., Bippus A.C., Stockey R.A., Rothwell G.W., Tomescu A.M.F. 2018 Grimmiaceae in the Early Cretaceous: Tricarinella crassiphylla gen. et sp. nov. and the value of anatomically preserved bryophytes. Annals of Botany 121(7), 1275–1286. (doi:10.1093/aob/mcy015).

59. Shelton G., Stockey R., Rothwell G.W., Tomescu A. 2016 Krassiloviella Limbelloides GEN. ET SP. NOV.: Additional diversity in the hypnanaean moss family tricostaceae (Valanginian, Vancouver Island, British Columbia). International Journal of Plant Sciences 177, 000–000. (doi:10.1086/688707).

60. Rubinstein C.V., Vajda V. 2019 Baltica cradle of early land plants? Oldest record of trilete spores and diverse cryptospore assemblages; evidence from Ordovician successions of Sweden. GFF 141(3), 181–190. (doi:10.1080/11035897.2019.1636860).

61. Bergström S.M., Lehnert O., Calner M., Joachimski M.M. 2012 A new upper Middle Ordovician–Lower Silurian drillcore standard succession from Borenshult in Östergötland, southern Sweden: 2. Significance of δ13C chemostratigraphy. Gff 134(1), 39-63.

62. Sell B., Ainsaar L., Leslie S. 2013 Precise timing of the Late Ordovician (Sandbian) super-eruptions and associated environmental, biological, and climatological events. Journal of the Geological Society 170(5), 711–714.

63. Bauert H., Isozaki Y., Holmer L.E., Aoki K., Sakata S., Hirata T. 2014 New U–Pb zircon ages of the Sandbian (Upper Ordovician)“Big K-bentonite” in Baltoscandia (Estonia and Sweden) by LA-ICPMS. GFF 136(1), 30–33.

64. Goldman D., Sadler P., Leslie S., Melchin M., Agterberg F., Gradstein F. 2020 The Ordovician Period. In Geologic Time Scale 2020 (pp. 631–694, Elsevier.

65. Hao S., Xue J., Wang Q., Liu Z. 2007 Yuguangia ordinata gen. et sp. nov., a New Lycopsid from the Middle Devonian (Late Givetian) of Yunnan, China, and Its Phylogenetic Implications. International Journal of Plant Sciences - INT J PLANT SCI 168, 1161–1175. (doi:10.1086/520727).

66. Marshall J.E., Zhu H., Wellman C.H., Berry C.M., Wang Y., Xu H., Breuer P. 2017 Provincial Devonian spores from South China, Saudi Arabia and Australia. Revue de Micropaléontologie 60(3), 403–409.

67. Rowe N. 1988 A herbaceous Lycophyte from the Lower Carboniferous Drybrook Sandstone of the Forest of Dean, Gloucestershire. Palaeontology 31, 69–83.

68. Klaus K., Schulz C., Bauer D., Stützel T. 2016 Historical biogeography of the ancient lycophyte genus Selaginella: Early adaptation to xeric habitats on Pangea. Cladistics. (doi:10.1111/cla.12184).

69. Weststrand S., Korall P. 2016 Phylogeny of Selaginellaceae: There is value in morphology after all! Am J Bot 103(12), 2136-2159. (doi:10.3732/ajb.1600156).

70. Lele K., Walton J. 1962 Fossil flora of the Drybrook Sandstone in the Forest of Dean, Gloucestershire, British Museum (Natural History).

71. Rowe N. 1988 New observations on the Lower Carboniferous pteridosperm Diplopteridium Walton and an associated synangiate organ. Botanical journal of the Linnean Society 97(2), 125–158.

72. Clayton G., Coquel R., Doubinger J., Gueinn K., Loboziak S., Owens B., Streel M. 1977 Carboniferous miospores of western Europe: illustration and zonation. Mededelingen- Rijks Geologische Dienst 29, 1–71.

73. Poty E., Aretz M., Hance L. 2014 Belgian substages as a basis for an international chronostratigraphic division of the Tournaisian and Viséan. Geological Magazine 151(2), 229–243.

74. Schmidt A.R., Regalado L., Weststrand S., Korall P., Sadowski E.-M., Schneider H., Jansen E., Bechteler J., Krings M., Müller P., et al. 2020 Selaginella was hyperdiverse already in the Cretaceous. New Phytologist 228(4), 1176–1182. (doi:https://doi.org/10.1111/nph.16600).

75. Field A.R., Testo W., Bostock P.D., Holtum J.A., Waycott M. 2016 Molecular phylogenetics and the morphology of the Lycopodiaceae subfamily Huperzioideae supports three genera: Huperzia, Phlegmariurus and Phylloglossum. Molecular Phylogenetics and Evolution 94, 635–657.

76. Bauret L., Field A.R., Gaudeul M., Selosse M.-A., Rouhan G. 2018 First insights on the biogeographical history of Phlegmariurus (Lycopodiaceae), with a focus on Madagascar. Molecular Phylogenetics and Evolution 127, 488–501. (doi:https://doi.org/10.1016/j.ympev.2018.05.004).

77. Juhász M. 1975 Lycopodiaceae spores from Lower Cretaceous deposits of Hungary. Acta Biological Szeged 21, 21–34.

78. Bomfleur B., Schöner R., Schneider J.W., Viereck L., Kerp H., McKellar J.L. 2014 From the Transantarctic Basin to the Ferrar Large Igneous Province—new palynostratigraphic age constraints for Triassic–Jurassic sedimentation and magmatism in East Antarctica. Review of Palaeobotany and Palynology 207, 18–37.

79. Price P. 1997 Permian to Jurassic palynostratigraphic nomenclature of the Bowen and Surat Basins. The Surat and Bowen basins, south-east Queensland, 137-178.

80. Elsner M., Schöner R., Gerdes A., Gaupp R. 2013 Reconstruction of the early Mesozoic plate margin of Gondwana by U–Pb ages of detrital zircons from northern Victoria Land, Antarctica. Geological Society, London, Special Publications 383(1), 211–232.

81. Hesselbo S., Ogg J., Ruhl M., Hinnov L., Huang C. 2020 The Jurassic Period. In Geologic time scale 2020 (pp. 955–1021, Elsevier.

82. Toledo S., Bippus A.C., Atkinson B.A., Bronson A.W., Tomescu A.M.F. 2021 Taxon sampling and alternative hypotheses of relationships in the euphyllophyte plexus that gave rise to seed plants: insights from an Early Devonian radiatopsid. New Phytologist 232(2), 914–927. (doi:https://doi.org/10.1111/nph.17511).

83. Cant D.J., Walker R.G. 1976 Development of a braided-fluvial facies model for the Devonian Battery Point Sandstone, Quebec. Canadian Journal of Earth Sciences 13(1), 102–119.

84. Griffing D.H., Bridge J.S., Hotton C.L. 2000 Coastal-fluvial palaeoenvironments and plant palaeoecology of the Lower Devonian (Emsian), Gaspé Bay, Québec, Canada. Geological Society, London, Special Publications 180(1), 61–84.

85. Mcgregor D., DC M. 1977 LOWER AND MIDDLE DEVONIAN SPORES OF EASTERN GASPE. II. BIOSTRATIGRAPHY.

86. Becker R., Marshall J., Da Silva A.-C., Agterberg F., Gradstein F., Ogg J. 2020 The devonian period. In Geologic Time Scale 2020 (pp. 733–810, Elsevier.

87. McIver E.E., Basinger J.F. 1989 The morphology and relationships of Equisetum fluviatoides sp.nov. from the Paleocene Ravenscrag Formation of Saskatchewan, Canada. Canadian Journal of Botany 67(10), 2937–2943. (doi:10.1139/b89-376).

88. Elgorriaga A., Escapa I.H., Rothwell G.W., Tomescu A.M.F., Rubén Cúneo N. 2018 Origin of Equisetum: Evolution of horsetails (Equisetales) within the major euphyllophyte clade Sphenopsida. American Journal of Botany 105(8), 1286–1303. (doi:10.1002/ajb2.1125).

89. Lerbekmo J. 1985 Magnetostratigraphic and biostratigraphic correlations of Maastrichtian to early Paleocene strata between south-central Alberta and southwestern Saskatchewan. Bulletin of Canadian Petroleum Geology 33(2), 213–226.

90. Rothwell G.W., Stockey R. 1989 Fossil Ophioglossales in the Paleocene of Western North America. American Journal of Botany - AMER J BOT 76. (doi:10.2307/2444411).

91. Chandrasekharam A. 1974 Megafossil flora from the Genesee locality, Alberta, Canada.

92. Stockey R.A., Hoffman G.L., Vavrek M.J. 2014 Paleobotany and paleoecology of the Munce’s Hill fossil locality near Red Deer, Alberta, Canada. Paleobotany and Biogeography: A Festschrift for Alan Graham in His 80th Year MSB 128, 367-388.

93. Lerbekmo J., Sweet A. 2007 Magnetobiostratigraphy of the continental Paleocene upper Coalspur and Paskapoo formations near Hinton, Alberta. Bulletin of Canadian Petroleum Geology 56(2), 118–146.

94. Eberth D.A., Kamo S.L. 2019 First high-precision U–Pb CA–ID–TIMS age for the Battle Formation (Upper Cretaceous), Red Deer River valley, Alberta, Canada: implications for ages, correlations, and dinosaur biostratigraphy of the Scollard, Frenchman, and Hell Creek formations. Canadian Journal of Earth Sciences 56(10), 1041–1051. (doi:10.1139/cjes-2018-0098).

95. Stevens L., Hilton J. 2009 Ontogeny and ecology of the filicalean fern Oligocarpia gothanii (Gleicheniaceae) from the Middle Permian of China. American journal of botany 96, 475–486. (doi:10.3732/ajb.0800221).

96. Wang Y., Guignard G., Barale G. 1999 Morphological and Ultrastructural Studies on in situ Spores of Oligocarpia (Gleicheniaceae) from the Lower Permian of Xinjiang, China. Int J Plant Sci 160(5), 1035–1045. (doi:10.1086/314178).

97. Glasspool I.J., Hilton J., Collinson M.E., Wang S.-J., Li Cheng S. 2004 Foliar physiognomy in Cathaysian gigantopterids and the potential to track Palaeozoic climates using an extinct plant group. Palaeogeography, Palaeoclimatology, Palaeoecology 205(1), 69–110. (doi:https://doi.org/10.1016/j.palaeo.2003.12.002).

98. Hilton J., Cleal C.J. 2007 The relationship between Euramerican and Cathaysian tropical floras in the Late Palaeozoic: Palaeobiogeographical and palaeogeographical implications. Earth-Science Reviews 85(3), 85–116. (doi:https://doi.org/10.1016/j.earscirev.2007.07.003).

99. Tidwell W.D., Nishida H., Webster N. 1989 Oguracaulis banksii gen. et sp. nov., a mid-Mesozoic tree-fern from Tasmania, Australia. In Papers and Proceedings of the Royal Society of Tasmania (pp. 15–25.

100. Lantz T.C., Rothwell G.W., Stockey R.A. 1999 Conantiopteris schuchmanii, gen. et sp. nov., and the Role of Fossils in Resolving the Phylogeny of Cyatheaceae sl. Journal of Plant Research 112(3), 361–381.

101. Gould R.E. 1972 Cibotium tasmanense sp. nov., A fossil tree-fern from the Tertiary of Tasmania. Australian Journal of Botany 20(1), 119–126.

102. Tidwell W.D. 1987 A new species of Osmundacaulis (O. jonesii sp. nov.) from Tasmania, Australia. Review of Palaeobotany and Palynology 52(2-3), 205–216.

103. Hergt J., Chappell B., McCulloch M., McDougall I., Chivas A. 1989 Geochemical and isotopic constraints on the origin of the Jurassic dolerites of Tasmania. Journal of Petrology 30(4), 841–883.

104. Bromfield K., Burrett C., Leslie R.A., Meffre S. 2007 Jurassic volcaniclastic-basaltic andesite-dolerite sequence in Tasmania: New age constraints for fossil plants from Lune River. Australian Journal of Earth Sciences 54, 965–974. (doi:10.1080/08120090701488297).

105. Regalado L., Schmidt A.R., Müller P., Kobbert M.J., Schneider H., Heinrichs J. 2017 The first fossil of Lindsaeaceae (Polypodiales) from the Cretaceous amber forest of Myanmar. Cretaceous Research 72, 8–12. (doi:https://doi.org/10.1016/j.cretres.2016.12.003).

106. Krassilov V., Bacchia F. 2000 Cenomanian florule of Nammoura, Lebanon. Cretaceous Research 21(6), 785–799. (doi:https://doi.org/10.1006/cres.2000.0229).

107. Schneider H., Schuettpelz E., Pryer K., Cranfill R., Magallon S., Lupia Ii R. 2004 Ferns diversified in the shade of angiosperms. Nature 428, 553–557. (doi:10.1038/nature02361).

108. Upchurch G., Mack G. 1998 Latest Cretaceous leaf megafloras from the Jose Creek Member, McRae Formation of New Mexico. New Mexico Geological Society Guidebook 49, 209–222.

109. Amato J.M., Mack G.H., Jonell T.N., Seager W.R., Upchurch G.R. 2017 Onset of the Laramide orogeny and associated magmatism in southern New Mexico based on U-Pb geochronology. GSA Bulletin 129(9-10), 1209–1226.

110. Regalado L., Schmidt A., Krings M., Bechteler J., Schneider H., Heinrichs J. 2017 Fossil evidence of eupolypod ferns in the mid-Cretaceous of Myanmar. Plant Systematics and Evolution. (doi:10.1007/s00606-017-1439-2).

111. Guerra-Sommer M. 1981 Plano-lenhoso vinculado a Sigillariaceae no Irati de S. Paulo. SIMPÓSIO REGIONAL DE GEOLOGIA 3, 176–179.

112. Spiekermann R., Jasper A., Siegloch A.M., Guerra-Sommer M., Uhl D. 2021 Not a lycopsid but a cycad-like plant: Iratinia australis gen. nov. et sp. nov. from the Irati Formation, Kungurian of the Paraná Basin, Brazil. Review of Palaeobotany and Palynology 289, 104415. (doi:https://doi.org/10.1016/j.revpalbo.2021.104415).

113. Rocha-Campos A., Basei M.A.S., Nutman A.P., Santos P., Passarelli C.R., Canile F., Rosa O., Fernandes M., Santa Ana H., Veroslavsky G. 2019 U-Pb zircon dating of ash fall deposits from the paleozoic Paraná basin of Brazil and Uruguay: a reevaluation of the stratigraphic correlations. The Journal of Geology 127(2), 167–182.

114. Rocha H., Mendes M., Pereira Z., Rodrigues C., Fernandes P., Lopes G., Sant’Anna L., Tassinari C., de Sousa M.L. 2020 New palynostratigraphic data of the Irati (Assistência Member) and the Corumbataí formations, Paraná Basin, Brazil, and correlation with other south American basins. Journal of South American Earth Sciences 102, 102631.

115. Florin R. 1948 On the Morphology and Relationships on the Taxaceae. Botanical Gazette 110(1), 31–39. (doi:10.1086/335515).

116. Dong C., Shi G., Herrera F., Wang Y., Herendeen P.S., Crane P.R. 2020 Middle–Late Jurassic fossils from northeastern China reveal morphological stasis in the catkin-yew. National Science Review 7(11), 1765–1767. (doi:10.1093/nsr/nwaa138).

117. Sivhed U. 1984 Litho-and biostratigraphy of the Upper Triassic-Middle Jurassic in Scania, southern Sweden, Sveriges geologiska undersökning.

118. Pott C., Mcloughlin S. 2011 The Rhaetian flora of Rögla, northern Scania, Sweden. Palaeontology 54(5), 1025–1051.

119. McElwain J., Beerling D., Woodward F. 1999 Fossil plants and global warming at the Triassic-Jurassic boundary. Science 285(5432), 1386–1390.

120. Harris T. 1931 Rhaetic floras. Biological reviews 6(2), 133–162.

121. McIver E. 1994 An early Chamaecyparis (Cupressaceae) from the Late Cretaceous of Vancouver Island, British Columbia, Canada. Canadian journal of botany 72(12), 1787–1796.

122. Stockey R.A., Kvacek J., Hill R.S., Rothwell G.W., Kvacek Z. 2005 The fossil record of Cupressaceae s. lat. A monograph of Cupressaceae and Sciadopitys 54, 68.

123. Bell W.A. 1957 Flora of the Upper Cretaceous Nanaimo Group of Vancouver Island, British Columbia. Geological Survey of Canada, Memoir (293), 1-84.

124. Rothwell G.W., Mapes G., Stockey R.A., Hilton J. 2012 The seed cone Eathiestrobus gen. nov.: fossil evidence for a Jurassic origin of Pinaceae. Am J Bot 99(4), 708–720. (doi:10.3732/ajb.1100595).

125. Riding J.B. 2005 Middle and Upper Jurassic (Callovian to Kimmeridgian) palynology of the onshore Moray Firth Basin, northeast Scotland. Palynology 29(1), 87–142.

126. Cope J.C.W. 1980 A Correlation of Jurassic rocks in the British Isles: part two: Middle and Upper Jurassic, Blackwell Scientific Publications.

127. Gradstein F.M., Ogg J.G., Schmitz M., Ogg G. 2012 The geologic time scale 2012, elsevier.

128. Ryberg P.E., Rothwell G.W., Stockey R.A., Hilton J., Mapes G., Riding J.B. 2012 Reconsidering Relationships among Stem and Crown Group Pinaceae: Oldest Record of the Genus Pinus from the Early Cretaceous of Yorkshire, United Kingdom. International Journal of Plant Sciences 173(8), 917–932. (doi:10.1086/667228).

129. Duxbury S. 1977 A palynostratigraphy of the Berriasian to Barremian of the Speeton Clay of Speeton, England.

130. Saladin B., Leslie A.B., Wüest R.O., Litsios G., Conti E., Salamin N., Zimmermann N.E. 2017 Fossils matter: improved estimates of divergence times in Pinus reveal older diversification. BMC evolutionary biology 17(1), 95.

131. Robison C.R. 1977 Pinus triphylla and Pinus quinquefolia from the Upper Cretaceous of Massachusetts. American Journal of Botany 64(6), 726–732.

132. Groot J.J., Penny J.S., Groot C.R. 1961 Plant microfossils and age of the Raritan, Tuscaloosa and Magothy formations of the eastern United States. Palaeontographica Abteilung B, 121-140.

133. Doyle J.A., Robbins E.I. 1977 Angiosperm pollen zonation of the continental Cretaceous of the Atlantic Coastal Plain and its application to deep wells in the Salisbury Embayment.

134. Axelrod D.I. 1986 Cenozoic History of Some Western American Pines. Annals of the Missouri Botanical Garden 73(3), 565–641. (doi:10.2307/2399194).

135. Evernden J.F., Savage D.E., Curtis G.H., James G.T. 1964 Potassium-argon dates and the Cenozoic mammalian chronology of North America. American Journal of Science 262(2), 145–198. (doi:10.2475/ajs.262.2.145).

136. Barba-Montoya J., dos Reis M., Schneider H., Donoghue P.C., Yang Z. 2018 Constraining uncertainty in the timescale of angiosperm evolution and the veracity of a Cretaceous Terrestrial Revolution. New Phytologist 218(2), 819-834.

137. Coiffard C., Kardjilov N., Manke I., Bernardes-de-Oliveira M.E.C. 2019 Fossil evidence of core monocots in the Early Cretaceous. Nature Plants 5(7), 691–696. (doi:10.1038/s41477-019-0468-y).

138. Ogg J., Hinnov L., Huang C. 2012 Cretaceous. In The geologic time scale (pp. 793–853, Elsevier.

139. Lúcio T., Souza Neto J.A., Selby D. 2020 Late Barremian / Early Aptian Re–Os age of the Ipubi Formation black shales: Stratigraphic and paleoenvironmental implications for Araripe Basin, northeastern Brazil. Journal of South American Earth Sciences 102, 102699. (doi:https://doi.org/10.1016/j.jsames.2020.102699).

140. Conran J.G., Carpenter R.J., Jordan G.J. 2009 Early eocene Ripogonum (Liliales: Ripogonaceae) leaf macrofossils from southern Australia. Australian Systematic Botany 22(3), 219–228.

141. Macphail M. 2005 Palynostratigraphic analysis of plant microfossils preserved in Early Eocene mudstones at Lowana Road, Strahan, west coast of Tasmania. Report© prepared for RSH & RJC) Consultant Palynological Services, Aranda, ACT.

142. Partridge A. 2006 Late Cretaceous–Cenozoic palynology zonations Gippsland Basin. In ‘Australian Mesozoic and Cenozoic palynology zonations’.(Coord. E Monteil) Geoscience Australia Record 2006/23. Chart 4.

143. Prasad V., Strömberg C., Leaché A., Samant B., Patnaik R., Tang L., Mohabey D., Ge S., Sahni A. 2011 Late Cretaceous origin of the rice tribe provides evidence for early diversification in Poaceae. Nature communications 2(1), 1–9.

144. Christin P.-A., Spriggs E., Osborne C.P., Strömberg C.A., Salamin N., Edwards E.J. 2014 Molecular dating, evolutionary rates, and the age of the grasses. Systematic biology 63(2), 153–165.

145. Iles W.J., Smith S.Y., Gandolfo M.A., Graham S.W. 2015 Monocot fossils suitable for molecular dating analyses. Botanical Journal of the Linnean Society 178(3), 346–374.

146. Courtillot V.E., Renne P.R. 2003 On the ages of flood basalt events. Comptes Rendus Geoscience 335(1), 113–140.

147. Schoene B., Eddy M.P., Samperton K.M., Keller C.B., Keller G., Adatte T., Khadri S.F.R. 2019 U-Pb constraints on pulsed eruption of the Deccan Traps across the end-Cretaceous mass extinction. Science 363(6429), 862–866. (doi:doi:10.1126/science.aau2422).

148. MacGinitie H.D. 1953 Fossil plants of the Florissant beds, Colorado, Carnegie institution of Washington.

149. Romaschenko K., Peterson P., Soreng R. 2010 Phylogenetics of Stipeae (Poaceae: Pooideae) based on plastid and nuclear DNA sequences/Ed. O. Seberg, G. Petersen, AS Barfod and JI Davis. Diversity, phylogeny and evolution in the Monocotyledons. (Aarhus, Denmark: Aarhus University Press.

150. Manchester S. 2001 Update on the megafossil flora of Florissant, Colorado. ser 4, 137–161.

151. II G.P.W.G. 2012 New grass phylogeny resolves deep evolutionary relationships and discovers C4 origins. New Phytologist 193(2), 304–312.

152. Evanoff E., Gregory-Wodzicki K.M., Johnson K. 2001 Fossil flora and stratigraphy of the Florissant Formation, Colorado.

153. Prothero D.R., Upton E. 2016 MAGNETIC STRATIGRAPHY OF THE EOCENE-OLIGOCENE TRANSITION FLORAS AND FAUNAS FROM THE WARNER MOUNTAINS, NORTHEASTERN CALIFORNIA. Fossil Record 4: Bulletin 67 67, 265.

