## Supplementary Figures for "Divergent evolutionary trajectories of bryophytes and tracheophytes from a complex common ancestor of land plants"

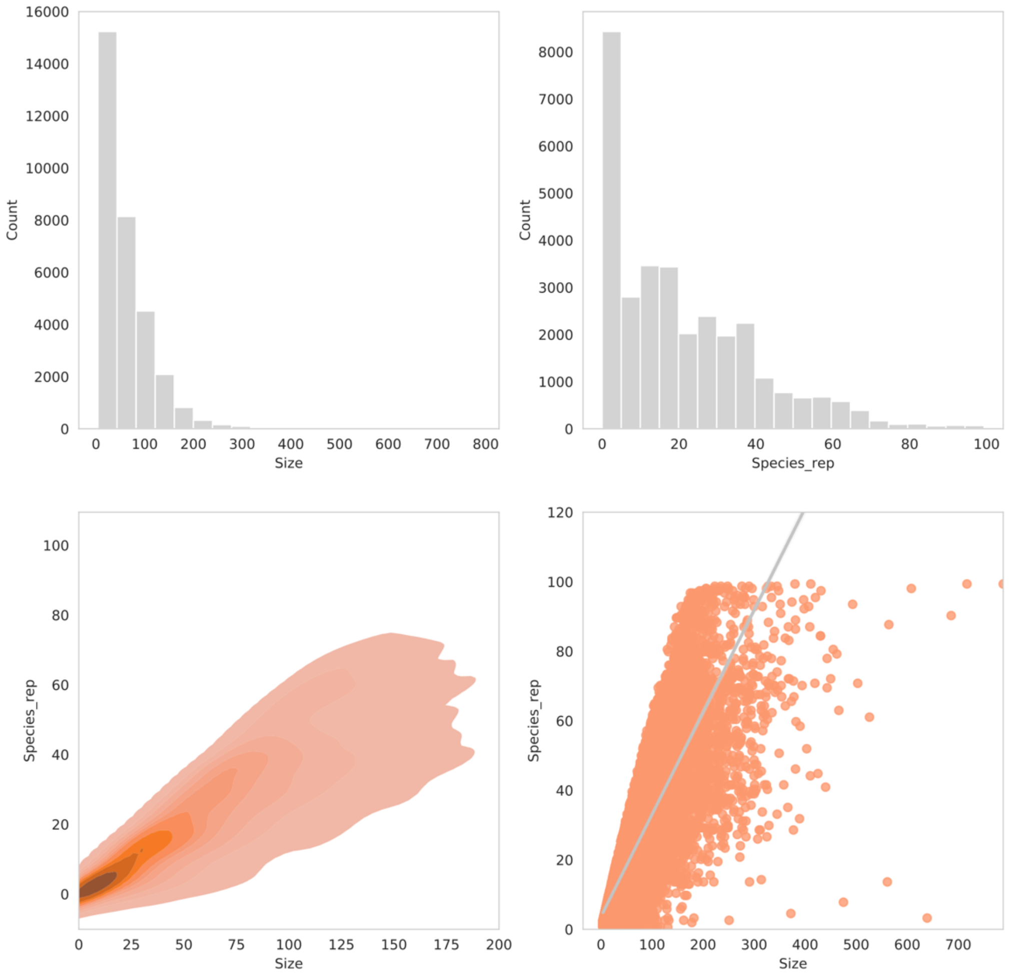


**Supplementary Figure 1. Gene family clusters for genome and transcriptome dataset – 154 species.** Top left, distribution of gene family size. Top right, distribution of gene family species representation. Bottom left,  density of gene family species repesentation and size. Bottom right, regression of gene family species representation and size.

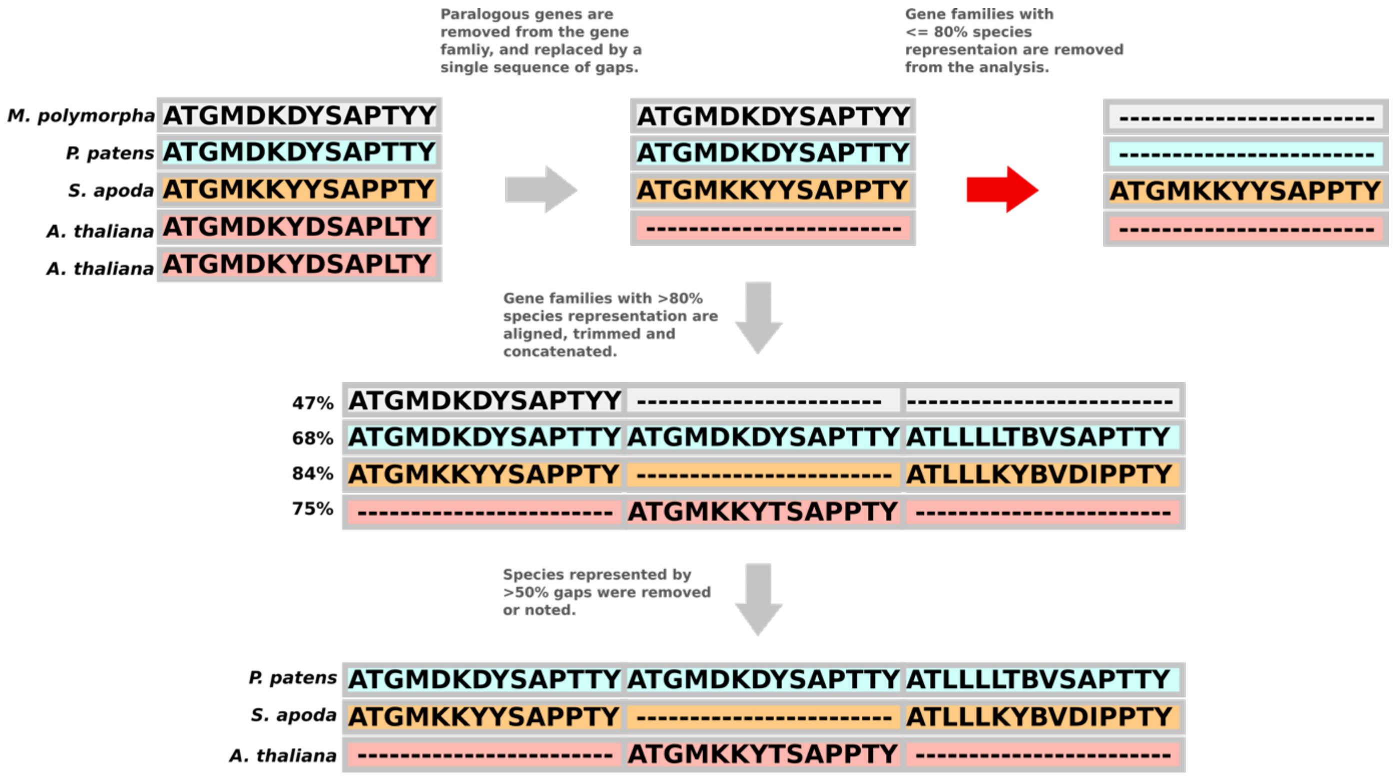


**Supplementary Figure 2. Graphical representation of orthology inference algorithm**. Species represented by more than one gene copy are removed from the gene family alignment and replaced by a single sequence of gaps. Gene families with more than 80% of the original species present are retained. Gene families are then aligned, trimmed and concatenated together to form a super matrix. Species with less than 50% gaps in the alignment are removed or noted if within 10% error – this step prevents single species that are continually removed from gene families to be in the final alignment.

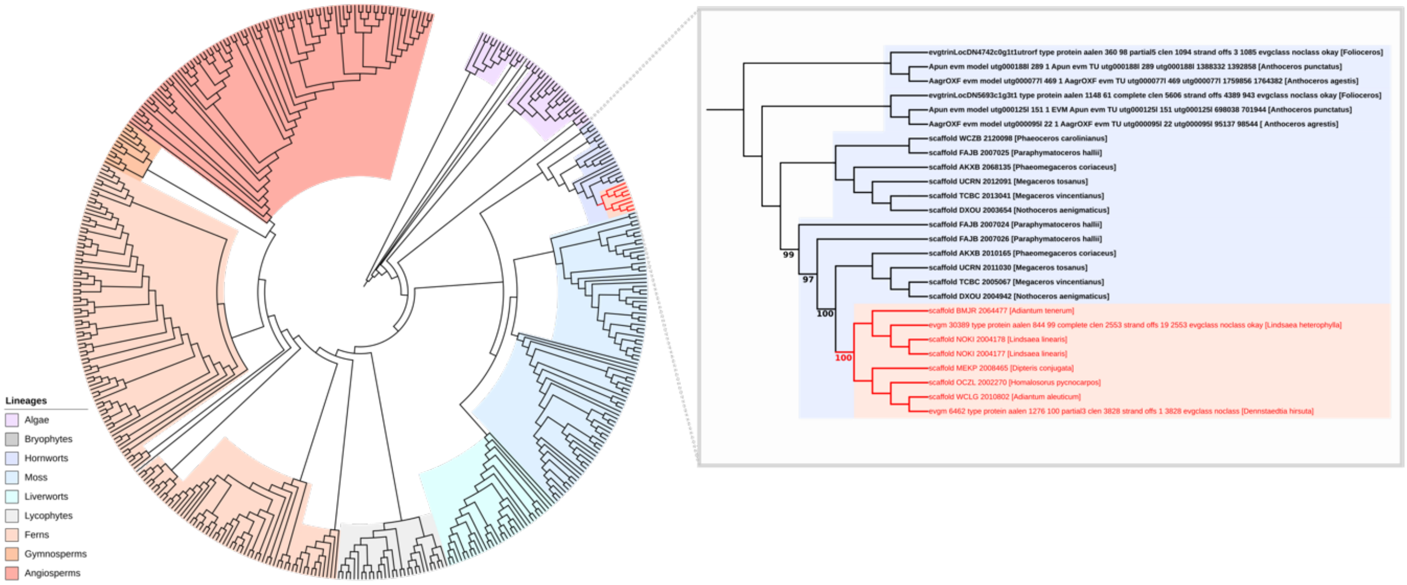


**Supplementary Figure 3. Phylogenetic tree highlighting the horizontal transfer of the chimeric neochrome photoreceptor (NEO).** The *Arabidopsis thaliana* protein sequence for PHOT1 was used to BLAST a database of 177 species of plant and transcriptomes. The homologous sequences were aligned with MAFFT and trimmed with BMGE. A maximum likelihood tree was inferred in IQ-TREE under the best fitting substitution model inferred with Bayesian Inference Criterion. 8 fern genes were resolved within the hornworts and were inferred to have undergone horizontal gene transfer (coloured red). This transfer was previously characterised (Li *et al.,* 2014), and we corroborate this finding with maximum bootstrap support.

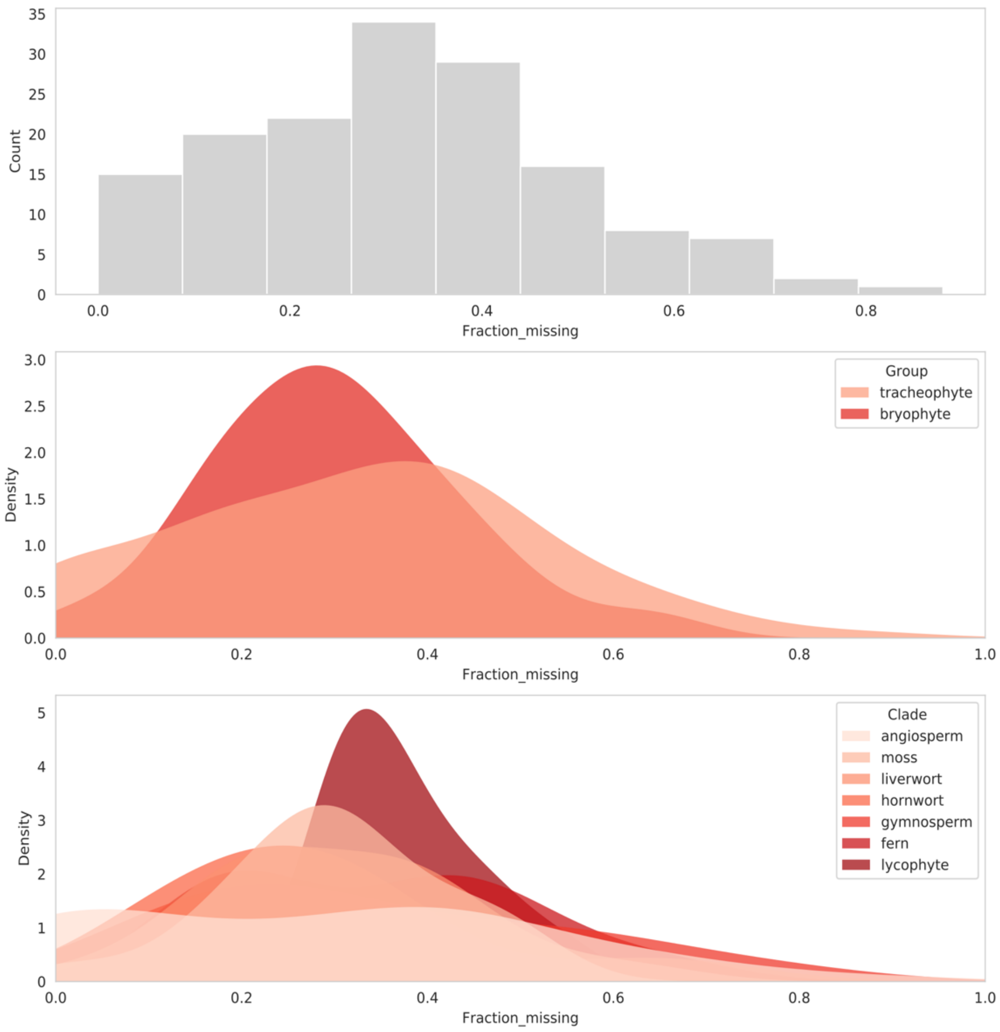


**Supplementary Figure 4. BUSCO completion of genome and transcriptome dataset – 154 species.** Top, histogram of BUSCO completeness. Middle, distribution of BUSCO completeness separated into the two main plant groups. Bryophytes, on average having more complete BUSCOs than tracheophytes. Bottom, distribution of BUSCO completeness separated by lineage. Angiosperms had the most varied completeness values. As the dataset comprised of both genomes and transcriptomes, the underlying variation per lineage may be ascribed to the more incomplete transcriptomes.

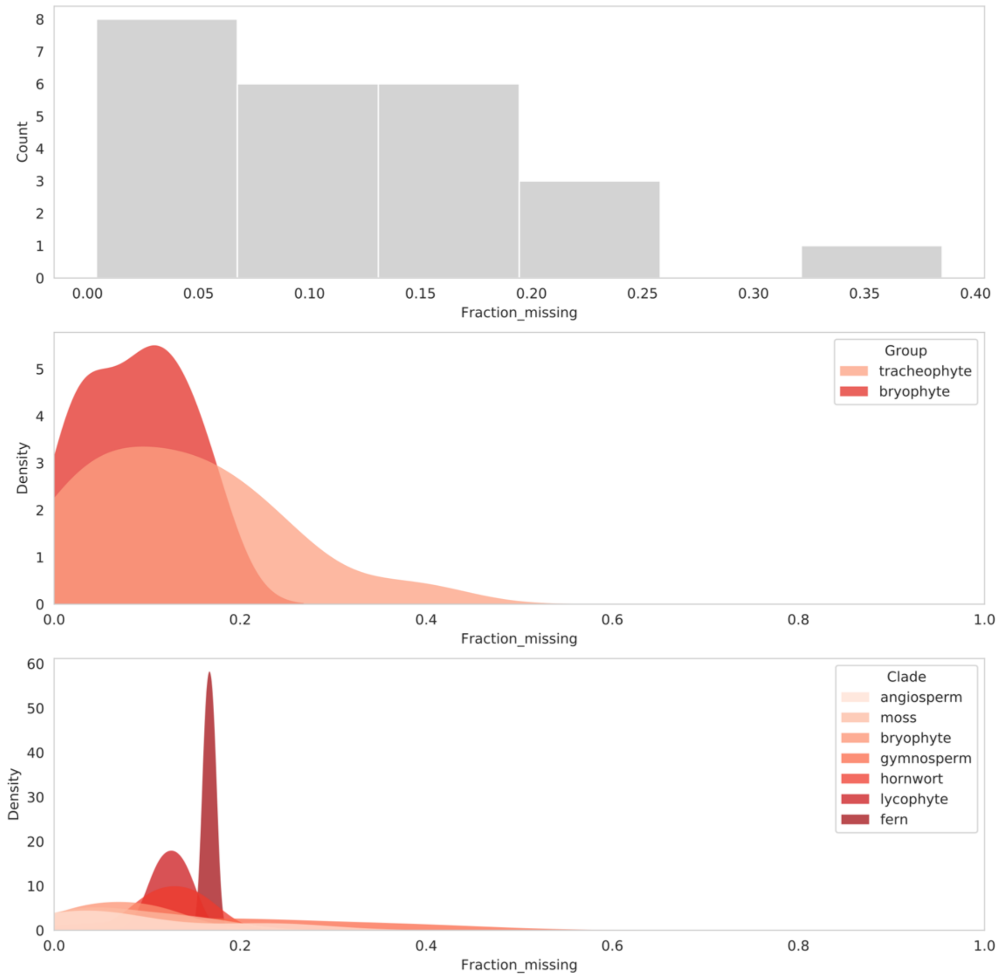


**Supplementary Figure 5. BUSCO completion of genome only dataset – 24 species.** Top, histogram of BUSCO completeness. Middle, distribution of BUSCO completeness separated into the two main plant groups. Bryophytes, on average having more complete BUSCOs than tracheophytes. Bottom, distribution of genome completeness separated by lineage. BUSCO completeness is significantly higher for the genome only dataset.

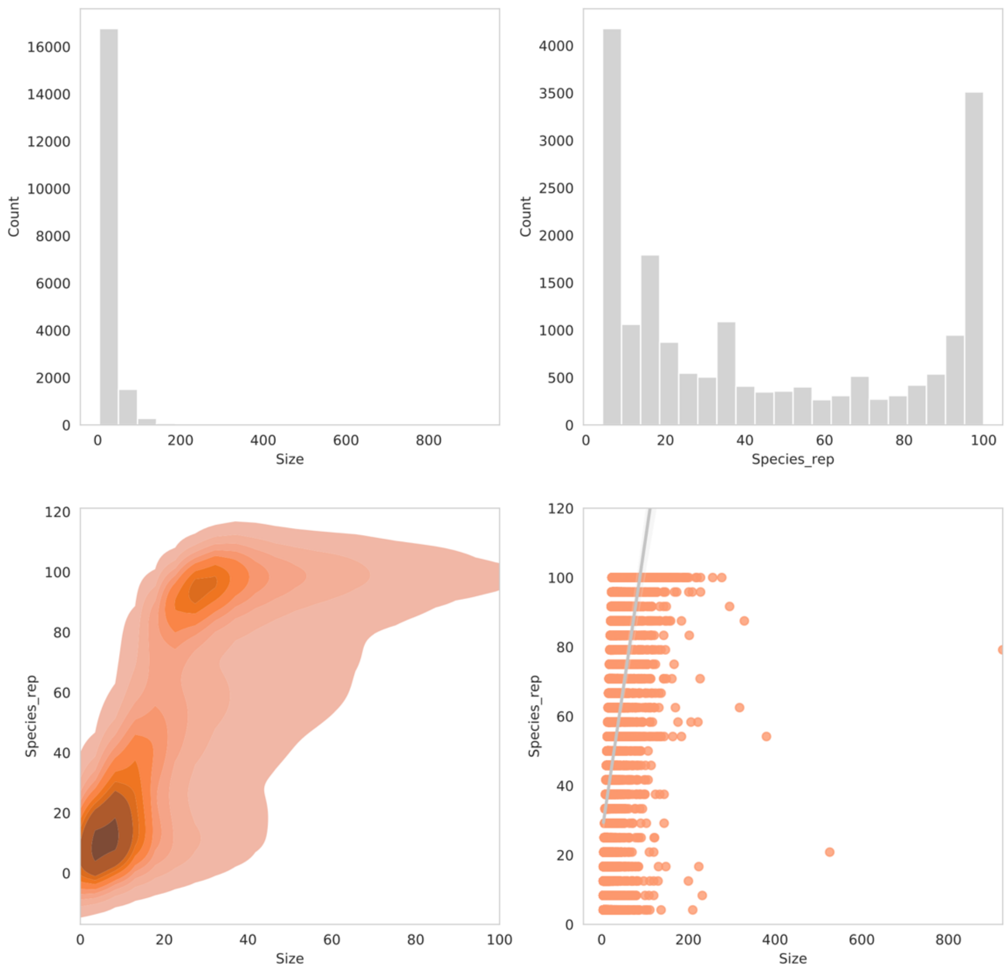


**Supplementary Figure 6. Gene family clusters for genome dataset - 24 species.** Top left, distribution of gene family size. Top right, distribution of gene family species representation. Bottom left, density of gene family species repesentation and size. Bottom right, regression of gene family species representation and size.

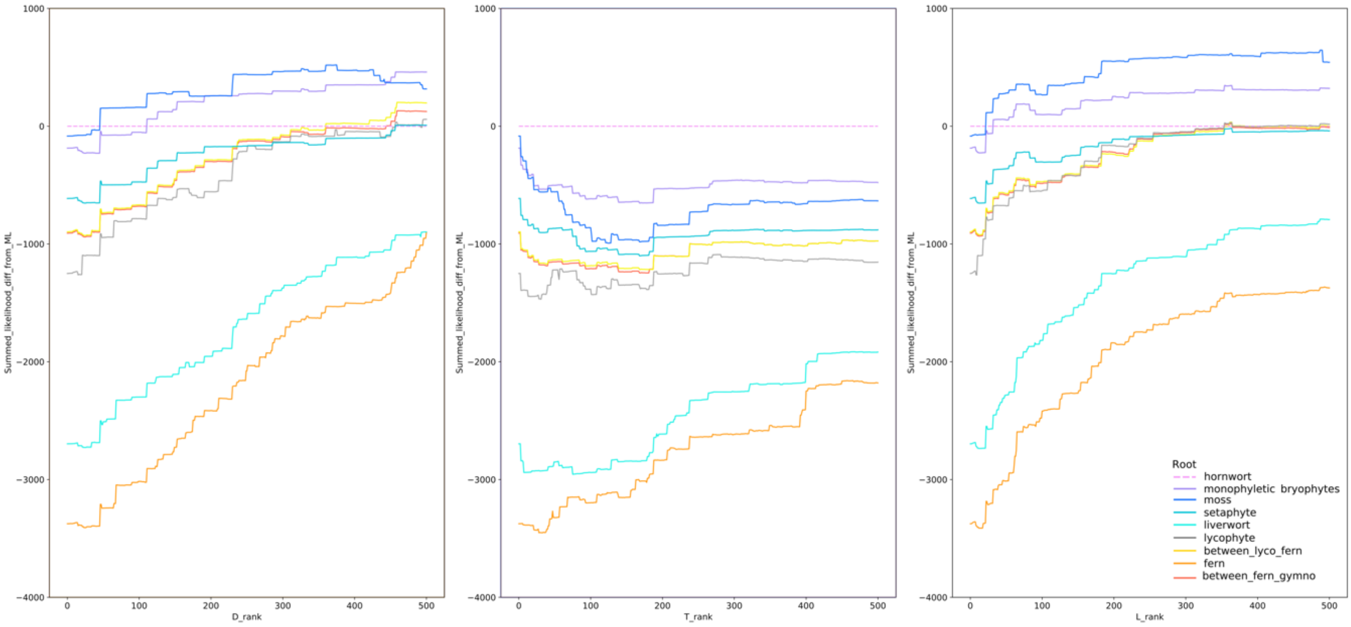


**Supplementary Figure 7. Evaluating the root signal inferred by ALE.** The impact of high duplication, transfer or loss rate families on inferred root order was investigated by iteratively removing top-ranked families and recalculating the summed likelihood on remaining families. While the roots of the credible set remained the top three by summed likelihood, their order varied as high D, T or L-rate families were removed. Removal of high-D and high-L families reduced support for the hornwort root (the ML root on all data), while removal of high-T families increased support for the hornwort root relative to the second- and third-ranked roots. As D, T and L rates were calculated based on the ML root, the preference of high-T families for alternative roots might reflect the difficulty of placing the sparsely-sampled hornworts accurately in individual gene trees.

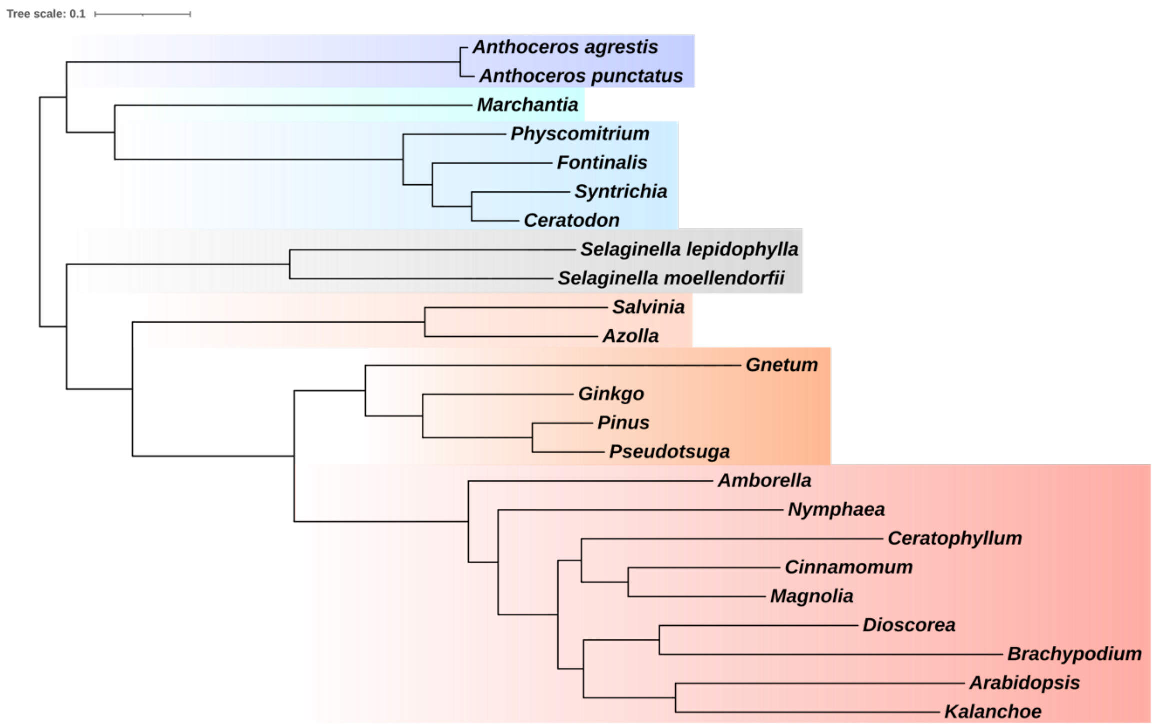


**Supplementary Figure 8. Embryophyte tree rooted with STRIDE.** OrthoFinder 2.0 (Emms & Kelly, 2020) was used to infer orthology groups from the 24 high quality genomes. 23145 orthogroups were inferred. Gene duplication events in each of the orthogroups were assessed using STRIDE (Emms & Kelly, 2017) to root the species tree.

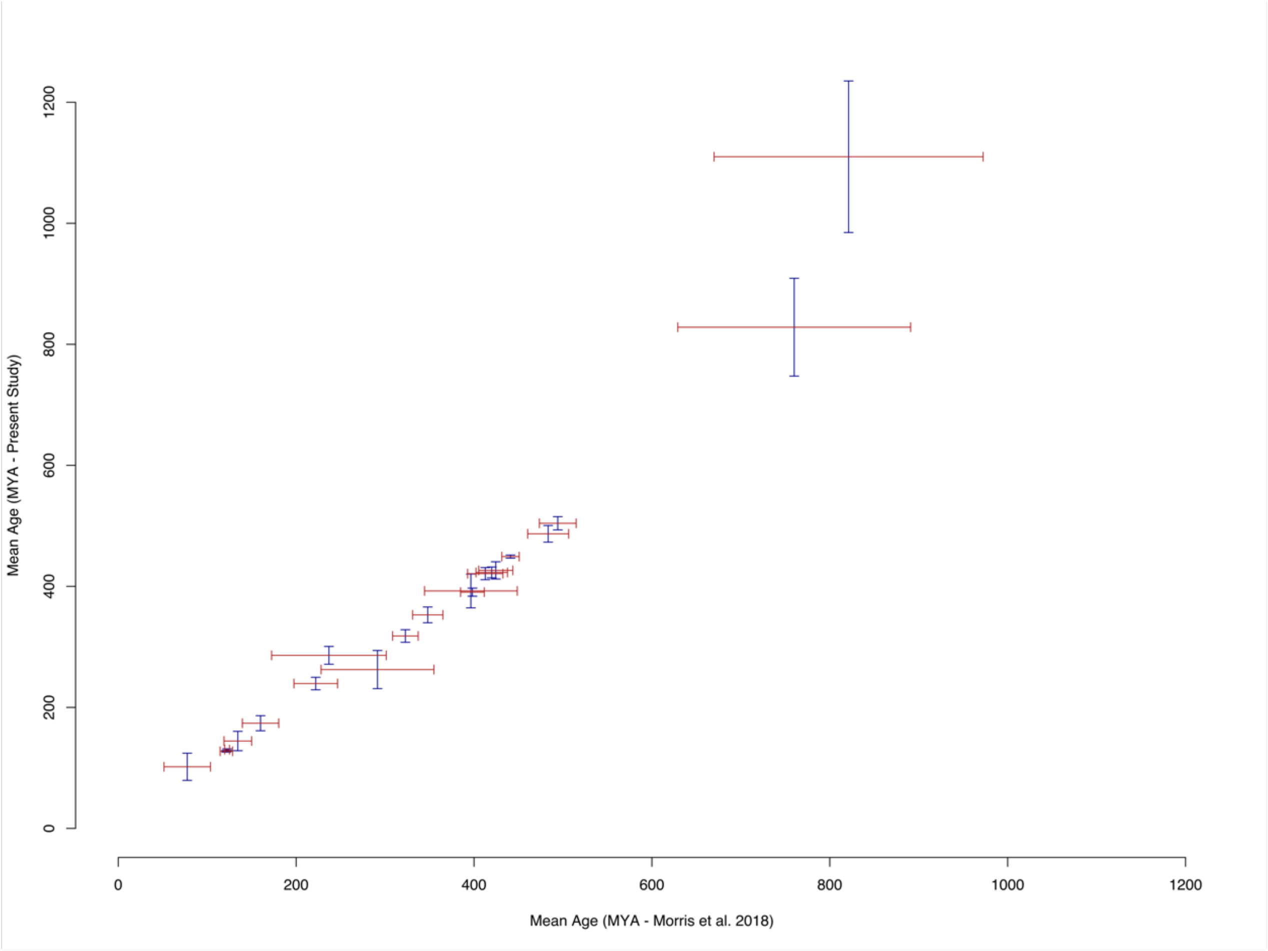


**Supplementary Figure 9**. **A comparison of divergence time estimates between the current study and the closest benchmark, that of Morris et al 2018.** Points on each axis are centred on the mean age, with the 95% highest posterior density represented by the width of the bars.

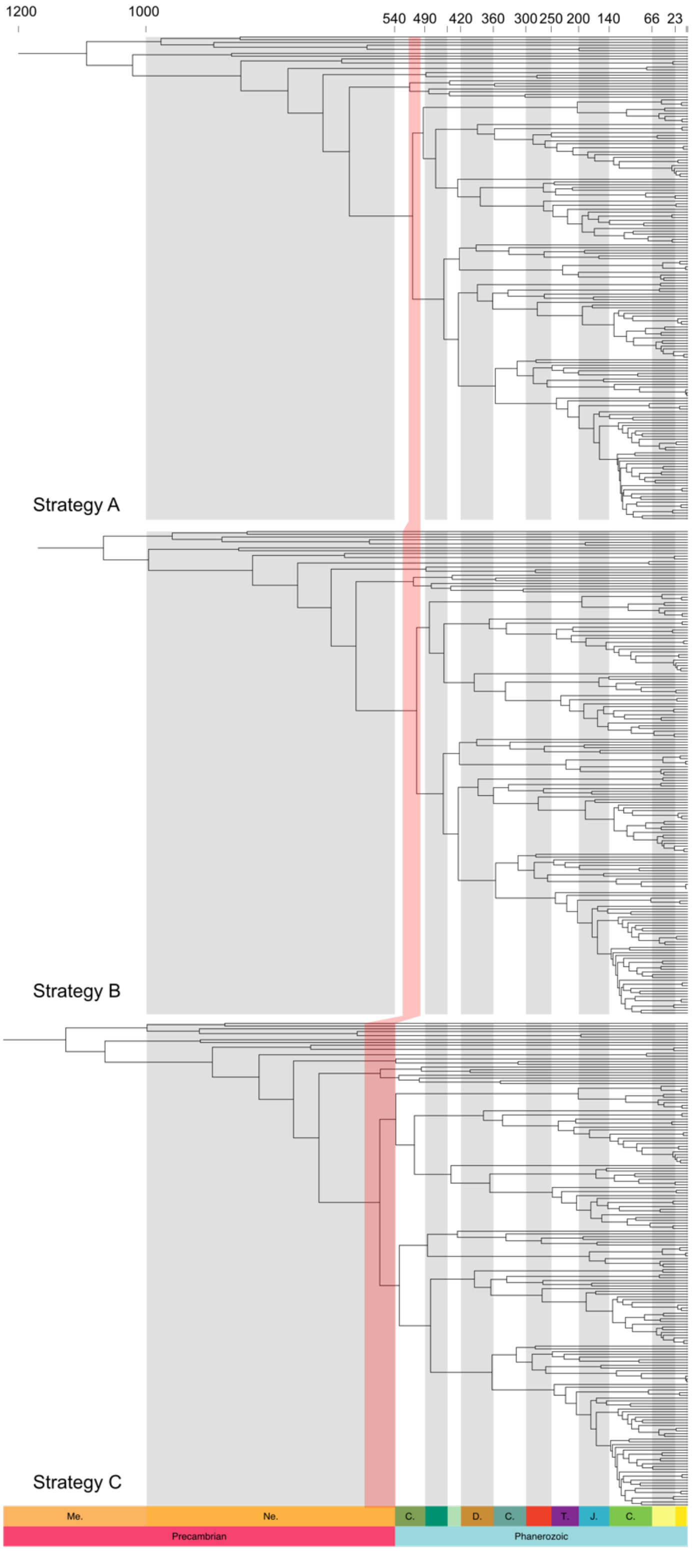


**Supplementary Figure 10. The effect of alternative calibration strategies on the age of crown group embryophytes.** Calibrations were altered by variously relaxing maximum age calibrations on the age of embryophytes (Strategy B) and embryophytes and tracheophytes (Strategy C). The width of the red band across the phylogenies represents the 95% HPD interval.

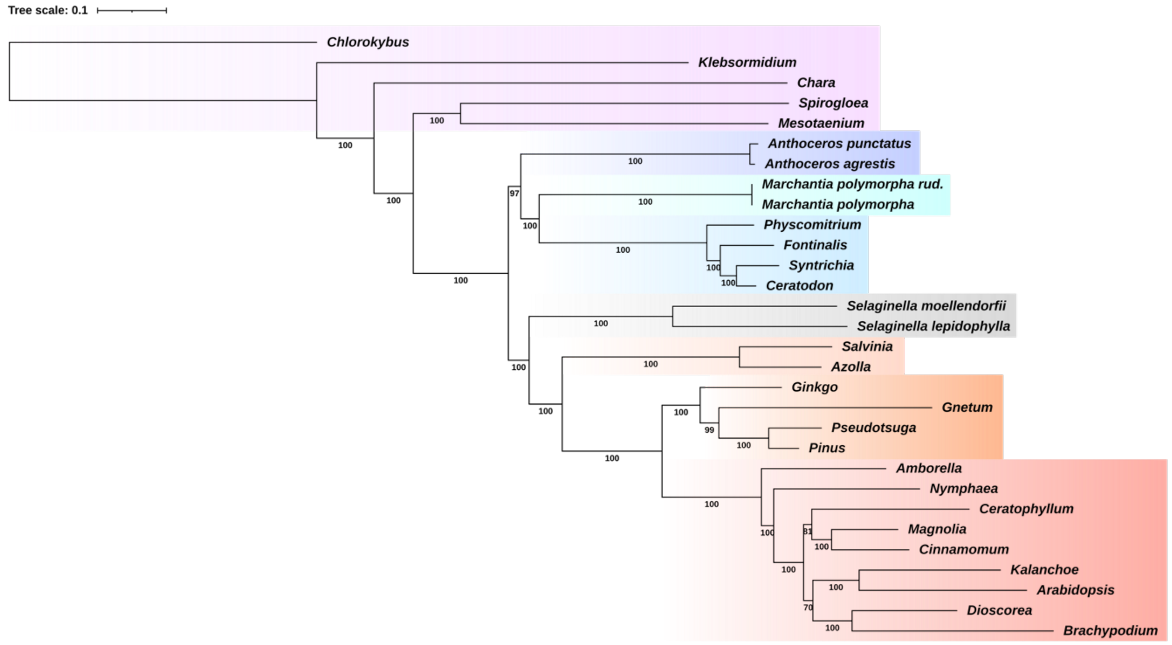


**Supplementary Figure 11. Phylogenetic tree of embryophytes used for ancestral reconstruction**. The maximum likelihood tree was inferred from an alignment comprised of 30 species, 185 single copy orthologs and 71855 sites under the LG+C60+G4+F model in IQ-Tree (Nguyen *et al.,* 2015). Bootstrap support values are placed underneath the corresponding branches. The branch lengths are proportional to the number of average number of substitutions.

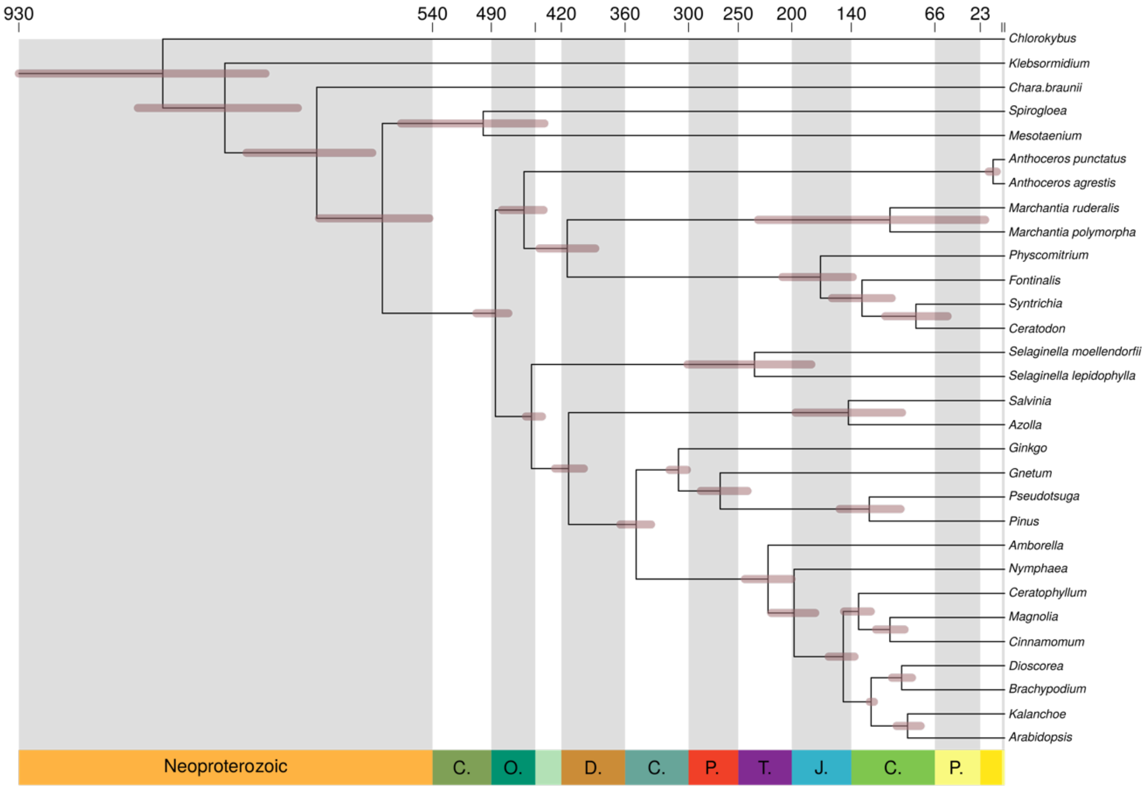


**Supplementary Figure 12. Time scale of embryophyte tree used for ancestral reconstruction**. The amino acid sequence alignment was combined with 18 fossil calibrations subsampled from the list of calibrations from the transcriptomic analysis. The normal approximation method was used in MCMCTree, with branch lengths estimated under the LG+G4 model. Calibrations were modelled as a uniform distribution between a minimum and soft maximum. 95% highest posterior density (HPD) width is shown as bars on node.

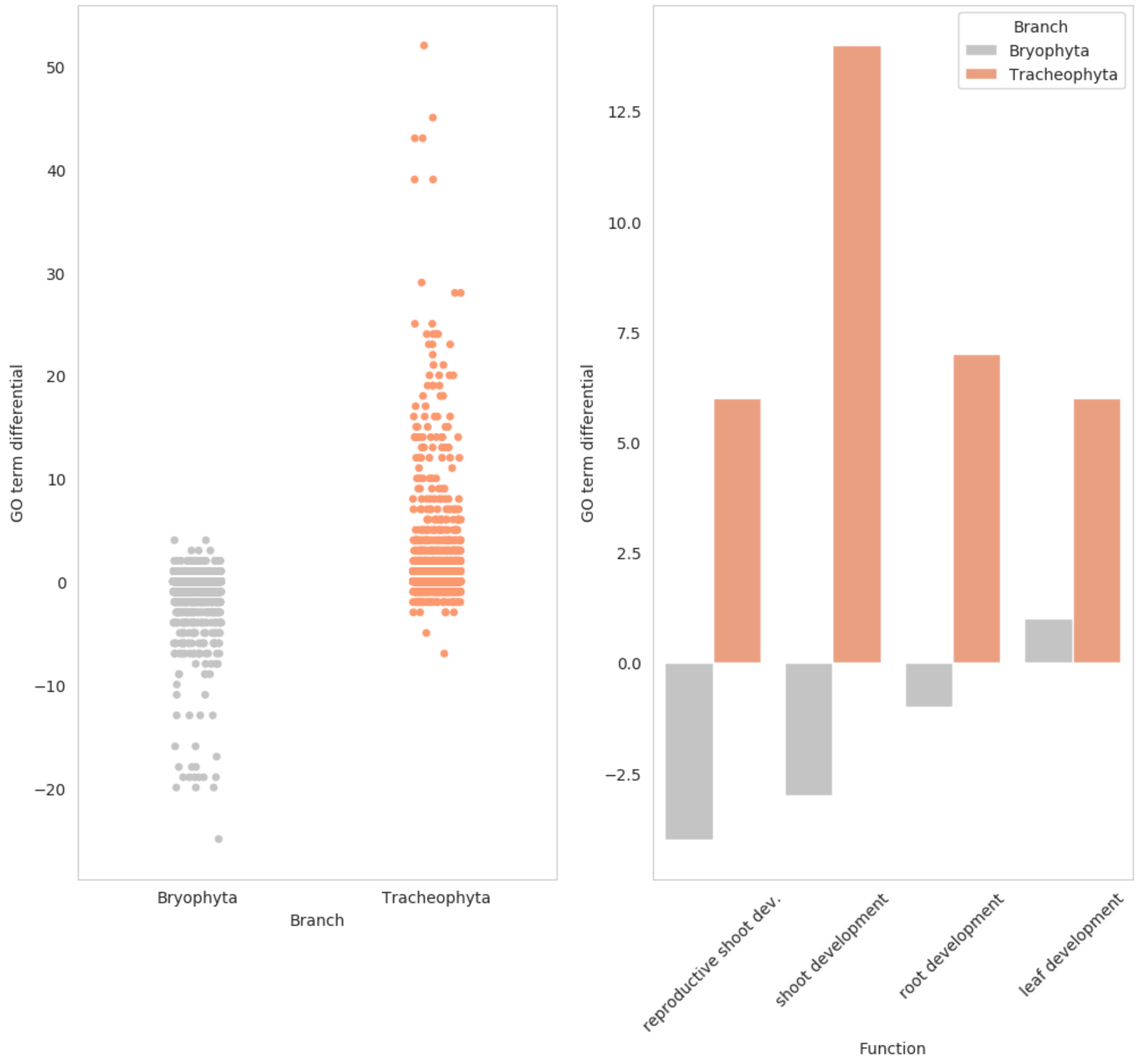


**Supplementary Figure 13**. **Functional annotation of gene family changes between the ancestral embryophyte, bryophytes and tracheophytes.** Left, overall change in GO term frequency between the ancestral embryophyte and the ancestral bryophyte/tracheophyte. GO terms on average become less frequent in bryophytes. Right, change in the frequency of specific GO terms between the ancestral embryophyte and the ancestral bryophyte/tracheophyte. Bryophytes have a reduction in gene families associated with shoot and root development, whilst we see an increase in gene families associated with these GO terms in the tracheophyte ancestor.

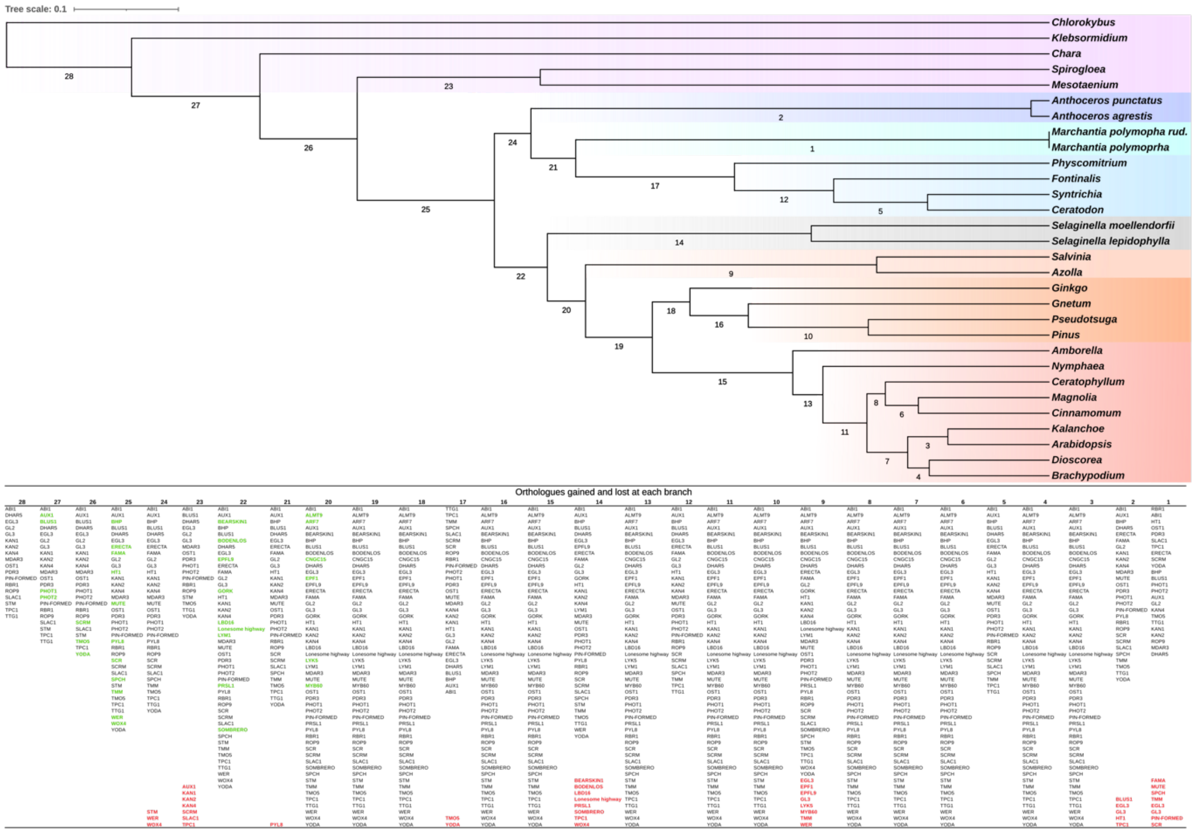


**Supplementary Figure 14**. **Gain and loss of orthologous gene family clusters across the embryophyte tree.** Above, maximum likelihood time calibrated tree, with labels assigned to each branch. Below, the labels correspond to a list of the key orthologous gene family clusters present at each branch. Fifty critical genes for *Arabidopsis,* stomata, vasculature and symbiosis related function/development were used to illustrate the gain and loss of orthologous gene family clusters across the embryophyte tree. Gene families are labelled according to their characterised representatives; for example, the "SLAC1" gene family is the family that contains *Arabidopsis* SLAC1. Gene family clusters that originated on the branch are coloured in green, whilst ones that are lost are coloured red and placed at the bottom of the list. The order of the branches in the table is not consistent with the path of embryophyte evolution i.e. branch 1 is not a child of branch two, but is a child of branch 21, with gain and loss following the phylogenetic tree and not the numerical order of the branch labelling.

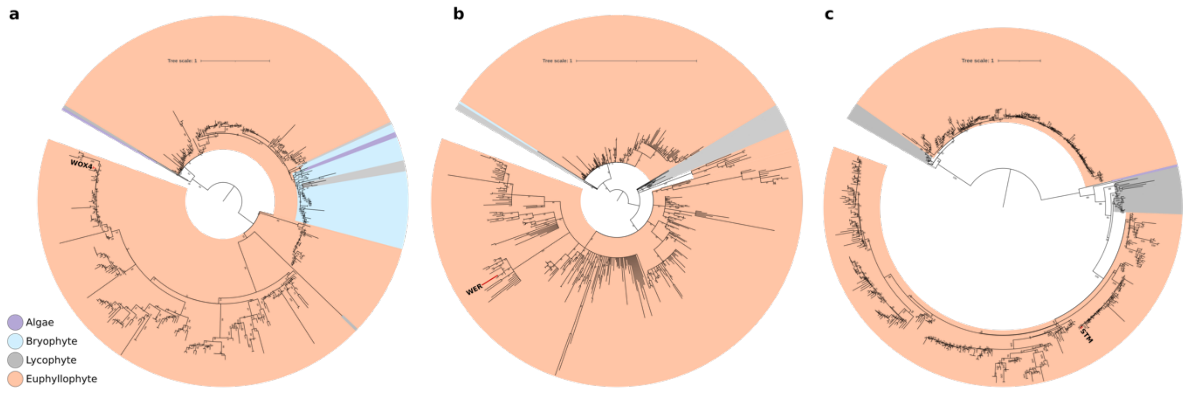


**Supplementary Figure 15**. **Phylogenetic trees of key losses on the bryophyte stem**. Gene trees were constructed from homologous sequence families determined via a BLAST. Gene families were aligned with MAFFT and trimmed with BMGE. Gene trees were inferred in IQ-Tree, under the best fitting phylogenetic model determined by Bayesian Inference Criterion. The trees are rooted in accordance with the species tree displayed in Fig. 1. **a,** Gene tree of WOX4 (*Arabidopsis*) and homologous sequences. The tree suggests that an ancestral WOX gene duplicated prior to the divergence of bryophytes and tracheophytes. Bryophytes have retained one of the WOX genes, but have appeared to have lost the duplicate that went on to become WOX4 in *Arabidopsis,* consistent with the ALE analysis. **b,** Gene tree of WER/WEREWOLF (*Arabidopsis*) and homologous sequences. One bryophyte (*Atrichum)* has retained a WER homolog, thus dating the duplication of the ancestral WER gene to prior to the divergence of bryophytes and tracheophytes. Bryophytes have subsequently lost the duplicate that went on to become WER in *Arabidopsis,* consistent with the ALE analysis. **c,** Gene tree of STM (*Arabidopsis*) and homologous sequences. The ancestral STM gene underwent a duplication, with an extant alga (*Mougeotia*) retaining one of these STM paralogs, thus dating the duplication of the ancestral STM gene to prior the divergence of bryophytes and tracheophytes. Bryophytes have subsequently lost both these paralogs, including the gene that went on to become WER in *Arabidopsis,* consistent with the ALE analysis.
